# PLZF is a new substrate of CRBN with thalidomide and 5-hydroxythalidomide

**DOI:** 10.1101/2020.02.28.969071

**Authors:** Satoshi Yamanaka, Hidetaka Murai, Daisuke Saito, Gembu Abe, Etsuko Tokunaga, Takahiro Iwasaki, Hirotaka Takahashi, Hiroyuki Takeda, Takayuki Suzuki, Norio Shibata, Koji Tamura, Tatsuya Sawasaki

## Abstract

Thalidomide induces cereblon (CRBN)-dependent degradation of proteins. Human cytochrome P450s are thought to provide two monohydroxylated metabolites from thalidomide, and the metabolites are also considered to be involved in thalidomide effects. However, it remains unclear. We report that human PLZF/ZBTB16 is a target protein of CRBN with thalidomide and its derivatives, and that 5-hydroxythalidomide has high potential for degrading PLZF. Using a human transcription factor protein array produced by a wheat cell-free protein synthesis system, PLZF was found to bind to CRBN with thalidomide. PLZF is degraded by the CRL4^CRBN^ complex with thalidomide and its derivatives. Mutagenesis analysis revealed that both 1st and 3rd zinc finger domains conserved in vertebrates are recognized for thalidomide-dependent binding and degradation by CRBN. In chicken limbs, knockdown of Plzf induced skeletal abnormalities, and Plzf was degraded after thalidomide or 5-hydroxythalidomide treatment. Our findings suggest that PLZF is a pivotal substrate involving thalidomide-induced teratogenesis.

## Introduction

In many countries, thalidomide (Fig. 1a) was widely used to treat morning sickness in pregnant women, and caused embryopathies such as limb defects, ear damage, and congenital heart diseases^1–3^. Handa’s group in Japan found that thalidomide binds to the cereblon (CRBN) protein within the CRL4 E3 ubiquitin ligase complex and that CRBN is a key molecule for thalidomide-induced teratogenesis^4^. Recently many studies have reported that CRBN, by interacting with thalidomide or its derivatives, lenalidomide, pomalidomide (Fig. 1a), and CC-885, changes its binding specificity to proteins, then induces ubiquitination and degradation of binding proteins such as Ikaros (IKZF1)^5,6^, casein kinase I^7^, GSPT1^8^, and SALL4^9,10^. In humans, mutations of SALL4 are found in Duane-radial ray syndrome (DRRS, Okihiro syndrome), which has phenotypic features such as limb deformities^11^. In *Sall4*-conditional knockout (SALL4-CKO) mice, hindlimb defects are also reported^12^. These findings strongly suggest that SALL4 is partially involved in teratogenesis. However, it was reported that forelimb of SALL4-CKO mice were not abnormality and posterior of hindlimb were formed in the SALL4-CKO mice. Furthermore, although thalidomide embryopathy occurs in chicken and zebrafish^4^, the sequence of thalidomide-binding sites in these Sall4s is quite different between human and these animals^9^, suggesting the possibility of other target proteins for teratogenesis.

**Figure 1.**
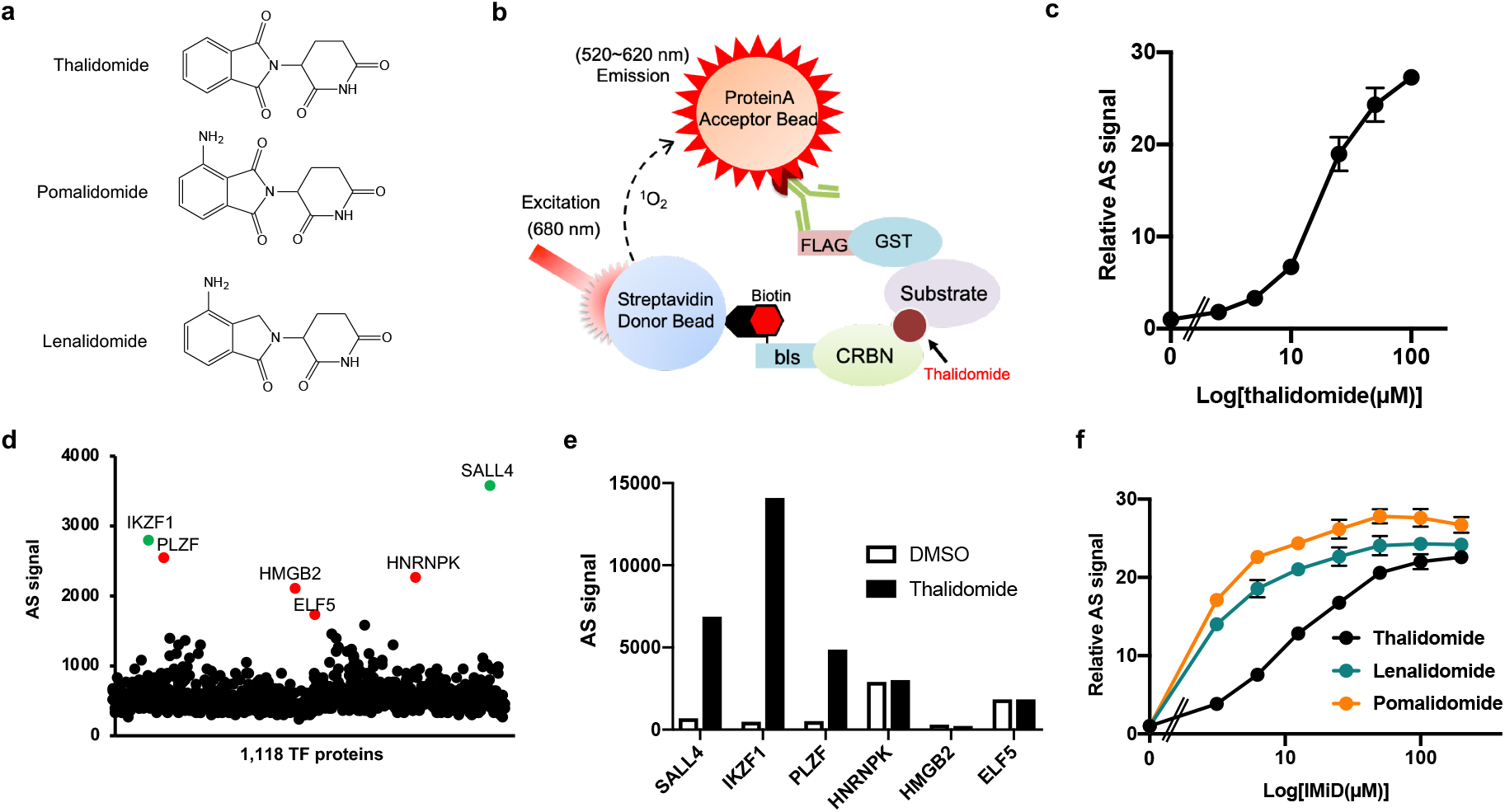
Identification of thalidomide-dependent interactors of CRBN using a cell-free based human TF protein array. **a**, Chemical structures of thalidomide, pomalidomide and lenalidomide. **b**, Schematic diagram of the thalidomide-dependent *in vitro* binding; assay between CRBN and substrates using AlphaScreen technology. **c**, Detection of luminescent signals of thalidomide-dependent interactions between bls-CRBN and FLAG-GST-IKZF1. Dose-dependent signals (DMSO, 2.5, 5, 10, 25, 50, or 100 μM thalidomide) was analysed with an *in vitro* binding assay using AlphaScreen technology. **d**, Results of *in vitro* high-throughput screening, targeting 1,118 human transcription factors. Green and red spots denote known neosubstrates and candidate clones, respectively. **e**, Confirmation of thalidomide-dependency on six hit proteins using an *in vitro* binding assay. Interaction between bls-CRBN and FLAG-GST-protein in the presence of DMSO or 50 μM thalidomide was detected using AlphaScreen technology. **f**, *In vitro* binding assay for thalidomide, pomalidomide, and lenalidomide. Interaction between bls-CRBN and FLAG-GST-PLZF in the presence of DMSO, (3.125, 6.25, 12.5, 25, 50, 100, or 200 μM) thalidomide, pomalidomide or lenalidomide was analysed using AlphaScreen technology. All relative AS (AlphaScreen) signals were expressed as relative luminescent signal with luminescent signal of DMSO as one, and error bars mean ± standard deviation (n=3).

Promyelocytic leukaemia zinc finger (PLZF), also known as ZBTB16 or ZFP145, is a transcription factor (TF) that has nine C2H2-type zinc finger domains (ZNFs)^13^, and is involved in a broad range of developmental and biological processes, such as haematopoiesis, limb skeletal formation, spermatogenesis, and immune regulation^13–15^. Loss of PLZF function in both human patients and mouse mutants indicates limb defects, which mice phenotype was elongation defect of zeugopod and thumb^14–16^. A recent study has indicated that the 6th and 7th ZNFs in PLZF were not targets of CRBN^17^ However, it remains unknown whether full-length PLZF protein is a target for CRBN with thalidomide.

Thalidomide is metabolized into 5-hydroxythalidomide and 5’-hydroxythalidomide by several types of human cytochrome P450 (CYP)^18–20^, which are oxidized in phthalimido and glutarimide rings, respectively (Fig. 4a). Because CRBN mainly recognizes the glutarimide ring in thalidomide^21,22^, 5’-hydroxythalidomide has no functions in either CRBN binding or teratogenesis^23,24^. Recently, mice having humanized-CYP3A that could potentially provide 5-hydroxythalidomide were shown to undergo teratogenesis after thalidomide treatment^25^, suggesting that 5-hydroxythalidomide functions like thalidomide. However, there is no evidence to suggest whether 5-hydroxythalidomide induces interactions between CRBN and substrate proteins.

We aimed to identify the protein that binds thalidomide-dependently to CRBN. For this, we constructed a human TF protein array (HuTFPA) consisting of 1,118 human recombinant proteins including mainly TFs and zinc finger proteins (Supplementary Table 1), which was produced using a wheat cell-free protein production system. Biochemical screening based on an interaction between TF and CRBN with thalidomide using the AlphaScreen system, identified PLZF as an interactor of CRBN with thalidomide. PLZF was shown to be a novel substrate of the CRL4^CRBN^ E3 ubiquitin ligase complex with thalidomide, its derivatives, and 5-hydroxythalidomide. Amino acid sequences of PLZF were very similar among vertebrates. In the chick embryo, knockdown of Plzf/Zbtb16 induced abnormal limb development. More importantly, Plzf protein was decreased in the limb buds during thalidomide-induced teratogenesis, whereas expression level of Sall4 protein was not changed.

## Results

### Screening for thalidomide-dependent substrates of CRBN using a human transcription factor protein array

Many TFs function as master regulators during development and differentiation of embryos. In addition, many substrates of CRBN with thalidomide were ZNF-type TFs such as IKZF1^5,6^, IKZF3^5,6^, and SALL4^9,10^. We aimed to identify new substrates of CRBN with thalidomide from human TFs. Based on a wheat cell-free protein synthesis system^26^, we had previously developed a technology for the construction of a protein array that synthesizes an individual protein in each well of a 96- or 384-well plate, and have published many reports regarding the identification of substrate proteins of protein kinase^27^ and E3 ligase^28,29^ protein arrays. The combination of our protein array and AlphaScreen technology provides several advantageous features for the screening of protein-protein interactions: it is 1) used directly without protein purification, 2) highly sensitive, and 3) a high-throughput system. The human TF protein array (HuTFPA) consisting of mainly human TFs (Supplementary Table 1), synthesized as N-terminal FLAG-GST fusions, produced by a wheat cell-free system. CRBN was synthesized as an N-terminal single biotin-labelled form, using the same system. The principle of detecting this biochemical interaction is shown in Fig. 1b. Using this cell-free system, an interaction between FLAG-GST-IKZF1 and biotinylated CRBN was thalidomide dose-dependently detected using the AlphaScreen method (Fig. 1c). As shown in the flowchart (Supplementary Fig. 1a), we screened the substrate human TFs of CRBN with thalidomide (50 μM) on the HuTFPA, identifying six TF proteins as being CRBN binding proteins in the presence of thalidomide (Fig. 1d). In contrast, several known substrates such as IKZF3 and CK1α, were not detected because these substrates were not included in the HuTFPA (Supplementary Table 1).

To investigate the thalidomide dependency of these proteins for CRBN binding, the biochemical assay was carried out with and without thalidomide. Three human TFs, IKZF1, SALL4, and PLZF, indicated characteristics of thalidomide-dependent binding to CRBN (Fig. 1e), whereas HNRNPK, HMGB2, and ELF5 bound to CRBN without thalidomide. Furthermore, in the presence of thalidomide, these did not bind to mutant CRBN-YW/AA^3^ that is unable to bind to thalidomide (Supplementary Fig. 1b). Using recombinant proteins, *in vitro* pull-down assay confirmed that PLZF binds to wild-type (WT) CRBN in the presence of thalidomide (Supplementary Fig. 1c), as do IKZF1 and SALL4, whereas this binding was not observed under the condition of using mutant CRBN-YW/AA (MT) and no addition of thalidomide (–). These results indicate that PLZF is an interactor of CRBN with thalidomide.

### PLZF binds to CRBN with thalidomide, pomalidomide, and lenalidomide

Through the screening above, PLZF was identified as a candidate substrate for CRBN with thalidomide. PLZF is classified as being part of the ZBTB (zinc finger and bric à brac, tramtrack, and broad) protein family^13^. Since the other ZBTB proteins are included in the HuTFPA, these were analysed for interactions with or without thalidomide. However, the ZBTB proteins did not bind to CRBN with thalidomide (Supplementary Fig. 1d), suggesting that CRBN with thalidomide recognizes a specific region(s) in PLZF, but not one common to the ZBTB family.

During the last two decades, two thalidomide derivatives, lenalidomide and pomalidomide (Fig. 1a), have been developed for multiple myeloma (MM) or as immunomodulatory drugs (IMiDs)^30^. Recent studies have reported that some proteins have different preferences between thalidomide and its derivatives for binding to CRBN^7,9^. For an example, a CK1α– CRBN interaction is enabled by lenalidomide, but not thalidomide or pomalidomide^7^. We therefore investigated the biochemical characteristics of interactions between PLZF and CRBN with thalidomide and its two derivatives. As a result, thalidomide, pomalidomide and lenalidomide induced PLZF–CRBN interaction, and it is showed that the biochemical binding potency is pomalidomide>lenalidomide>thalidomide (Fig. 1f). In a previous report^9^, it was showed that thalidomide and its two derivatives induced degradation of SALL4. In addition, in vitro binding assay using the AlphaScreen method confirmed that SALL4’s binding potency is pomalidomide>thalidomide>lenalidomide (Supplementary Fig. 1e). Because the binding potency of PLZF-thalidomide is similar to that of SALL4-lenalidomide, it was predicted that PLZF and SALL4 are pan-substrates on thalidomide and its two derivatives.

### PLZF is degraded by the CRL4^CRBN^ complex with thalidomide and its derivatives

CRBN consists of a CRL4^CRBN^ E3 ubiquitin ligase complex which includes DDB1, RBX1, and CUL4^4^. Therefore, the PLZF–CRBN interaction with thalidomide and its two derivatives is expected to lead to degradation of PLZF. For a cell-based immunoblotting assay, we used the AGIA-tag system because it is a highly-sensitive tag based on a rabbit monoclonal antibody^31^. To investigate the stability of PLZF or SALL4, AGIA-tagged PLZF or SALL4 was transfected into HEK293T cells with thalidomide, pomalidomide, and lenalidomide. These compounds all decreased the stability of PLZF and SALL4 (Supplementary Fig. 2a). In addition, to demonstrate whether PLZF is also pan-substrate, such as SALL4, endogenous PLZF or SALL4 protein levels were examined in HuH7 or HEK293T cells treated with thalidomide and its two derivatives. Immunoblot analyses revealed that endogenous PLZF were destabilized by all of them in both cell lines (Supplementary Fig. 2b and c), indicating that PLZF is also pan-substrate on thalidomide and its two derivatives. To reduce experimental complexity, therefore, we focused thalidomide and lenalidomide in further analyses. A remarkable decrease in PLZF was observed 6 hours after lenalidomide treatment (Supplementary Fig. 2d). This time course analysis suggests that the reduction in PLZF was late as compared to other CRBN substrates because degradations of IKZF1, IKZF3, and SALL4 were observed only 3 hours after the treatment^5,6,9^. The lenalidomide-dependent destabilisation of PLZF was completely inhibited by the proteasome inhibitor MG132 and the NEDD8 inhibitor MLN4924 (Fig. 2a), suggesting that PLZF is ubiquitinated by the CRL4 complex for degradation by the proteasome.

**Figure 2.**
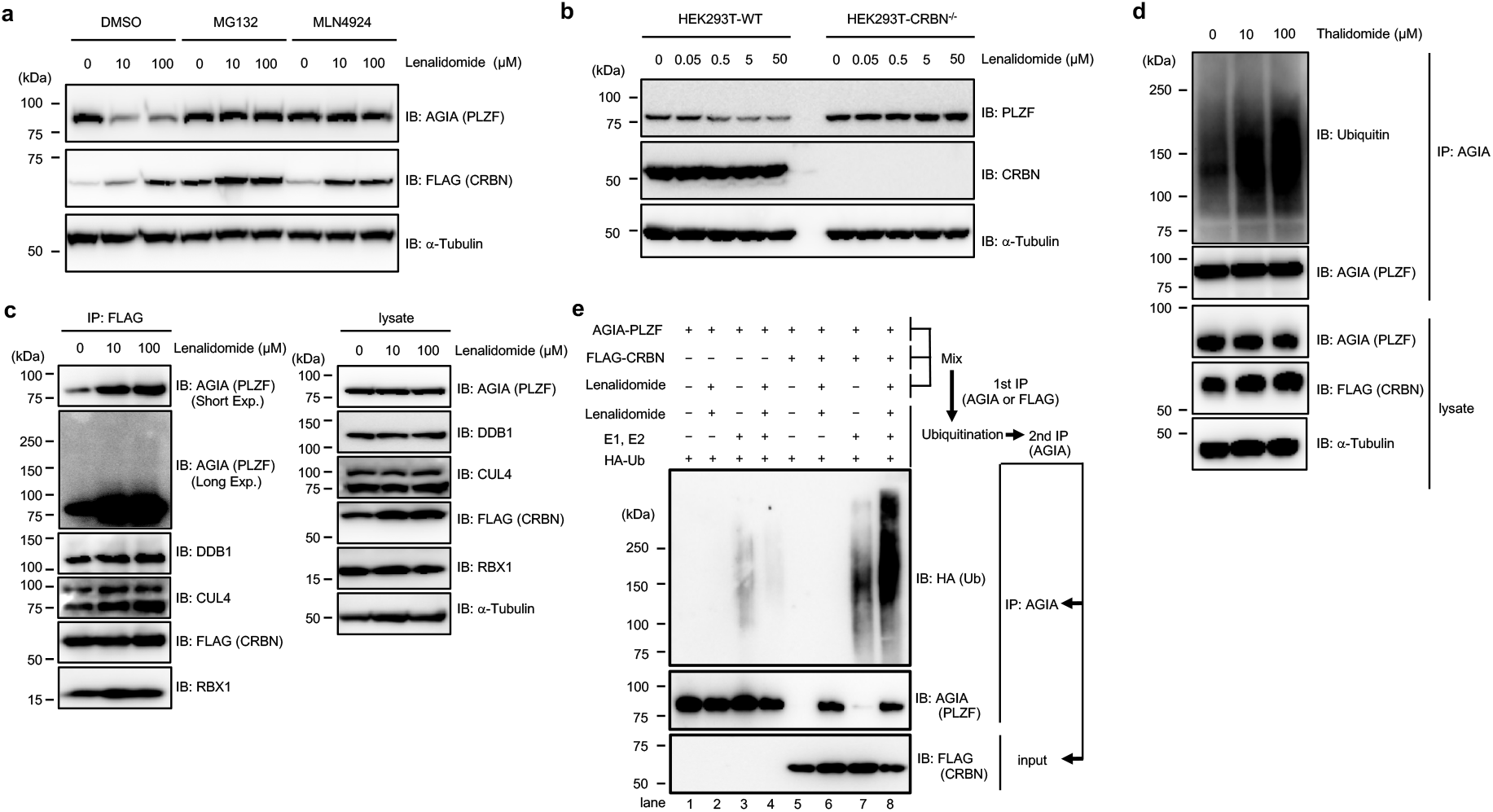
PLZF is a substrate of CRL4^CRBN^ with thalidomide and lenalidomide for E3 ubiquitin ligase. **a**, Immunoblot analysis of AGIA-PLZF protein levels in AGIA-PLZF and FLAG-CRBN expressing HEK293T cells treated with DMSO or lenalidomide in the presence of DMSO, MG132, or MLN4924 for 9 h. **b**, Immunoblot analysis of endogenous PLZF protein levels in HEK293T cells or CRBN^-/-^ HEK293T cells treated with DMSO or lenalidomide for 24 h. **c**, Immunoprecipitation of FLAG-CRBN in FLAG-CRBN and AGIA-PLZF expressing HEK293T cells treated with DMSO or lenalidomide in the presence of DMSO or MG132 for 8 h. Components of CRL^FLAG-CRBN^ and AGIA-PLZF were detected using each specific antibody, as indicated. **d**, Ubiquitination of AGIA-PLZF in AGIA-PLZF and FLAG-CRBN expressing CRBN^-/-^ HEK293T cells treated with DMSO or thalidomide in the presence of DMSO or MG132 for 10 h. AGIA-PLZF was immunoprecipitated using anti-AGIA antibody and the polyubiquitin chain on AGIA-PLZF was analysed by immunoblot. **e**, *In vitro* binding and ubiquitination assay of AGIA-PLZF. Empty vector, AGIA-PLZF, or FLAG-CRBN expressing HEK293T cells were lysed and the lysates were mixed. The first immunoprecipitation with anti-AGIA or anti-FLAG antibodies was performed in the presence of DMSO or 200 μM lenalidomide. The purified AGIA-PLZF or CRL4^FLAG-CRBN^ complex, including AGIA-PLZF and FLAG-CRBN, was incubated with recombinant E1, E2, and HA-ubiquitin in the presence of DMSO or 200 μM lenalidomide, and the second immunoprecipitation was performed using anti-AGIA antibody. Ubiquitination of PLZF was analysed by immunoblot.

To investigate whether the destabilisation of PLZF is dependent on CRBN, we made CRBN-deficient HEK293T cells using CRISPR/Cas9. In the presence of lenalidomide, degradation of endogenous PLZF was observed in normal HEK293T cells, whereas endogenous PLZF was not degraded in CRBN-deficient cells (Fig. 2b). Endogenous PLZF protein was also reduced by lenalidomide in HuH7 and THP-1 cells (Supplementary Fig. 3b and c). Expression of endogenous *PLZF* mRNA in these cells was unaffected by lenalidomide treatment (Supplementary Fig. 3a-c). In addition, the FLAG-CRBN mutant (YW/AA) did not degrade AGIA-PLZF in the presence of lenalidomide (Supplementary Fig. 2e), suggesting the CRBN-dependent degradation of PLZF.

We next investigated whether PLZF is recruited to the CRL4^CRBN^ complex and ubiquitinated. By immunoprecipitation of FLAG-CRBN using an anti-FLAG antibody, the CRL4^CRBN^ complex consisting of DDB1, RBX1, and CUL4 was pulled-down. At the same time AGIA-tagged PLZF was also strongly associated with the complex after thalidomide or lenalidomide treatment (Fig. 2c and Supplementary Fig. 4a), indicating that PLZF is thalidomide- or its two derivatives-dependently included in the CRL4^CRBN^ complex. In addition, to analyse the ubiquitination of PLZF, AGIA-PLZF was transfected with FLAG-CRBN and immunoprecipitated for immunoblotting. The smear band resulting from the immunoprecipitation of AGIA-PLZF was increased by supplementation with thalidomide or lenalidomide, suggesting that PLZF ubiquitination was induced by thalidomide and its two derivatives treatment (Fig. 2d and Supplementary Fig. 4b). Next, to investigate the in vitro ubiquitination of PLZF, CRL4^FLAG-CRBN^ and AGIA-PLZF were coimmunoprecipitated by anti-FLAG antibody in the presence or absence of lenalidomide (lane 5-8 in Fig. 2e). As negative control, empty vector or AGIA-PLZF expressing HEK293T cells were lysed and mixed, and the lysates were immunoprecipitated by anti-AGIA antibody to demonstrate the ubiquitination of PLZF caused by CRL4^FLAG-CRBN^ (lane 1-4 in Fig. 2e). When coimmunoprecipitating PLZF and CRBN plus exogenous E1 and E2 enzymes, PLZF was ubiquitinated in the presence of lenalidomide (lane 8 in Fig. 2e), but not in its absence (lane 6 in Fig.2e), indicating that CRL4^FLAG-CRBN^ ubiquitinates PLZF *in vitro*. Taken together, these results show that PLZF is a target of the CRL4^CRBN^ complex for proteasome degradation.

Next, because it was reported that PLZF play important role in immune responses^13^, we investigated whether degradation of PLZF is also caused in lymphoma cell lines. Immunoblot analyses showed that IMiD treatment induced protein degradation of PLZF in ABC-DLBCL (TK), GCB-DLBCL (BJAB and HT), Adult T-cell lymphoma/ATL (MT-4), and Burkitt’s lymphoma (Raji) cell lines (Supplementary Fig. 5a and b). In GCB-DLBCL (SU-DHL-4) cells, IMiD treatment scarcely induced protein degradation of PLZF but protein expression of CRBN in SU-DHL-4 cells was very weak (Supplementary Fig. 5a). These results strongly suggest that PLZF degradation is caused by IMiD treatment in various cells.

### Both ZNF1 and ZNF3 domains in PLZF are recognized for its thalidomide-dependent interaction with CRBN

We next attempted to determine the thalidomide-dependent CRBN interaction region within PLZF. PLZF has a single BTB domain and nine ZNFs^13^ (Fig. 3a). A single ZNF domain present in IKZF1^17^ and SALL4^9,10^ is recognized by CRBN with thalidomide. Expectedly, the BTB domain alone in PLZF did not induce CRBN binding (Fig. 3b). We thus constructed a total of five clones lacking different ZNFs by N-terminal FLAG-GST-fusions, then measured the interaction signals between each clone and CRBN with thalidomide. As a result, a ZNF1-5 clone indicated sufficient binding ability to CRBN with thalidomide (Fig. 3b). To refine the key domains on the native form of PLZF, each of the five ZNFs were individually swapped with ZNF7 (Fig. 3c), as this had no effect on binding^17^. These binding assays showed that the ZNF1 and ZNF3 domains were most important for binding (Fig. 3d). In previous reports, a glycine residue in a ZNF domain of SALL4 and ZFP91 proteins was shown to be a key amino acid for binding to CRBN with thalidomide^9,10,32^. We thus made mutant clones (Gly to Ala shown in Fig. 3e) having a single or double substitution in ZNF1 and ZNF3, which were then analysed in biochemical (Fig. 3f) and cell-based (Fig. 3g) assays. Surprisingly, the double mutation completely lost the ability for binding and degradation, whereas binding ability of both single mutants significantly low but retained degradation ability (Fig. 3f and g). Taken together, these results indicate that both ZNF1 and ZNF3 domains in PLZF are recognized for binding and degradation by CRBN with thalidomide.

**Figure 3.**
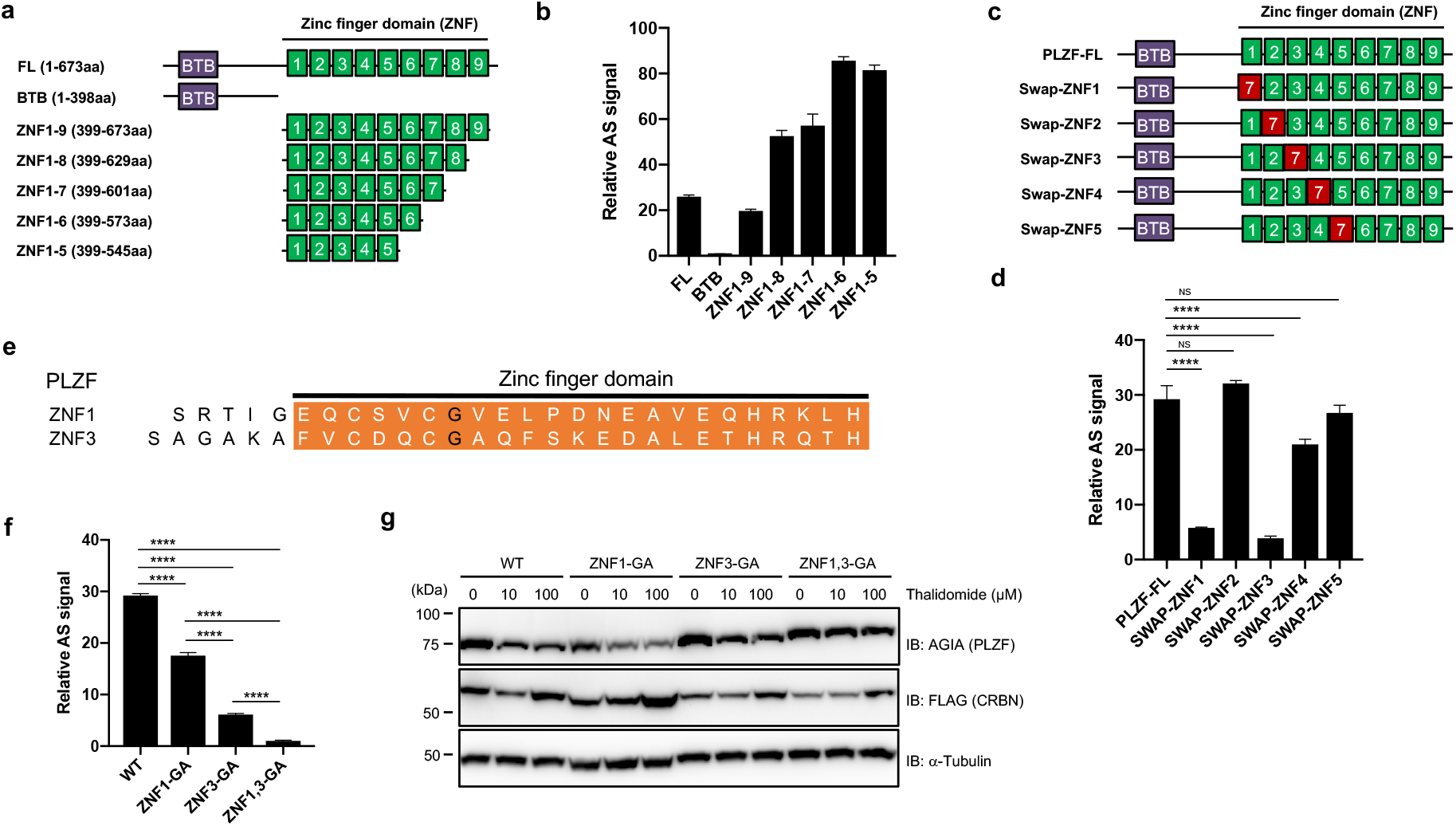
Interaction regions in PLZF for binding to CRBN with thalidomide. **a**, Schematic diagram of PLZF and truncated PLZFs. **b**, *In vitro* binding assay using truncated PLZF. Thalidomide-dependent interaction between bls-CRBN and FLAG-GST-PLZF-full length (FL) or truncated FLAG-GST-PLZF was analysed in the presence of DMSO or 50 μM thalidomide using AlphaScreen technology. **c**, Schematic diagram of swapped PLZF mutants. **d**, *In vitro* binding assay using swapped PLZF mutants was performed using the same procedure as in Figure 3b. **e**, Amino acid sequences of ZNF1 and ZNF3 in PLZF. **f**, *In vitro* binding assay using point mutants of PLZF was performed using the same procedure as in Figure 3b. **g**, Immunoblot analysis of AGIA-PLZF protein levels in FLAG-CRBN and PLZF-WT, PLZF-ZNF1-GA, PLZF-ZNF3-GA, or PLZF-ZNF1,3-GA expressing CRBN^-/-^ HEK293T cells treated with DMSO or thalidomide for 16 h. All relative AS (AlphaScreen) signals were expressed as relative luminescent signal with luminescent signal of DMSO as one. Error bars mean ± standard deviation (n = 3) and *P* values were calculated by one-way ANOVA with Tukey’s post-hoc test (NS = Not Significant, and \*\*\*\**P* < 0.0001).

### 5-hydroxythalidomide induces degradation of PLZF and SALL4, but not of IKZF1

5-hydroxythalidomide (Fig. 4a) is produced as a primary metabolite of thalidomide by several types of human CYP^18–20^. However, the function of 5-hydroxythalidomide remains unclear. We thus investigated whether it induces a CRBN–protein interaction. Surprisingly, 5-hydroxythalidomide induces CRBN–PLZF and CRBN–SALL4 interactions (Fig. 4b), whereas it did not induce an interaction between CRBN and IKZF1. Furthermore, 5-hydroxythalidomide induces the degradation of PLZF and SALL4 in cells, but not of IKZF1 (Fig. 4c). Endogenous PLZF and SALL4 were also degraded in HuH7 and THP-1 cells, respectively, after treatment with 5-hydroxythalidomide (Fig. 4d and e). Endogenous IKZF1 showed no change in the presence of 5-hydroxythalidomide (Fig. 4e), even though PLZF was degraded under the same conditions. Notably, dose-dependency between thalidomide and 5-hydroxythalidomide in the biochemical CRBN–SALL4 interaction was almost identical (middle panel in Fig. 4b). In addition, degradations of PLZF and SALL4 by 5-hydroxythalidomide in cells occurred at almost the same levels as those in the thalidomide treatment. Taken together with the function of human CYP3A^25^, these results suggest that 5-hydroxythalidomide has potential similar to that of thalidomide for PLZF and SALL4 degradation in humans.

### Plzf is a substrate of Crbn with thalidomide in chicken

Thalidomide-induced teratogenesis occurs in several animals, including zebrafish, chicken, and rabbit^4,10^. The ZNF1/3 in PLZF (Fig3) and ZNF2 in SALL4^9,10^ are important regions for the interaction with CRBN. We thus compared these amino acid sequences among vertebrates. The sequences of the ZNF1 and ZNF3 and other ZNFs in PLZF are significantly conserved among many animals (Supplementary Fig. 6a), although the SALL4-ZNF2 sequence is not (Supplementary Fig. 6b). To investigate the thalidomide-dependent degradation of Plzf or Sall4 by Crbn from other animals, the recombinant proteins of Crbn, Plzf (Zbtb16 in chicken), and Sall4 from mouse (Mm) and chicken (Gg) were biochemically analysed. In addition, Val388 of human CRBN is a key residue for thalidomide-dependent CRBN–protein interaction^7,9,10^. Because the corresponding residue in mouse and chicken Crbn is isoleucine (Supplementary Fig. 6c), we substituted the Ile to Val in both, to produce MmCrbn-I391V and GgCrbn-I390V. In our biochemical analysis, the binding ability of HsCRBN-V388I was dramatically decreased compared with that of the wild-type protein (Supplementary Fig. 7a). Although MmCrbn did not bind with either protein in the presence of thalidomide (middle panel), interestingly, wild-type GgCrbn bound to GgPlzf following thalidomide treatment (right panel), whereas GgSall4 did not bind to GgCrbn with thalidomide. Both MmCrbn-I391V and GgCrbn-I390V indicated highly thalidomide-dependent binding with both Sall4 and Plzf.

In Supplementary Fig. 7a, thalidomide induced GgCrbn–GgPlzf interaction although it did not provide an interaction between MmCrbn and MmPlzf. To investigate this reason, we compared amino acid sequences of a thalidomide-binding region among human, mouse and chicken (Supplementary Fig. 6c). As a result, Glu377 in HsCRBN was conserved in GgCrbn but the Glu in MmCrbn was substituted to Vla. Actually, it was reported that Glu377 in HsCRBN was an important amino acid for interaction between CRBN and GSPT1^8^. Therefore, we investigated whether the Glu377 is important for the interaction between CRBN and PLZF or SALL4. In vitro binding assay showed that substitution of the Glu to Vla in HsCRBN significantly decreased binding ability to both SALL4 and PLZF (Supplementary Fig. 7b). In MmCrbn, double substitution of the Vla380 to Glu and the Ile391 to Vla significantly increased binding ability to Sall4 and Plzf, although single substitution of the Vla to Glu did not significantly increased (Supplementary Fig. 7c).

Next, each protein pair was transiently expressed in CRBN-deficient HEK293T cells. In the mouse pair, wild-type MmCrbn did not degrade either MmPlzf or MmSall4, whereas the MmCrbn-I391V and MmCrbn-V380E/I391V mutants degraded both (Fig. 5a and b, respectively). In the chicken pair, wild-type GgCrbn significantly induced thalidomi-dedependent degradation of GgPlzf (Fig. 5c), while degradation of GgSall4 was almost never observed (Fig. 5d). GgCrbn-I390V also degraded GgPlzf and GgSall4 with thalidomide (Fig. 5c and d, respectively). In contrast, the GgCrbn-E379V mutant did not induce degradation of GgPlzf (Supplementary Fig. 7d).

**Figure 4.**
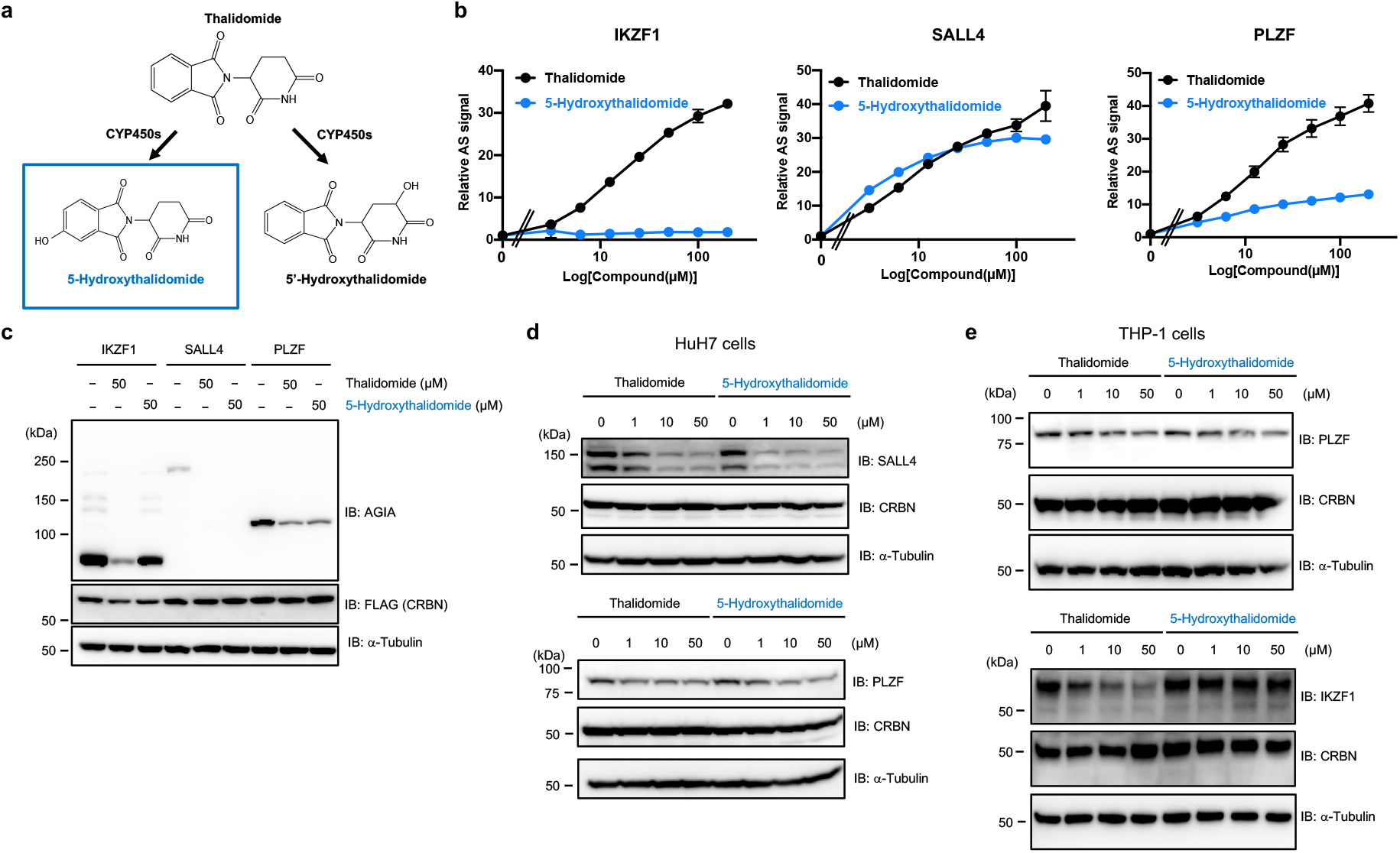
5-Hydroxythalidomide induces degradation of PLZF and SALL4 by CRBN. **a**, Schematic diagram of thalidomide metabolites by CYPs. **b**, *In vitro* binding assay for thalidomide and 5-hydroxythalidomide. Interaction between bls-CRBN and FLAG-GST-IKZF1, -SALL4, -PLZF in the presence of DMSO, thalidomide or 5-hydroxythalidomide (3.125, 6.25, 12.5, 25, 50, 100, or 200 μM) was analysed using AlphaScreen technology. **c**, Immunoblot analysis of AGIA-PLZF, AGIA-SALL4, or AGIA-PLZF in FLAG-CRBN expressing CRBN^-/-^ HEK293T cells treated with DMSO, thalidomide, or 5-hydroxythalidomide for 16 h. **d**, Immunoblot analysis of endogenous SALL4 or PLZF protein levels in HuH7 cells treated with DMSO, thalidomide, or 5-hydroxytahalidomide for 24 h. **e**, Immunoblot analysis of endogenous PLZF or IKZF1 protein levels in THP-1 cells treated with DMSO, thalidomide, or 5-hydroxytahalidomide for 24 h.

**Figure 5.**
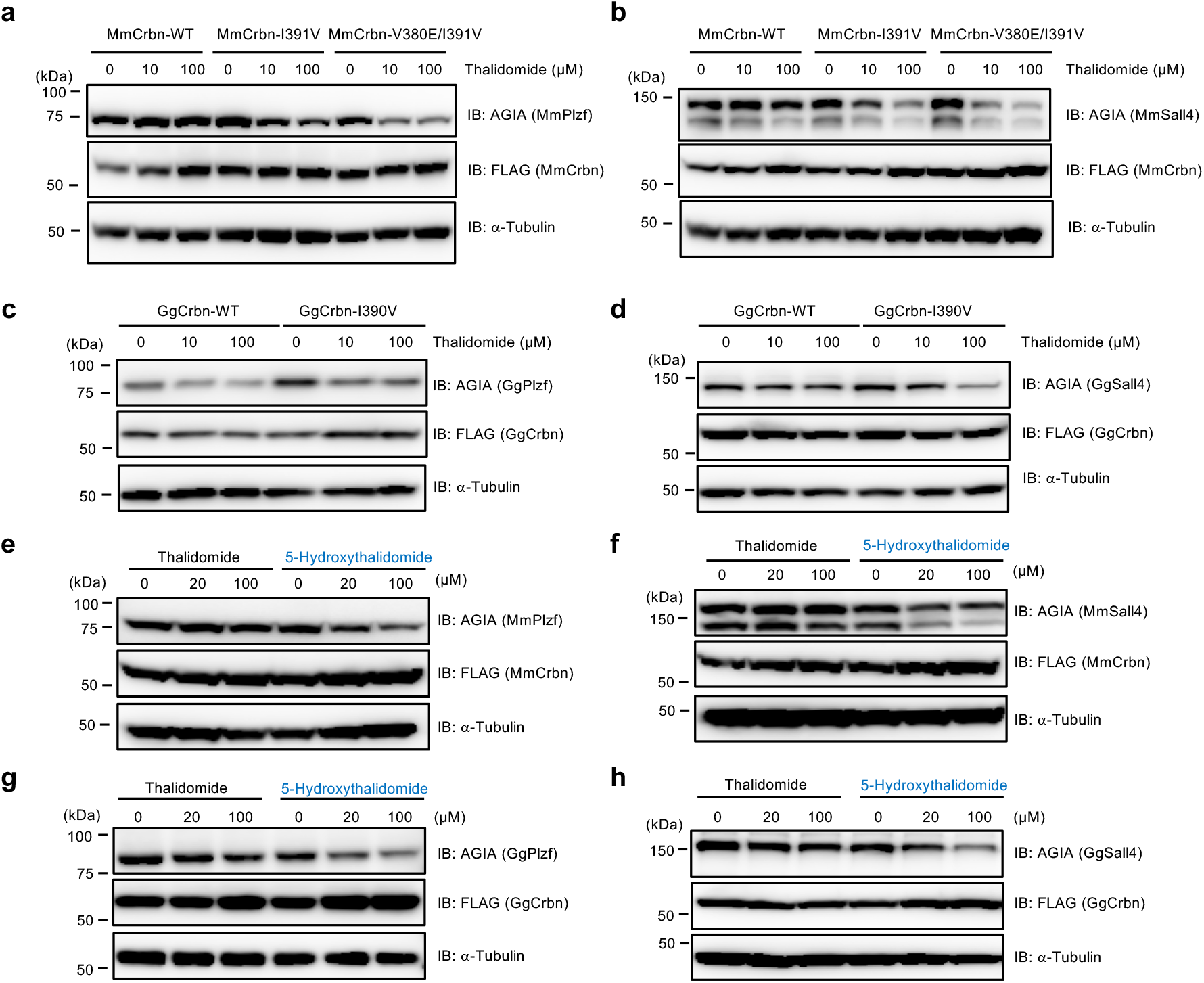
Crbn-dependent degradation of Plzf and Sall4 from mouse and chicken by treatment with thalidomide and 5-hydroxythalidomide. **a**, **b**, Immunoblot analysis of AGIA-MmPlzf (**a**) or -MmSall4 (**b**) in FLAG-MmCrbn-WT, - MmCrbn-I391V or -MmCrbn-V380E/I391V expressing CRBN^-/-^ HEK293T cells treated with DMSO or thalidomide for 16 h. **c**, **d**, Immunoblot analysis of AGIA-GgPlzf (**c**) or - GgSall4 (**d**) in FLAG-GgCrbn-WT or -GgCrb-I390V expressing CRBN^-/-^ HEK293T cells treated with DMSO or thalidomide for 16 h. **e**, **f**, Immunoblot analysis of AGIA-MmPlzf (**e**) or -MmSall4 (**f**) in FLAG-MmCrbn-WT expressing CRBN^-/-^ HEK293T cells treated with indicated concentration of DMSO, thalidomide or 5-hydroxythalidomide for 16 h. **g**, **h**, Immunoblot analysis of AGIA-GgPlzf (**g**) or -GgSall4 (**h**) in FLAG-GgCrbn-WT expressing CRBN^-/-^ HEK293T cells treated with indicated concentration of DMSO, thalidomide or 5-hydroxythalidomide for 16 h.

Taken together, we concluded that the Glu in thalidomide-binding region of CRBN is an important amino acid for thalidomide-dependent interaction with PLZF and SALL4, and that conservative amino acid sequence of CRBN-binding region with thalidomide in substrate proteins was also required, like wild-type GgCrbn could not induce degradation of GgSall4. In addition, these results suggest that thalidomide-dependent PLZF degradation occurs in many animals, including chickens and humans, while in contrast, SALL4 degradation by thalidomide may occur in a limited number of animals, including rabbits^10^, humans, and monkeys.

Recently, humanized-CYP3A mice were reported to show abnormal limb development after treatment with thalidomide^25^, suggesting that the thalidomide metabolites induce teratogenesis. We therefore investigated the function of 5-hydroxythalidomide on mouse and chicken Crbn-dependent degradation. Surprisingly, 5-hydroxythalidomide induced Crbn-dependent degradation of both Plzf (Fig. 5e and g) and Sall4 (Fig. 5f and h), whereas thalidomide had no function in MmCrbn and GgSall4 degradation by GgCrbn. Furthermore, thalidomide and 5-hydroxythalidomide did not induce downregulation of *PLZF* and *SALL4* mRNA expression in HuH7 cells (Supplementary Fig. 8). These results suggest that 5-hydroxythalidomide, rather than thalidomide, has a high potential for degradation of both Plzf and Sall4 in many animals.

### Plzf plays important roles in chicken limb development

It has been reported that both PLZF protein and *Plzf* mRNA are expressed in the limb buds of mouse and rat embryos^14,33^, suggesting direct function of Cbrn-thalidomide on PLZF in the developing limb bud. We thus investigated whether PLZF plays important roles in the development of chick limb bud. We first examined expression of *Plzf* gene in the chick limb bud and confirmed *Plzf* mRNA expression by whole mount *in situ* hybridization (Fig. 6a). Expression of *Sall4* and *Crbn* genes was also observed in the same region (Fig.6a). Expression of *Plzf* as well as *Sall4* mRNA was confirmed to be in limb mesenchyme by section in situ hybridization (Supplementary Fig. 9a). Then, to investigate whether downregulation of *Plzf* mRNA induces limb teratogenicity in chicken embryos, we constructed shRNA expression vector of GgPlzf. Immunoblot analysis showed that the constructed shRNA vector downregulated protein expression of overexpressed GgPlzf in DF-1 cells, which is chicken culture cells (Fig. 6b). Next, to elucidate developmental role of Plzf in chick limb development, RCAN retrovirus, which express shRNA (#2) against chick Plzf, was infected into blastderm cells containing prospective lateral plate mesoderm cells that gives rise to the limb bud. As shown Fig. 6c, it was confirmed that Plzf protein expression level was downregulated in the chicken embryos treated with Plzf shRNA by immunoblot analysis. In Fig. 6d, interestingly, Plzf shRNA infected limb bud showed several types of malformations (28%, n=32). Forelimb and hindlimb were shortened compared to control shRNA (GFP shRNA), infected limb bud (0%, n=10). We also observed that only one bone was formed in the zeugopod and digit number was also reduced in Plzf shRNA infected limb bud. These results suggest that chick Plzf has a pivotal role for limb bud outgrowth and that downregulation of Plzf causes teratogenicity.

**Figure 6.**
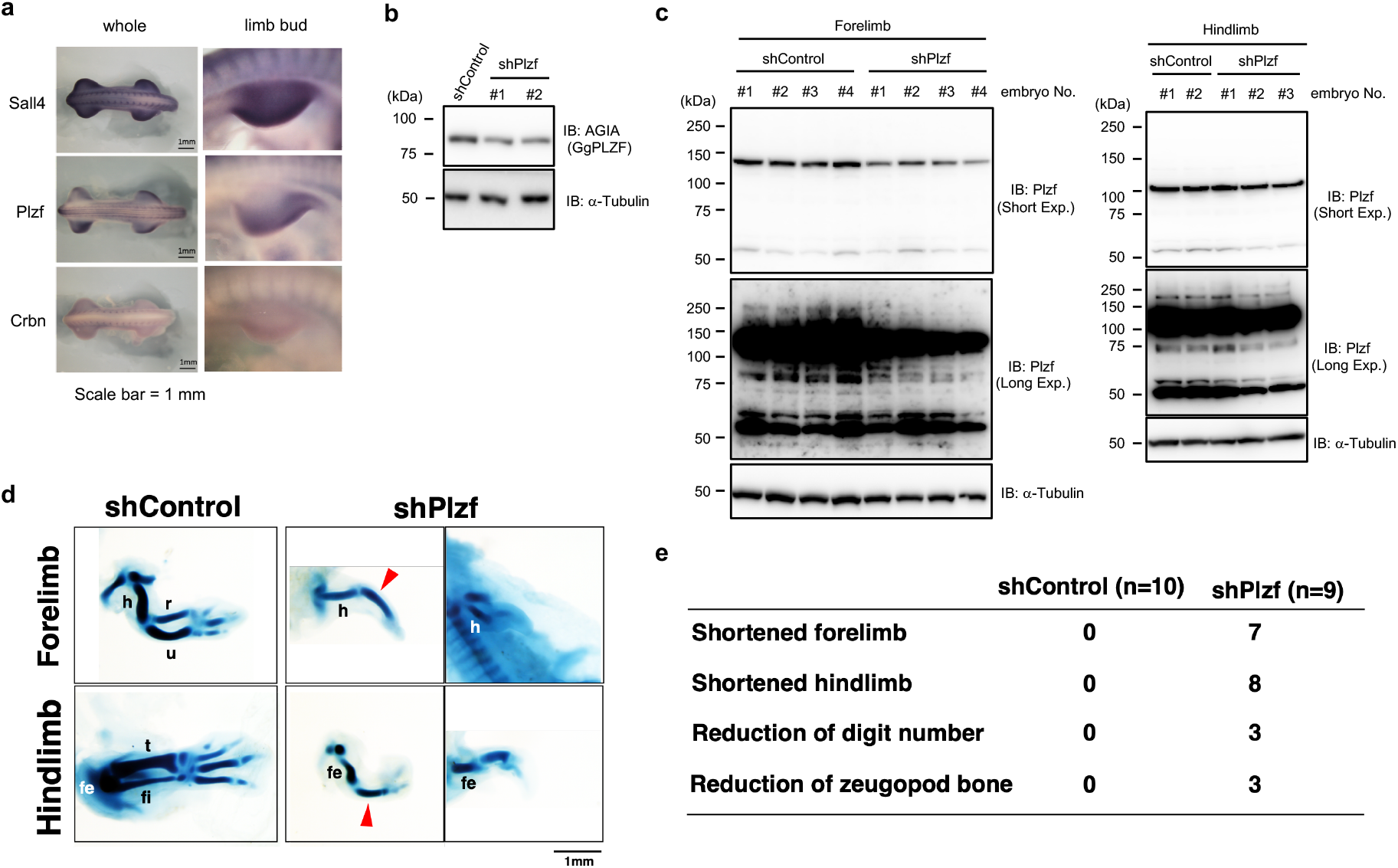
Downregulation of Plzf causes abnormal limb development in chicken embryo. **a,** *Sall4, Plzf* or *Crbn* mRNA expression in E4 chicken embryos was analysed by wholemount *in situ* hybridization. Left panel shows whole chicken embryo and right panel shows right forelimb bud. **b**, Immunoblot analysis of AGIA-GgPlzf in AGIA-GgPlzf expressing DF-1 cells transfected with shContol (shGFP) or shPlzf expression vector. **c**, Immunoblot analysis of Plzf from tissue of chicken forelimb or hindlimb bud. Endogenous Plzf protein expression was detected by immunoblot using chicken embryos infected with RCAN virus packaging shControl or shPlzf (forelimb shControl (n = 4), forelimb shPlzf (n = 4), hindlimb shControl (n = 2) or hindlimb shPlzf (n = 4)). **d**, Limb skeletal stained with Victoria blue. Skeletal patterning of forelimb and hindlimb in E6 chicken embryos infected RCAN virus packaging shControl (n = 10) or shPlzf (n = 9) were analysed by Victoria blue staining. h; humerus, r; radius, u; ulna, fe; femur, fi; fibula, t; tibia. **e**, Teratogenic phenotypes of chicken embryos in Fig. 6d.

### Thalidomide reduces Plzf protein levels in abnormal limb buds of chickens

Next, we investigated whether thalidomide targets PLZF in the chick limb bud. In previous reports, decreased expression of the *fgf8* gene is an indicator of abnormal limb bud development following thalidomide treatment^4^, and we confirmed reduced/skewed pattern of *fgf8* in truncated limb buds after thalidomide treatment (Supplementary Fig. 10a). In addition, limb skeletal defects were observed in this condition (Supplementary Fig. 10b). We investigated whether the amount of Plzf protein changes in these abnormal limb buds. Interestingly, reduced Plzf protein levels were observed in abnormal limb buds, although immunostaining of the Sall4 protein showed no change (Fig. 7a). Furthermore, to confirm the protein levels of Plzf and Sall4 in them, chick limb buds after thalidomide treatment were collected and characterized. Immunoblotting analysis also indicated that a reduction in Plzf was predominantly observed in teratogenic limb buds (T3, T8, and T18 in Fig. 7b and c). Taken together with the limb defects induced by human and mouse Plzf deficiency^15,16^, these results suggest that PLZF is a pivotal molecule for teratogenesis in chicken. Furthermore, we investigated whether 5-hydroxythalidomide induces teratogenicity in chicken embryo. As showed in Fig. 7d, 5-hydroxythalidomide also induced teratogenicity, and immunoblot analysis indicated that 5-hydroxythalidomide more strongly induced degradation of Plzf and Sall4, compared to thalidomide (Fig. 7e). These results suggest that 5-hydroxythalidomide plays an important role in thalidomide teratogenicity. The phenotypes of chicken embryos provided between 5-hydroxythalidomide and thalidomide were very similar (Fig. 7d), suggesting that degradation of other proteins including Sall4 is required much more to provide severe teratogenicity.

**Figure 7.**
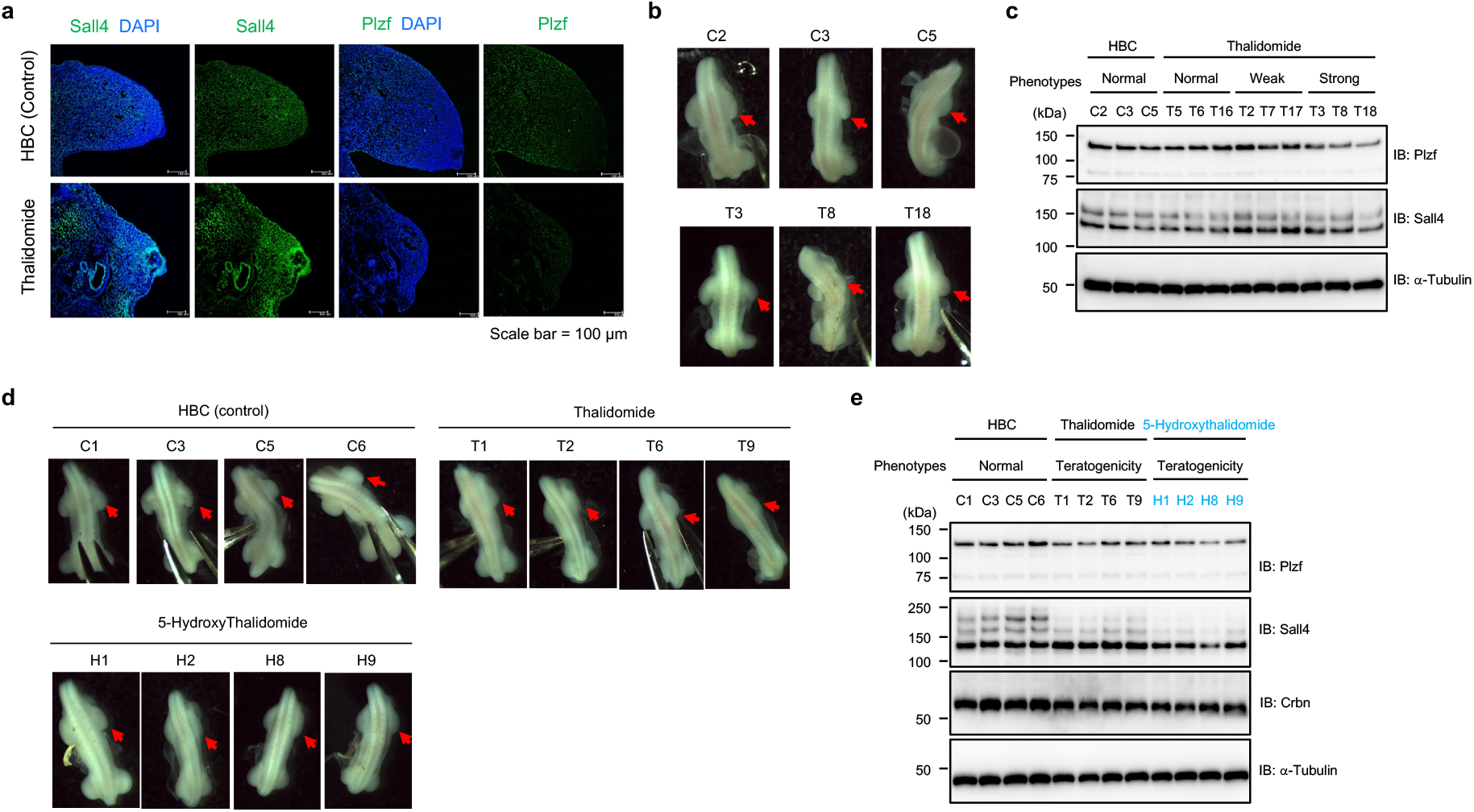
Thalidomide induces degradation of PLZF in abnormal chick limb buds. **a**, Immunohistochemical staining of Sall4 or Plzf in chicken forelimb bud. Endogenous Sall4 or Plzf protein expression was detected using forelimb bud section in chicken embryos treated with HBC (n = 4) or 1 μg/μl thalidomide (n = 4). **b**, Photographs show chicken embryos treated with HBC (control, n = 6, C2, C3, and C5) or strong phenotype (1 μg/μl thalidomide, n = 18, T3, T8, T18) corresponding to immunoblot analysis in Figure 7c. Red arrows show treated regions. **c**, Endogenous Plzf or Sall4 protein expression in chicken embryos in Fig. 7b was detected by immunoblot. **d,** Photographs show chicken embryos treated with HBC (control, n = 10, C1, C3, C5 and C6), 1 μg/μl thalidomide (n = 11, T1, T2, T6, T9) or 1 μg/μl 5-hydroxythalidomide (n = 10, H1, H2, H8, H9) corresponding to immunoblot analysis in Figure 6d. Red arrows show treated regions. **e**, Endogenous Plzf or Sall4 protein expression in chicken embryos in Fig. 7d was detected by immunoblot.

## Discussion

Many research groups have attempted to identify thalidomide-dependent substrates of CRBN^5–7,9,10,32^. The methodologies they have used have largely been based on cell-based assays such as SILAC (stable isotope labelling by amino acids in cell culture) and transient expression systems. In this study, we have used a biochemical assay using cell-free HuTFPA and AlphaScreen technologies. This method could easily detect thalidomide-dependent interactions between CRBN and PLZF, SALL4, or IKZF1, without the need for protein purification (Fig. 1), as well as the variations in the biochemical response of CRBN–protein depending on the presence of thalidomide, its derivatives, or 5-hydroxythalidomide (Fig. 3 to 5). Thalidomide-dependent interaction between CRBN and PLZF related to two ZNF domains (Fig. 3) and slowness of IMiD-dependent protein degradation, suggesting that the interaction manner is more complicated manner and may be different from known substrates, such as IKZF1, IKZF3, and SALL4. These results support a reason that PLZF has not been identified by method using cell-based assay or *in silico* so far, suggesting that our new biochemical approach using protein array could be useful in the identification. However, our assay system cannot identify substrate which requires the interaction with CRBN in protein complex. Therefore, our cell-free and conventional cell-based assay are complementary relationship each other, and we believe that the cell-free method can be used in the future research for exploration and confirmation of substrates with the cell-based assay. Furthermore, as IKZF1 interacted with CRBN in the presence of thalidomide but not 5-hdroxythalidomide, this method would be used for analysis of compound-dependent protein-protein interaction in development of novel drugs including thalidomide derivatives.

In this study, we identified PLZF as a new target of the thalidomide–CRBN system. In chicken embryo, downregulation of Plzf showed hypoplasia of limb bud (Fig. 6d and e), indicating that Plzf is required for proper chicken limb development. Furthermore, thalidomide and 5-hydroxythalidomide treatments decreased Plzf protein level but Sall4 was not induced protein degradation in the abnormal limb buds of chicken embryos (Fig. 7). According to these findings, we made a model for teratogenesis of chick limb buds (Fig. 8a). In mouse studies, Plzf^-/-^ deficient mice had major musculoskeletal limb defects^14^, and it was reported that PLZF deficient caused alteration of *Hoxds* or *Bmps* expression in developing limb^14^. Because Bmp proteins function as regulators in programming cell death and Plzf/Gli3 deficient induced cell death^15^, suggesting that degradation of PLZF by thalidomide treatment may affect cell proliferation in limb development. In contrast, Sall4 conditional knockout mice (Sall4-CKO) driven by T-Cre, which express early mesoderm, did not show any phenotype in the forelimb^12^. Furthermore, only 5% of Plzf^-/-^ mice showed forelimb phenotype which shows autopod abnormality^14^. These results indicate that thalidomide-induced teratogenicity in human cannot be explained by the results of each single knockout mice of Sall4 or Plzf. In previous report, Plzf and Gli3 double knockout mice showed phenotypes of remarkably reduction of stylopod and zeugopod^15^, and these phenotypes similar to phocomelia that is a typical phenotype of thalidomide embryopathy. It was reported that Gli3 expression was reduced in Sall4-CKO mice, therefore, we expect that double knockout mice of Sall4; T-Cre; Plzf^-/-^ would explain thalidomide phenotype in human patients. Our chick results showed that PLZF would be the pivotal target of thalidomide. Given that Sall4 protein was not degraded by thalidomide treatment in the chick embryo, it is thought that PLZF has more crucial function for normal development of the limb than Plzf in the mouse. Thus, variation of the protein sensitivity to the thalidomide and/or difference of the genes that are necessary for normal development of the limb between species would bring about difference of the phenotype between species in thalidomide teratology, which we showed in this study. PLZF that we found in this study as a new target of thalidomide will be a crucial target to solve thalidomide mystery between species in future.

**Figure 8.**
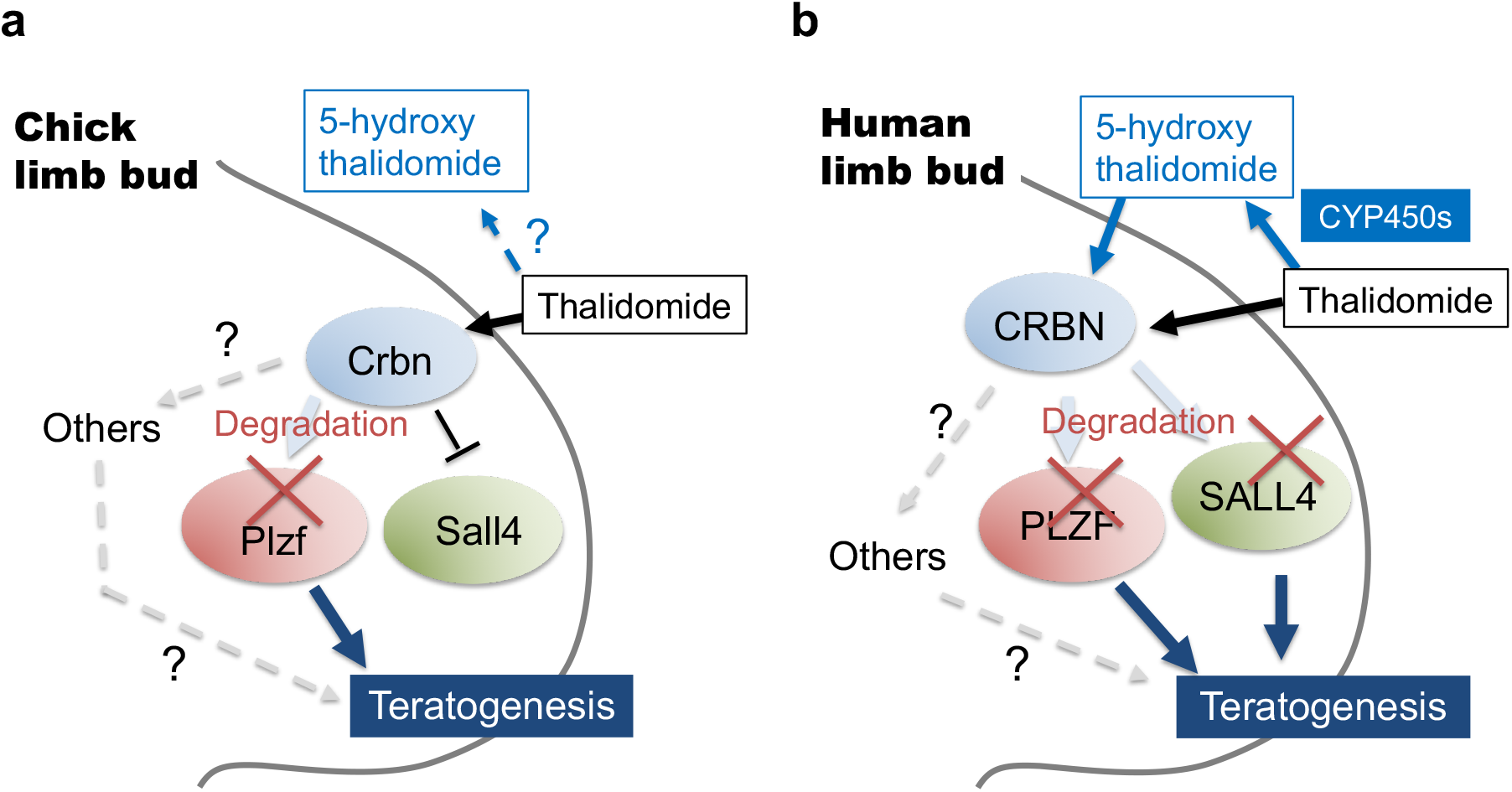
Model cartoon of thalidomide-induced teratogenicity in chicken or human limb bud. **a**, Model of thalidomide teratogenesis in chicken limb bud. **b**, Model of thalidomide teratogenesis in human limb bud.

PLZF is expressed in many cell types^34^, and is a multifunctional TF modulating many developmental biological processes^13^ including cellular proliferation and cell cycle control, myeloid and lymphoid cell development and differentiation, programming of NKT and iNKT cells, spermatogenesis and spermatogonial stem cell renewal, haematopoiesis, musculoskeletal-limb development, megakaryocytic development, and cytokine production. Notably, PLZF functions as a regulator for many immunoresponses^35–38^. Thalidomide is an immunomodulatory imide drug, and IKZF1 and IKZF3 are thought to be key targets of immunomodulation by thalidomide and its derivatives^39,40^. Therefore, dysfunction of PLZF by thalidomide may also result from its function as an IMiD, though further analysis will be required to confirm this.

Human CYP2C and 3A produce two metabolites: 5-hydroxythalidomide and 5’-hydroxythalidomide, as primary metabolites of thalidomide^18–20^. 5’-hydroxythalidomide has no effect on teratogenesis in chicken embryos^23^. However, it was no evidence whether metabolites from thalidomide induce CRBN-dependent protein degradation. In this study, 5-hydroxythalidomide induced degradation of both human PLZF and SALL4 proteins at the same level as thalidomide (Fig. 4). It is known that humans are highly sensitive to thalidomide^3,41^. These reports and our results suggest that 5-hydroxythalidomide has the potential for teratogenesis. For this reason, we used our findings to develop the hypothesis that the double degradations of PLZF and SALL4 by both thalidomide and 5-hydroxythalidomide produce high sensitivity to thalidomide in human embryopathy (Fig. 8b). We are convinced that researching the generation and action of 5-hydroxythalidomide will be important for understanding the function of thalidomide.

Thalidomide is a typical drug which shows species specificity^3^. From several researches, the species specificity of thalidomide has been thought to be provide by the difference of a thalidomide-binding sequence in CRBN^7,9,10^. Consistent with these reports, in this study, humanized-mouse and chicken Crbns indicated thalidomide-dependently interactions and degradation of both Plzf and Sall4 proteins (Fig. 5a-d and Supplementary Fig. 7). In addition, with the sequence difference in Crbn, it has been thought that the metabolism cascade of thalidomide is also important in the species specificity^3^. In this study, 5-hydroxythalidomide showed higher potential than thalidomide for degradations of both Plzf and Sall4 proteins from mouse and chicken (Fig. 5e-h, Fig. 7d and e). Furthermore, humanized-CYP3A mice have been recently shown to undergo abnormal limb development at a high rate (>40%) after thalidomide treatment^25^, although in general thalidomide does not produce teratogenesis in mice^3^. These reports and our results suggest that thalidomide metabolism cascade, including metabolism speed and kinds of metabolites, plays an important role in the species specificity of thalidomide.

## Methods

### Reagents

Thalidomide (Sigma-Aldrich and Tokyo Chemical Industry Co., Ltd), Pomalidomide (Sigma-Aldrich), Lenalidomide (FUJIFILM Wako Pure Chemical), 5-hydroxythalidomide (5-hydroxythalidomide was prepared according to a previously published method^20^), MG132 (Peptide Institute), and MLN4924 (Chemscene) were dissolved in DMSO (FUJIFILM Wako Pure Chemical) at 2 to 100 mM and stored at −20°C as stock solutions. All drugs were diluted 1,000, 500, or 250-fold for *in vivo* experiments, or diluted 200-fold for *in vitro* experiments.

### Production of recombinant proteins using the cell-free system

*In vitro* transcription and wheat cell-free protein synthesis were performed using a WEPRO1240 expression kit (Cell-Free Sciences). Transcripts were conducted using SP6 RNA polymerase with plasmids or DNA fragments as templates. The translation reaction was performed in bilayer mode using a WEPRO1240 expression kit (Cell-Free Sciences), according to the manufacturer’s instructions. For biotin labelling, 1 μl of cell-free synthesized crude biotin ligase (BirA), produced by the wheat cell-free expression system, was added to the lower layer, and 0.5 μM (final concentration) of d-biotin (Nacalai Tesque) was added to both the upper and lower layers, as described previously^42^.

### Interaction analysis of CRBN-IMiD-substrate using AlphaScreen technology

IMiD at the concentrations indicated in each figure and 0.5 μl of biotinylated HsCRBN, MmCrbn, or GgCrbn were mixed in a 15 μl of AlphaScreen buffer containing 100 mM Tris (pH 8.0), 0.01% Tween20, 100 mM NaCl, and 1 mg/ml BSA. Then, 5 μl of substrate mixture containing 0.8 μl of FLAG-GST-substrate in AlphaScreen buffer was added, and 20 μl of the reaction mixture was incubated at 26°C for 1 h in a 384-well AlphaPlate (PerkinElmer). Subsequently, 5 μl of detection mixture containing 0.2 μg/ml anti-DYKDDDDK mouse mAb (Wako), 0.08 μl of streptavidin-coated donor beads, and 0.08 μl of Protein A-coated acceptor beads (PerkinElmer) in AlphaScreen buffer, were added to each well. After incubation at 26°C for 1 h, luminescent signals were detected using an EnVision plate reader (PerkinElmer).

### Production of the human transcription factor protein array (HuTFPA)

For the construction of human TF protein array, we prepared pEU-E01-FLAG-GST-K1-02 vector containing FLAG tag, GST tag, SG linker, and *Asi*SI restriction enzyme site at 5’ upstream of multiple cloning site. cDNA clones coding proteins with DNA-binding domains were selected from cDNA resources collected by Kazusa DNA research institute^43^ (Supplementary Table 1). The plasmid of each clone was digested by combination of *Asi*SI and an appropriate restriction enzyme such as XhoI, SalI or NotI. The DNA fragment was inserted into pEU-E01-FLAG-GST-K1-02 vector digested by same restriction enzymes. After subcloning, pEU expression plasmids were arranged in 96 well format and stored as glycerol stock. Transcription template DNA fragments were amplified directly by PCR using PrimeStar Max PCR polymerase (Takara Bio), SPu-2 (5’–CAGTAAGCCAGATGCTACAC) and AODA2306 (5’–AGCGTCAGACCCCGTAGAAA) primers and diluted glycerol stocks as template. Transcription and translation reactions were conducted using WEPRO7240 expression kit (Cell-Free Sciences) in micro-titer plate format. Transcription reaction mixture was prepared by mixing 1.4 μl of transcription buffer LM, 0.7 μl of NTP mixture (25 mM each), 0.07 μl RNase Inhibitor (Promega), 0.26 μl SP6 polymerase (Promega) and 1.4 μl PCR product in 96 well plate. The transcription reaction was incubated at 37°C for 18 h. Translation reaction mixture containing 2.5 μl of mRNA, 1.67 μl of WEPRO 7240 wheat germ extract, 0.14 μl of creatine kinase (20 mg/ml) (Roche diagnostics) and 0.11 μl RNase Inhibitor was prepared and overlaid with 44 μl of SUB-AMIX SGC solution (Cell-Free Sciences) in V-bottom 384 well plate. The translation reaction was incubated at 26°C for 18 h. Expression of each protein product was confirmed by Western blotting using anti-DYKDDDDK tag antibody (FUJIFILM Wako Pure Chemical).

### High-throughput screening using AlphaScreen technology

We added 20 μl of bait mixture, containing 50 μM thalidomide and 0.5 μl of biotinylated HsCRBN in AlphaScreen buffer, to 384-well AlphaPlates using a FlexDrop Precision Reagent Dispenser (PerkinElmer). We next added 0.8 μl of FLAG-GST-transcription factor proteins to 384-well AlphaPlates using a NanoHead (PerkinElmer) and a Janus Workstation (PerkinElmer). After the 384-well AlphaPlates were incubated at 26°C for 1 h, 5 μl of detection mixture containing 0.2 μg/ml anti-DYKDDDDK mouse mAb (FUJIFILM Wako Pure Chemical), 0.08 μl of streptavidin-coated donor beads, and 0.08 μl of Protein A-coated acceptor beads (PerkinElmer) in AlphaScreen buffer, were added to each well using a FlexDrop precision reagent dispenser. After incubation at 26°C for 1 h, luminescent signals were detected using an EnVision plate reader (PerkinElmer).

### Plasmids

Plasmids pDONR221 and pcDNA3.1(+), based on Gateway technology, were purchased from Invitrogen, and the pEU vector for the wheat cell-free system was constructed in our laboratory, as previously described^26^. Plasmids pcDNA3.1(+)-FLAG-GW, pcDNA3.1(+)-FLAG-MCS, pcDNA3.1(+)-AGIA-MCS, pEU-bls-GW, and pEU-bls-MCS were constructed based on each original vector by PCR and using the In-Fusion system (Takara Bio), or PCR and restriction enzymes. pEU-FLAG-GST-IKZF1, -SALL4, -PLZF, - SALL1, -SALL2, -ZBTB17, -ZBTB20, -ZBTB38, -ZBTB48, -HNRNPK, -HMGB2, and ELF5 were purchased from the Kazusa DNA Research Institute. Plasmids HsSALL4, HsPLZF, and IKZF1 were amplified and restriction enzyme sites were added by PCR and cloned into pcDNA3.1(+)-AGIA-MCS. The open reading frame of HsCRBN was purchased from the Mammalian Gene Collection (MGC) and MmCrbn, MmSall4, and MmPlzf were did from Functional Annotation of Mouse (FAMTOM), respectively^28,31^. The open reading frame of GgCrbn was artificially synthesized by IDT, and pcDNA3.1(+)-GgSall4-DYKDDDDK and pcDNA3.1(+)-GgPlzf-DYKDDDDK were purchased from GenScript. HsCRBN was amplified and the BP reaction sequence (attB and attP) was added by PCR and cloned into pDONR221 using BP recombination (Invitrogen). Then, pDONR221-HsCRBN was recombined to pEU-bls-GW or pcDNA3.1(+)-FLAG-GW using LR recombination (attL and attR). MmCrbn and GgCrbn were amplified and restriction enzyme sites were added by PCR and cloned into pEU-bls-MCS or pcDNA3.1(+)-FLAG-MCS. MmSall4, GgSall4, MmPlzf, and GgPlzf were amplified and restriction enzyme sites were added by PCR and cloned into pEU-FLAG-GST-MCS or pcDNA3.1(+)-AGIA-MCS. Domain swapped HsPLZF was constructed by inverse PCR and the In-Fusion system (Takara Bio). Deletion mutation and amino acid mutation of each protein was performed by inverse PCR.

For *in situ* hybridization, the cDNAs for chicken *Fgf8, Crbn, Sall4*, and *Plzf* were obtained by PCR using the following primers: *Fgf8* (NM_001012767.1), 5’-attacgcgtATGGACCCCTGCTCCTCGCTCTTCA-3’ and 5’-attgataTCATGGGCGCAGGGAGGCGCTGGAG-3’; *Crbn* (XM_015293204.2), 5’-ctataggctagaattcacgcgtATGGCCGCCGAGGAGGGAGGTGACGGA-3’ and 5’-cactaaagggaagcggccgcgatatcTTACAAGCAGAGTAACGGAGATC-3’; *Sall4* (NM_001080872.1), 5’-ctataggctagaattcacgcgtATGTCGCGACGGAAGCAGGCGAAGCCC-3’ and 5’-cactaaagggaagcggccgcgatatcTTAACTAACGGCAATTTTGTTCT-3’; *Plzf* (XM_015298212.1), 5’-ctataggctagaattcacgcgtATGGATTTGACTAAGATGGGCATGATA-3’ and 5’-cactaaagggaagcggccgcgataTCAGACGTAGCAGAGGTAGAGATAG-3’. The amplified fragments of *Fgf8* were digested by MluI-EcoRV, and subcloned into pCMS-EGFP vector (Clontech). The amplified fragments of *Crbn, Sall4*, and *Plzf* were inserted into MluI-EcoRV site of pCMS-EGFP vector by In-fusion (Takara Bio).

For knockdown of chick Plzf, the shRNA sequences of chick Plzf (#1: 5’-GGAAATCGAGGTACATCAAGG-3’ or #2: 5’-GATTACTCGGCCATGATCAAA-3’) were used. The following DNA oligos: #1: 5’-gatcccGGAAATCGAGGTACATCAAGGgcttcctgtcacCCTTGATGTACCTCGATTTCCttt ttta-3’ and 5’-agcttaaaaaaGGAAATCGAGGTACATCAAGGgtgacaggaagcCCTTGATGTACCTCGATTTCCgg-3’; #2: 5’-gatcccGATTACTCGGCCATGATCAAAgcttcctgtcacTTTGATCATGGCCGAGTAATCttt ttta-3’ and 5’-agcttaaaaaaGATTACTCGGCCATGATCAAAgtgacaggaagcTTTGATCATGGCCGAGTAATgg-3’ were purchased from Invitrogen. The DNA oligo pairs were annealed and inserted into pEntryCla12-chickU6 shuttle vector using BamHI/HindIII site.

### Cell culture and transfection

HEK293T cells were cultured in DMEM (low glucose) medium (FUJIFILM Wako Pure Chemical) supplemented with 10% fetal bovine serum (FUJIFILM Wako Pure Chemical), 100 unit/ml penicillin, and 100 μg/ml streptomycin (Gibco) at 37°C under 5% CO_2_. HEK293T cells were transfected using *Trans*IT-LT1 transfection reagent (Mirus Bio) or PEI Max: Polyethyleneimine “Max” (MW 40,000) (PolyScience, Inc.).

HuH7 cells were cultured in DMEM (high glucose) medium (FUJIFILM Wako Pure Chemical) supplemented with 10% fetal bovine serum (Wako), 100 unit/ml penicillin, 100 μg/ml streptomycin (Gibco), 1 mM Sodium Pyruvate (Gibco), 10 mM HEPES (Gibco), and 1 × MEM NEAA (Gibco) at 37°C under 5% CO_2_.

THP-1 cells were cultured in RPMI160 GlutaMAX medium (Gibco) supplemented with 10% fetal bovine serum (FUJIFILM Wako Pure Chemical), 100 unit/ml penicillin, and 100 μg/ml streptomycin (Gibco) at 37°C under 5% CO_2_.

TK, HT, BJAB, SU-DHL-4, MT-4, Raji cells were cultured in RPMI1640 GlutaMAX medium supplemented with 10% fetal bovine serum (FUJIFILM Wako Pure Chemical), 100 unit/ml penicillin, 100 μg/ml streptomycin (Gibco), and 55 μM 2-Mercaptoethanol (Gubco) at 37°C under 5% CO_2_.

DF-1 cells were cultured in DMEM (low glucose) medium (FUJIFILM Wako Pure Chemical) supplemented with 10% fetal bovine serum (Wako), 100 unit/ml penicillin, and 100 μg/ml streptomycin (Gibco) at 37°C under 5% CO_2_. DF-1 cells were transfected using *Trans*IT-LT1 transfection reagent (Mirus Bio).

### Immunoblot and antibodies

Protein lysates were separated by SDS-PAGE and transferred onto polyvinylidene difluoride membranes (Millipore). After the membranes were blocked using 5% skimmed milk (Megmilk Snow Brand) in TBST (20 mM Tris-HCl pH 7.5, 150 mM NaCl, 0.05% Tween20) at room temperature for 1 h, the following antibodies were used. Anti-FLAG mouse mAb (HRP-conjugated, Sigma-Aldrich, A8592), anti-AGIA rabbit mAb^31^ (HRP-conjugated, produced in our laboratory) were used to detect epitope-tagged proteins. Anti-α-tubulin rabbit pAb (HRP-conjugated, MBL, PM054-7) was used to detect α-tubulin. Biotinylated proteins were detected by anti-biotin (HRP-conjugated, Cell Signaling Technology, #7075). Anti-CRBN rabbit mAb (Cell Signaling Technology, #71810), anti-PLZF rabbit mAb (Cell Signaling Technology, #39784), anti-PLZF rabbit pAb (GeneTex, GTX111046), anti-SALL4 rabbit pAb (Abcam, ab29112), anti-SALL4 mouse mAb (Santa Cruz Biotechnology, sc-101147), anti-DDB1 mouse mAb (Santa Cruz Biotechnology, sc-376860), anti-CUL4 mouse mAb (Santa Cruz Biotechnology, sc-377188), anti-RBX1 mouse mAb (Santa Cruz Biotechnology, sc-393640), and anti-ubiquitin mouse mAb (P4D1, Cell Signaling Technology, #3936) were used as primary antibodies. Anti-rabbit IgG (HRP-conjugated, Cell Signaling Technology, # 7074) and anti-mouse IgG (HRP-conjugated, Cell Signaling Technology, # 7076) were used as secondary antibodies. Immobilon (Millipore) or ImmunoStar LD (FUJIFILM Wako Pure Chemical) was used as substrate HRP and luminescent signals were detected using an ImageQuant LAS 4000mini (GE Healthcare). To perform re-probing, Stripping Solution (FUJIFILM Wako Pure Chemical) was used and re-blocked using 5% slim milk in TBST.

For immunoblot analysis of extract from chicken embryo, a right forelimb bud was dissected from HH st. 22/23 embryos, and boiled in 50 μl of buffer (50 mM Tris-HCl pH 7.5, 4% SDS) at 98°C for 10 min. Protein concentration of each lysate was quantified using BCA assay (Thermo Fisher Scientific).

### *In vitro* pull-down assay of CRBN and substrate

To confirm the thalidomide-dependent interactions between IKZF1, PLZF, or SALL4 and CRBN, we performed pull-down assays using Dynabeads M-280 Streptavidin (Invitrogen). Biotinylated CRBN-WT and CRBN-YW/AA were synthesized using the wheat cell-free system as described above. We then mixed 5 μl of Dynabeads M-280 Streptavidin with 5 μl of biotinylated CRBN-WT or CRBN-YW/AA and diluted this 10-fold with PBS containing 0.05% Tween20, and incubated this at room temperature for 1 h. The beads were washed three times in 500 μl PBS containing 0.05% Tween20 and substrate-thalidomide mixture was added containing 10 μl of FLAG-GST-IKZF1, -SALL4 or -PLZF, and 200 μM thalidomide (0.5% DMSO) in 300 μl of AlphaScreen buffer containing 100 mM NaCl, 0.01% Tween20, and 1 mg/ml BSA. After rotation at room temperature for 90 min, the beads were washed four times in 500 μl of 1× Lysis buffer (50 mM Tris-HCl pH 7.5, 150 mM NaCl, 1% Triton X-100) and proteins were eluted by boiling in 1× sample buffer (62.5 mM Tris-HCl pH 6.8, 2% SDS, 10% glycerol) containing 5% 2-mercaptoethanol. The proteins were then analysed by immunoblot.

### Construction of CRBN-KO HEK293T cells

The guide nucleotide sequence 5’– ACTCCGGGCGGTTACCAGGC-3’ was selected from the human CRBN gene. The Guide-it plasmid vector (Takara Bio) was used to construct CRBN-KO cells. We then cultured HEK293T cells in a 6-well plate and transfected the plasmid into them. Two days after transfection, GFP positive cells were sorted by FACSAria (Becton, Dickinson and Company) and cell clones were obtained by limiting dilution. Genomic DNA was then isolated and the mutation was confirmed by sequencing after TA cloning (Toyobo).

### *In vivo* IMiD-dependent degradation assay of substrates

To confirm IMiD-dependent degradation of PLZF, HEK293T or HEK293T-CRBN^-/-^ cells were cultured in 48-well plates and transfected with 200 ng pcDNA3.1(+)-FLAG-CRBN-WT or 200 ng pcDNA3.1(+)-FLAG-CRBN-YW/AA and 20 ng pcDNA3.1(+)-AGIA-PLZF or -AGIA-PLZF variants, or -AGIA-SALL4. After the cells were transfected for 8 h, they were treated with IMiD or DMSO (0.1%) in culture medium, at the times and concentrations indicated in each figure.

To show that IMiD-dependent PLZF degradation is caused by CRL and the 26S proteasome, the cells were treated with 2 μM MLN4924 and 10 μM MG132 (0.2% DMSO) at the times indicated in Fig. 2a.

To examine the degradation of endogenous PLZF or SALL4, we cultured HEK293T, HuH7, or THP-1 cells in 24 or 48-well plates and treated them with lenalidomide or DMSO (0.1%) in culture medium at the times and concentrations indicated in each figure.

To examine the degradation of endogenous PLZF in TK, HT, BJAB, SU-DHL-4, MT-4, Raji cells, we cultured induced the lymphoma cells in 12-well plate and treated then with lenalidomide, pomalidomide or DMSO (0.1%) in culture medium at the times and concentrations indicated in Supplementary Fig.5. Then, the cells were lysed with XXX μL of RIPA buffer containing a protease inhibitor cocktail (Sigma-Aldrich). Protein concentration of each lysate was quantified using BCA assay (Thermo Fisher Scientific).

To examine 5-hydroxythalidomide-dependent degradation of overexpressed PLZF, SALL4 or IKZF1, HEK293T-CRBN^-/-^ cells were cultured in 48-well plates and transfected with 200 ng pcDNA3.1(+)-FLAG-CRBN-WT and 20 ng pcDNA3.1(+)-AGIA-SALL4, - PLZF, or -IKZF1. After the cells were transfected for 8 h, they were treated with thalidomide, 5-hydroxythalidomide, or DMSO (0.1%) in culture medium at the indicated times and concentrations in Fig. 4c. For endogenous SALL4, PLZF, and IKZF1, HuH7 or THP-1 cells were cultured in 48-well plates and treated with thalidomide, 5-hydroxythalidomide, or DMSO (0.1%) in culture medium at the indicated times and concentrations in Fig. 4d and 4e.

To examine the species specificity of IMiD-dependent protein degradation, HEK293T-CRBN^-/-^ cells were cultured in 48-well plates and transfected with 200 ng pcDNA3.1(+)-FLAG-(mouse or chicken) Crbn-WT or -IV and 20 ng pcDNA3.1(+)-AGIA-(mouse or chicken) Plzf or -(mouse or chicken) Sall4. After the cells were transfected for 8 h, they were treated with thalidomide or DMSO (0.2%) in culture medium for the times and concentrations indicated in Fig. 5a-d.

To examine whether 5-hydroxythalidomide induced the degradation of (mouse or chicken) Sall4 or Plzf, HEK293T-CRBN^-/-^ cells were cultured in 48-well plates and transfected with 200 ng pcDNA3.1(+)-FLAG-(mouse or chicken) Crbn-WT and 20 ng pcDNA3.1(+)-AGIA-(mouse or chicken) Sall4 or -(mouse or chicken) Plzf. After the cells were transfected for 8 h, they were treated with thalidomide, 5-hydroxythalidomide or DMSO (0.2%) in culture medium for the times and concentrations indicated in Fig. 5e-h.

In all experiments, cells were lysed by boiling in 1× sample buffer containing 5% 2-mercaptoethanol, and the lysates were analysed by immunoblot.

### Quantitative RT-PCR

To demonstrate PLZF protein level decrease results from post-translational event, *PLZF* mRNA expression in HEK293T, HuH7, or THP-1 cells treated with DMSO or lenalidomide for 24 h, were assessed by quantitative Real-Time PCR (qRT-PCR). To analyse *SALL4* or *PLZF* mRNA expression, HuH7 cells were treated with DMSO, thalidomide or 5-hydroxythalidomide for 24 h, were assessed by qRT-PCR. Total RNA was isolated from HEK293T, HuH7, or THP-1 cells treated with DMSO or lenalidomide for 24 h using a SuperPrep cell lysis kit (Toyobo) and cDNA was synthesized using a SuperPrep RT kit (Toyobo), according to the manufacture’s protocol. RT-PCR was performed using a KOD SYBR qPCR Mix (Toyobo) and data was normalized against GAPDH mRNA levels. PCR primers are as follows: *PLZF* FW 5’-GCACAGTTTTCGAAGGAGGA-3’, *PLZF* RV 5’-GGCCATGTCAGTGCCAGT-3’, *SALL4* FW 5’-GGTCCTCGAGCAGATCTTGT-3’, *SALL4* RV 5’-GGCATCCAGAGACAGACCTT-3’, *GAPDH* FW 5’-AGCAACAGGGTGGTGGAC-3’, *GAPDH* RV 5’-GTGTGGTGGGGGACTGAG-3’

### Co-immunoprecipitation of CRL4^CRBN^-IMiD-PLZF

To examine whether PLZF interacts with CRL4^CRBN^-thalidomide or -lenalidomide, HEK293T cells were cultured in a 15-cm dish and transfected with 12 μg pcDNA3.1(+)-FLAG-CRBN and 12 μg pcDNA3.1(+)-AGIA-PLZF. After HEK293T cells were treated with DMSO, 10 μM or 100 μM thalidomide or lenalidomide in the presence of 10 μM MG132 for 8 h, cells were lysed in 1.5 mL IP Lysis buffer (Pierce) (25 mM Tris-HCl pH 7.5, 150 mM NaCl, 1 mM EDTA, 1% NP-40, 5% glycerol) containing a protease inhibitor cocktail (Sigma-Aldrich) and incubated on ice for 15 min. Lysates were clarified by centrifugation at 13,000 rpm for 15 min, and CRL^CRBN^ was immunoprecipitated using anti-FLAG M2 magnetic beads (Sigma-Aldrich) with DMSO or 10 μM or 100 μM thalidomide or lenalidomide in the presence of 10 μM MG132. After overnight rotation at 4°C, the beads were washed three times with 800 μL of IP Lysis buffer (Pierce) and the proteins were eluted with 1× sample buffer. After eluted proteins were transferred to another tube, 2-mercaptoethanol (final concentration is 5%) was added, and boiled at 98°C for 5 min. The proteins were then analysed by immunoblot.

### *In vivo* ubiquitination assay

To detect polyubiquitination of PLZF in cells, HEK293T cells were cultured in a 10-cm dish and transfected with 5 μg pcDNA3.1(+)-FLAG-CRBN and 5 μg pcDNA3.1(+)-AGIA-PLZF. After HEK293T cells were treated with DMSO, 10 μM or 100 μM thalidomide or lenalidomide in the presence of 10 μM MG132 for 10 h, cells were lysed in 600 μl of SDS Lysis buffer (50 mM Tris-HCl pH 7.5, 1% SDS) containing a protease inhibitor cocktail (Sigma-Aldrich) and boiled at 90°C for 15 min. Denatured lysates were treated with Benzonase Nuclease (Sigma-Aldrich) at 37°C for 1 h, and the lysates were clarified by centrifugation at 13,000 rpm for 15 min, then diluted 10-fold with IP Lysis buffer (Pierce). The proteins were immunoprecipitated overnight with Dynabeads Protein G (Invitrogen) interacting anti-AGIA antibody at 4°C, which were then washed three times with 800 μl of IP Lysis buffer (Pierce). Proteins were eluted by boiling in 1× sample buffer containing 5% 2-mercaptoethanol. The proteins were then analysed by immunoblot.

### *In vitro* binding and ubiquitination assay

To confirm that PLZF is a direct substrate, we performed an *in vitro* binding and ubiquitination assay as described previously^6^. HEK293T cells were cultured in a 15-cm dish and transfected with 28 μg pcDNA3.1(+)-FLAG-CRBN or empty vector. HEK293T cells were cultured in two 15-cm dishes and transfected with 25 μg pcDNA3.1(+)-AGIA-PLZF per one dish. After 24 h, the cells were lysed in 1.6 ml/dish of IP Lysis buffer (Pierce) containing a protease inhibitor cocktail (Sigma-Aldrich), and 400 μl of FLAG-CRBN or empty vector lysates were mixed with 400 μL of AGIA-PLZF lysates in the presence of DMSO or 100 μM lenalidomide. The FLAG-CRBN and AGIA-PLZF mixtures were immunoprecipitated using anti-FLAG M2 magnetic beads by rotating overnight at 4°C. As a negative control, empty vector and AGIA-PLZF mixtures were immunoprecipitated using anti-AGIA-conjugated magnetic beads (produced in our laboratory) by rotating overnight at 4°C. The beads were washed three times with 800 μl of IP Lysis buffer (Pierce) and washed twice with 600 μl of 1× ubiquitin reaction buffer (50 mM Tris-HCl pH 7.5, 5 mM KCl, 5 mM MgCl2, 0.5 mM DTT), then resuspended in 20 μl of 1× ubiquitin reaction buffer containing 200 nM UBE1 E1 (R&D systems, U-110), 1 μM UbcH5a E2 (Enzo, BML-UW9050-100), 1 μM UbcH5b (Enzo, BML-UW9060-100), 5 mM ATP, 10 μM HA-ubiquitin (BostonBiochem, U-110), 10 μM MG132, protease inhibitor cocktail, and DMSO or 200 μM lenalidomide. Then, *in vitro* ubiquitination was performed at 30°C for 3 h, the proteins were denatured in 2% SDS by boiling at 95°C for 15 min. The proteins were diluted 20-fold with IP Lysis buffer (Pierce) and immunoprecipitated anti-AGIA-conjugated magnetic beads (produced in our laboratory) at 4°C for 4 h. The beads were washed four times with 800 μl of IP Lysis buffer (Pierce) and the proteins were eluted with 750 μl (lane 1-4) or 20 μl (lane 5-6) of 1× sample buffer. After elution, proteins were transferred to another tube, 2-mercaptoethanol (final concentration is 5%) was added, and were boiled at 98°C for 5 min. The proteins were then analysed by immunoblot.

### Plzf knockdown experiment in DF-1 cells

To confirm efficiency of shRNA, DF-1 cells were cultured in a 48-well plate and transfected with 50 ng pcDNA3.1(+)-AGIA-Ggplzf and 400 ng shRNA vector. After 6 h, culture medium was exchanged with new culture medium and the DF-1 cells were harvested after 48 h of transfection. The lysates were denatured by boiling at 98°C for 5 min and analysed by immunoblot.

### Animals

Fertilized eggs of white leghorn chicken (*Gallus gallus domesticus*) were purchased from a domestic poultry farm (Kakeien, Sendai, Japan). Eggs were incubated at 38°C until appropriate developmental stage. Embryos were staged according to the criteria made by Hamburger and Hamilton^44^. All animal experiments were properly conducted in accordance with the guidelines of Tohoku University.

### Knockdown of Plzf in chicken embryos

5 μg of RCAN(A) retrovirus vector containing chick U6 promoter which expresses shRNA for *gfp* or *Plzf* was transfected into M/O chicken strain-derived embryonic fibroblast cells (CEF) cultured in 60-mm dish using Fugene HD transfection regent (Promega). CEF was cultured in DMEM-high glucose containing 10% FBS, 2% chicken serum, 1% penicillin-streptomycin, and spread three times. After ten 10-cm dishes reached to the confluent, medium was changed into DMEM-high glucose containing 2% FBS and maintained for 24 hours. Then, supernatant was harvested and centrifuged at 100,000g for 3 hours at 4°C to concentrate retrovirus virion. Isolated virus virion was stored at −80°C. To infect retrovirus into the chick embryo, virus virion was sprinkled on the blastderm cells at St. 8 M/O chick embryo and incubated for 5 days. Fertilized M/O chicken eggs were provided from the National BioResource Project (NBRP) “Chicken/Quail” in Nagoya University.

### Thalidomide treatment with chicken embryos

A solid crystal of thalidomide (Tokyo Chemical Industry Co., Ltd) was resolved in 45% 2-hydroxypropyl-beta-cyclodextrin (HBC, FUJIFILM Wako Pure Chemical) for 1-2 h at 60°C to make 2 μg/μl thalidomide or 5-hydroxythalidomide stock solution. This stock was mixed with same volume of 2× Hanks buffer as a working solution (1 μg/μl). To apply thalidomide to an embryo, a small hole was opened at amnion above a right forelimb bud at Hamburger and Hamilton (HH) st. 18, and working solution was injected in a space between amnion and a right forelimb bud. In the case of samples for *in situ* hybridization, immunofluorescence, and immunoblot analysis, embryos were treated by 30 μl of working solution, and incubated until HH st. 22/23. In the case of samples of skeletal pattern analysis, embryos were treated by 10 μl of working solution 3 times totally every 12 h for reducing lethality, and they were incubated until approximately HH st. 36 (embryonic day 10).

### *In situ* hybridization

Embryos for whole-mount *in situ* hybridization and fluorescent in situ hybridization in sections (FISH) were fixed by 4% paraformaldehyde (PFA)/phosphate buffered saline (PBS) at 4°C for 12 h. Digoxigenin (DIG)-labeled RNA probes were prepared according to the manufacturer’s instructions (Roche). Whole-mount *in situ* hybridization was performed as previously described^45^. In FISH, 10 μm thick-frozen sections were prepared with cryostat (LEICA CM3050S). FISH protocol was described previously^46^, with additional processes in order to amplify fluorescent signal. Additional processes are as follows. After the reaction between DIG-labeled RNA and anti-DIG antibody conjugated horseradish peroxidase (HRP) (Roche), sections were washed 3 times by TNT buffer (100 mM Tris-HCl, pH7.5, 150 mM NaCl, 0.05% Tween20), and were treated with DIG amplification working solution in TSA plus DIG Kit (PerkinElmer) for 5 minutes at room temperature (RT) in accordance with manufacturer’s instructions (PerkinElmer). After 3 times wash by TNT, samples were treated by anti-DIG antibody conjugated HRP at 4°C for 12 h again.

### Immunofluorescence staining

Chicken embryos for Immunofluorescence staining were fixed by 4% PFA/PBS at 4°C for 12 h. 10μm thick-frozen sections were prepared with cryostat. PLZF immunofluorescence staining was performed in accordance with Saito et al.^47^, using anti-PLZF rabbit pAb (1:250; GeneTex, GTX111046) as primary antibodies, and anti-rabbit conjugated HRP (1:500; GE Healthcare) as secondary antibodies. To detect fluorescent signals, we used the TSA Plus Fluorescent System (PerkinElmer) for 5 min at room temperature. The sections were stained with DAPI (Wako Pure Chemical Corporation) and finally sealed by FluorSave (Calbiochem).

SALL4 was detected as follows. After pre-blocking with blocking solution (1% blocking reagent in TNT) for 1 h at RT, the sections were incubated at 4°C overnight using an anti-SALL4 rabbit pAb (1:250; Abcam, ab29112). After three washes in TNT, the specimens were reacted with anti-rabbit IgG-Alexa Fluor 488-conjugated antibody (donkey, Invitrogen) diluted 1:500 with blocking solution for 1 h at RT, followed by washing three times in TNT. The sections were stained with DAPI and sealed with FluorSave reagent.

### Limb skeletal staining

Embryos for skeletal staining (alcian or Victoria blue staining) were fixed by 10% formalin/tyrode. Alcian blue staining protocol was described previously^48^. Embryos were harvested in PBS and fixed in 10% formalin solution overnight. Fixed embryos were washed with 3%HCl/70%EtOH solution for 3 times and stained in 1% Victoria blue B (Sigma) dissolved in 3%HCl/70%EtOH overnight. Embryos were washed with 3%HCl/70%EtOH solution overnight, then they were treated in methylsalicylate for transparent process.

### Imaging

Images of FISH and immunofluorescence staining on sections were obtained using a TCS SP5 confocal microscope (LEICA). Images of whole-mount *in situ* hybridization were obtained using a fluorescent stereo microscope (LEICA M165FC with Olympus DP74 camera). Images of skeletal staining were obtained using a stereo microscope (Olympus SZX16 with Olympus DP21 camera).

### Statistical analysis

Significant changes were analysed by a one-way or two-way ANOVA followed Tukey’s post-hoc test using Graph Pad Prism 8 software (GraphPad, Inc.). For all tests, a *P* value of less than 0.05 was considered statistically significant.

### Data availability

The authors declare that all data supporting the findings of this study are available in the manuscript and its supplementary files or are available from the corresponding author upon reasonable request.

## Acknowledgements

We thank H. Yamakawa (Kazusa DNA Research Institute) for the vector construction for the human protein array, C. Takahashi and C. Furukawa for technical assistance, and the Applied Protein Research Laboratory of Ehime University. We also thank Prof. F. Tokunaga and Dr. D. Oikawa (Osaka City University) for providing the DLBCL cell lines. This work was mainly supported by the Platform Project for Supporting Drug Discovery and Life Science Research (Basis for Supporting Innovative Drug Discovery and Life Science Research (BINDS)) from AMED under Grant Number JP19am0101077 (H. Takeda, T.S.), a Grant-in-Aid for Scientific Research on Innovative Areas (JP16H06579 for T.S.) from the Japan Society for the Promotion of Science (JSPS). This work was also partially supported by JSPS KAKENHI (JP17J08477 for S.Y., JP16H04729, JP19H03218 for T.S., JP17H06112 for N.S., and JP18H02446, JP18H04811, JP18H04756 for K.T), a Grant-in-Aid for JSPS Research Fellow (JP17J08477 for S.Y) from JSPS., and Takeda Science Foundation. Establishment and screening of the human protein array used in this study was conducted in Proteo-Science Center at Ehime University with the support of MEXT.

## Author Contributions

S.Y. performed the biochemical, molecular, and cellular biology experiments. T.I. and H.Takeda constructed the human protein array. H.Takahashi supported the screening. E.T. and N.S. synthesized and analysed the 5-hydroxythalidomide. H.M., D.S., G.A., T.Suzuki, and K.T. performed the chicken teratogenesis studies. S.Y. and T.Sawasaki analysed the data, designed the study, wrote the paper, and all authors contributed to the manuscript.

## Additional information

**Supplementary Information** accompanies this paper at XXXXXX.

### Competing interests

The authors declare no competing financial interests.

**Reprints and permission** information is available online at XXXXXXX.

