## Supplementary Information for "PLZF is a new substrate of CRBN with thalidomide and 5-hydroxythalidomide"

*[1]Division of Cell-Free Sciences, Proteo-Science Center, Ehime University, Matsuyama, 790-8577 Japan, [2]Department of Ecological Developmental Adaptability Life Sciences, Graduate School of Life Sciences, Tohoku University, Sendai, 980-8578, Japan, [3]Department of Nanopharmaceutical Sciences, Nagoya Institute of Technology, Nagoya, 466-8555, Japan, [4]Division of Proteo-Drug-Discovery Sciences, Proteo-Science Center, Ehime University, Matsuyama, 790-8577 Japan. [5]Avian Bioscience Research Center, Graduate School of Bioagricultural Sciences, Nagoya University, Nagoya, 464-8601, Japan.*

*#Present address.*

*Department of Biology, Faculty of Sciences, Kyushu University, Fukuoka, 819-0395 Japan.*

*Corresponding Author:

Tatsuya Sawasaki

Proteo-Science Center,

Ehime University, Matsuyama 790-8577, Japan

### Supplementary Figures

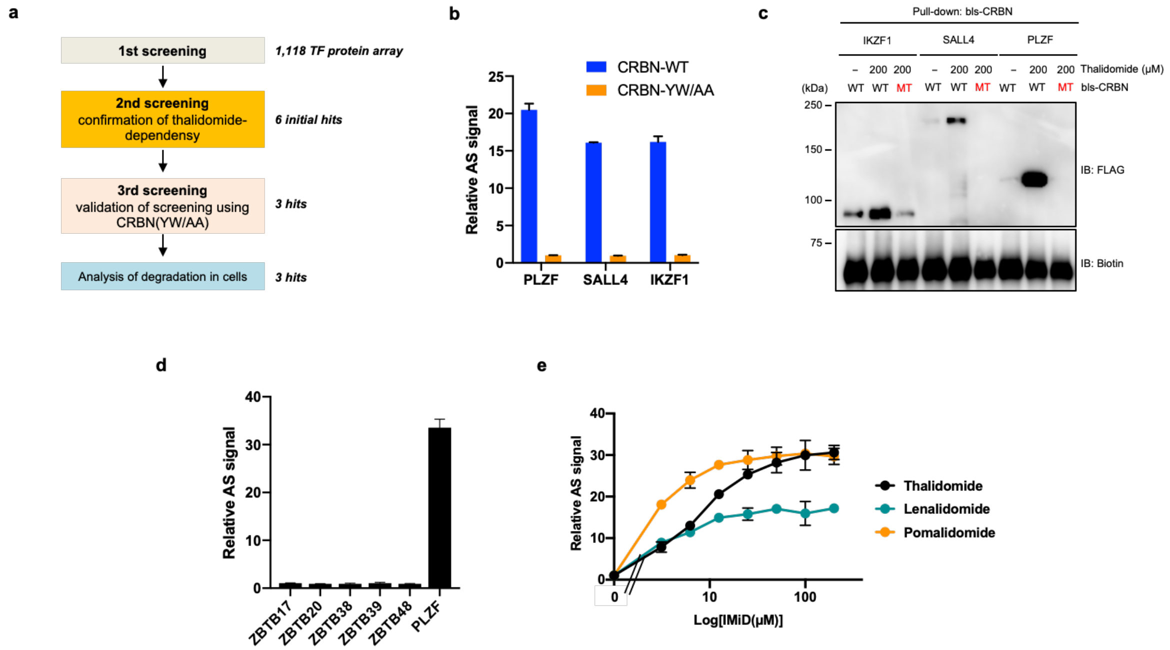


**Supplementary Figure 1. Flowchart of *in vitro* high-throughput screening and validation assay of candidate clones**

**a**, Flowchart of *in vitro* high-throughput screening. **b**, Validation of screening using CRBN mutant. Interaction between FLAG-GST-SALL4, -PLZF, or IKZF1 and bls-CRB-WT or -CRBN-YW/AA in the presence of DMSO or 50 µM thalidomide was analysed by *in vitro* binding assay using AlphaScreen technology. **c**, *In vitro* binding assay using pull-down and immunoblot analysis. bls-CRBN-WT or -CRBN-YW/AA was used as bait protein, and thalidomide-dependent interactions between bls-CRBN and FLAG-GST-SALL4, PLZF, and IKZF were confirmed by procedures described in Methods. **d**, *In vitro* binding assay for ZBTB family proteins using AlphaScreen technology. FLAG-GST-ZBTB proteins (ZBTB17, ZBTB20, ZBTB38, ZBTB39, ZBTB48, and PLZF) were evaluated for thalidomide-dependent interactions with bls-CRBN by same procedures indicated in Supplementary Fig. 1b. **e**, *In vitro* binding assay for thalidomide, pomalidomide, and lenalidomide. Interaction between bls-CRBN and FLAG-GST-SALL4 in the presence of DMSO, (3.125, 6.25, 12.5, 25, 50, 100, or 200 µM) thalidomide, pomalidomide, or lenalidomide was analysed using AlphaScreen technology. All relative AS (AlphaScreen) signals were expressed as relative luminescent signal with luminescent signal of DMSO as one, and error bars mean ± standard deviation (n = 3).

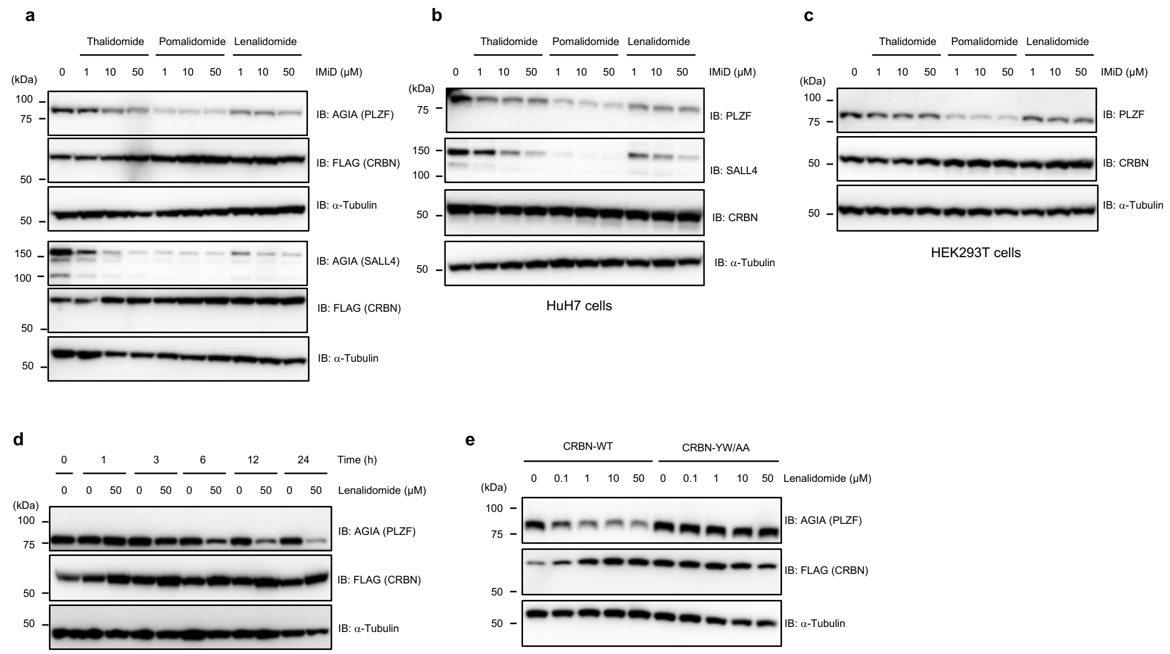


**Supplementary Figure 2. Destabilization of PLZF in IMiD-treated cells**

**a**, Immunoblot analysis of AGIA-PLZF or AGIA-SALL4 protein levels in FLAG-CRBN expressing HEK293T cells treated with DMSO, thalidomide, pomalidomide, or lenalidomide for 16 h. **b**, Immunoblot analysis of endogenous PLZF or SALL4 protein levels in HuH7 cells treated with DMSO, thalidomide, pomalidomide or lenalidomide for 24 h. **c**, Immunoblot analysis of endogenous PLZF protein levels in HEK293T cells treated with DMSO, thalidomide, pomalidomide or lenalidomide for 24 h. **d**, Time course of DMSO or lenalidomide treatment in AGIA-PLZF and FLAG-CRBN expressing HEK293T cells. AGIA-PLZF protein levels were detected by immunoblot analysis. **e**, Immunoblot analysis of AGIA-PLZF protein levels in FLAG-CRBN-WT or FLAG-CRBN-YW expressing CRBN$^{-/-}$ HEK293T cells treated with DMSO or lenalidomide for 16 h.

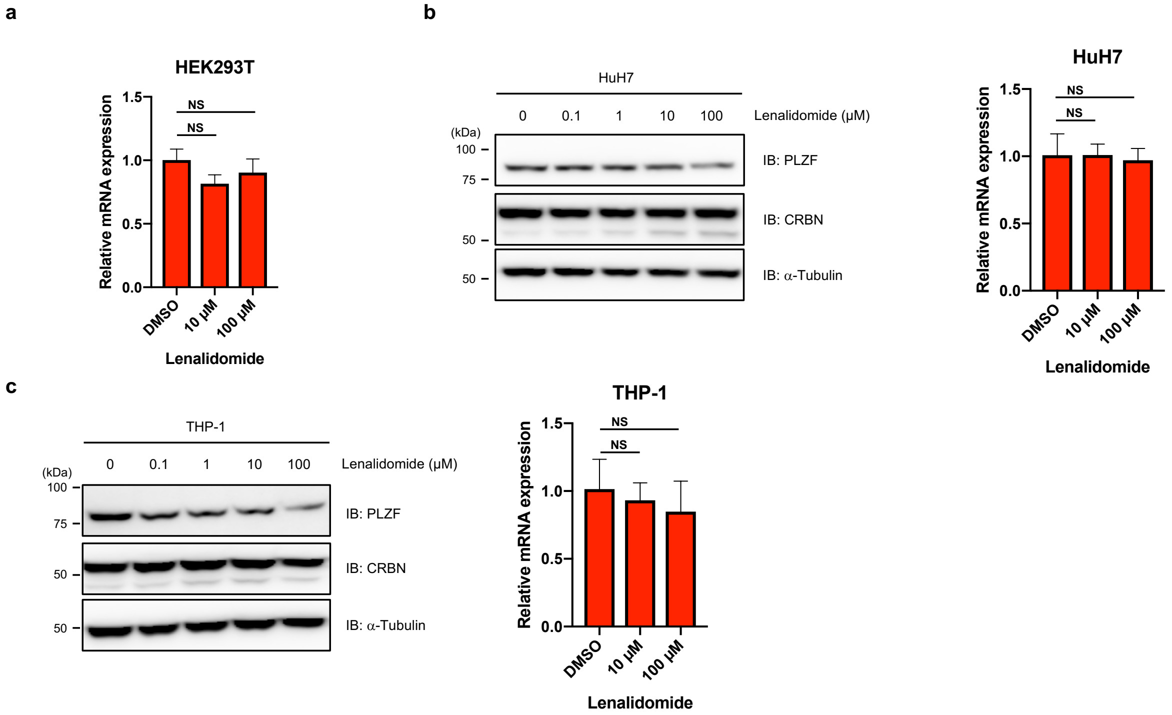


**Supplementary Figure 3. Analyses of expression of *PLZF* mRNA and degradation of endogenous PLZF in thalidomide-treated cells**

**a**, HEK293T cells were treated with the indicated concentrations of lenalidomide for 24 h and *PLZF* mRNA expression levels were measured by quantitative RT-PCR. **b**, HuH7 cells were treated with the indicated concentrations of lenalidomide for 24 h. PLZF protein levels were analysed by immunoblot and *PLZF* mRNA expression levels were measured by quantitative RT-PCR. **c**, THP-1 cells were treated with the indicated concentrations of lenalidomide for 24 h. PLZF protein levels were analysed by immunoblot and *PLZF* mRNA expression levels were measured by quantitative RT-PCR. Relative mRNA expression used the expression level with DMSO treatment as one. Error bars mean ± standard deviation (n = 3) and *P* values were calculated by one-way ANOVA with Tukey's post-hoc test (NS = Not Significant).

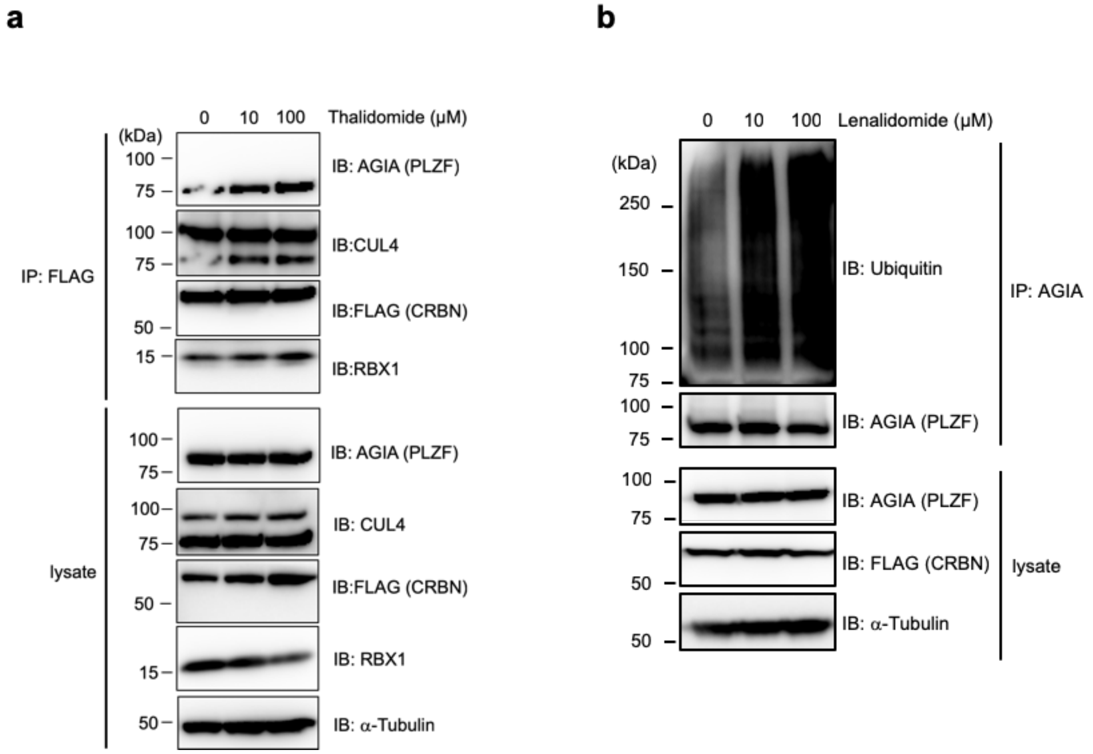


**Supplementary Figure 4. PLZF is a substrate of CRL4$^{CRBN}$ with thalidomide and lenalidomide for E3 ubiquitin ligase.**

**a**, Immunoprecipitation of FLAG-CRBN in FLAG-CRBN and AGIA-PLZF expressing CRBN$^{-/-}$ HEK293T cells treated with DMSO or thalidomide in the presence of DMSO or MG132 for 8 h. Components of CRL$^{FLAG-CRBN}$ and AGIA-PLZF were detected using each specific antibody, as indicated. **b**, Ubiquitination of AGIA-PLZF in AGIA-PLZF and FLAG-CRBN expressing HEK293T cells treated with DMSO or lenalidomide in the presence of DMSO or MG132 for 10 h.

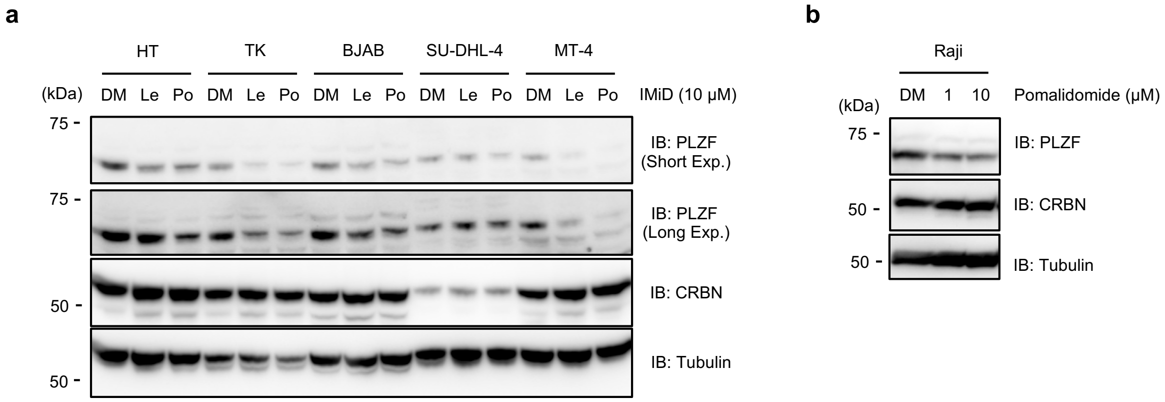


**Supplementary Figure 5. IMiD-induced protein degradation of PLZF in B cell lymphomas.**

**a**, HT, TK, BJAB, SY-DHL-4, and MT-4 cells were treated with DMSO, 10 µM lenalidomide, or pomalidomide for 24 h. PLZF protein levels were analysed by immunoblot. **b**, Raji cells were treated with DMSO, 10 µM lenalidomide, or pomalidomide for 24 h. PLZF protein levels were analysed by immunoblot.

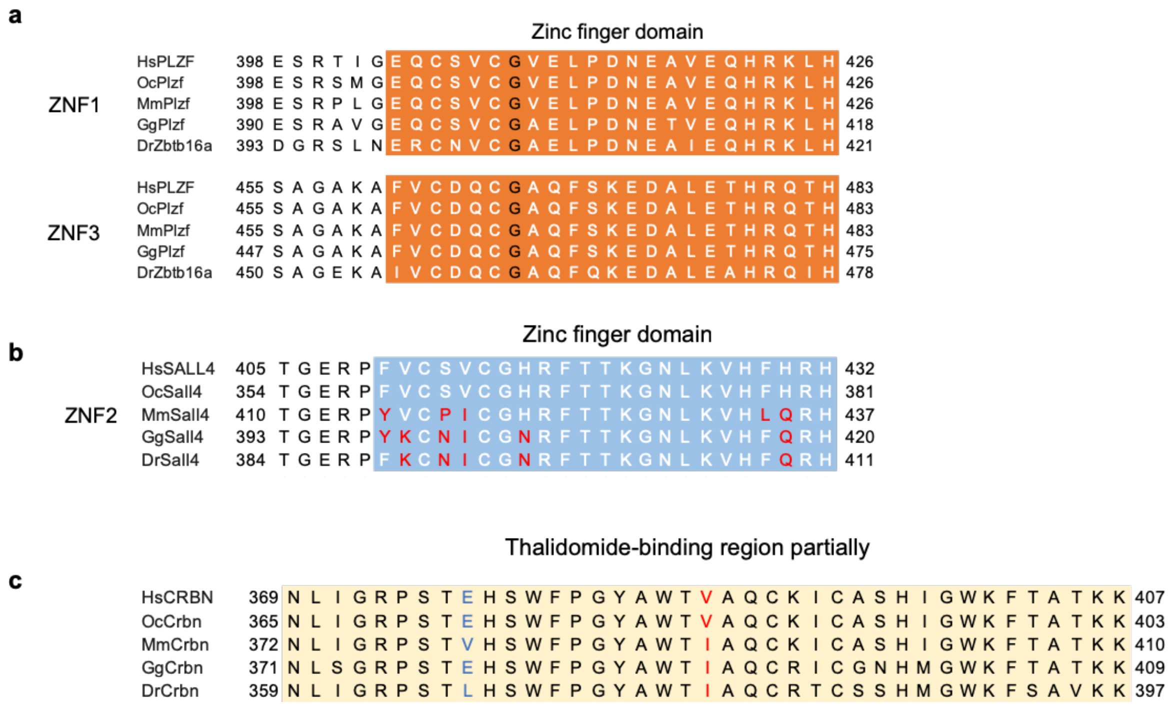


**Supplementary Figure 6. Sequence comparisons of thalidomide-related regions in vertebrate PLZF, SALL4 and CRBN.**

**a**, Alignment of amino acid sequence of ZNF1 and ZNF3 in PLZF among human (Hs), rabbit (Oc), mouse (Mm), chicken (Gg), and zebrafish (Dr). **b**, Alignment of amino acid sequence of ZNF2 in SALL4 among the species above. **c**, Alignment of amino acid sequence in CRBN among the species above.

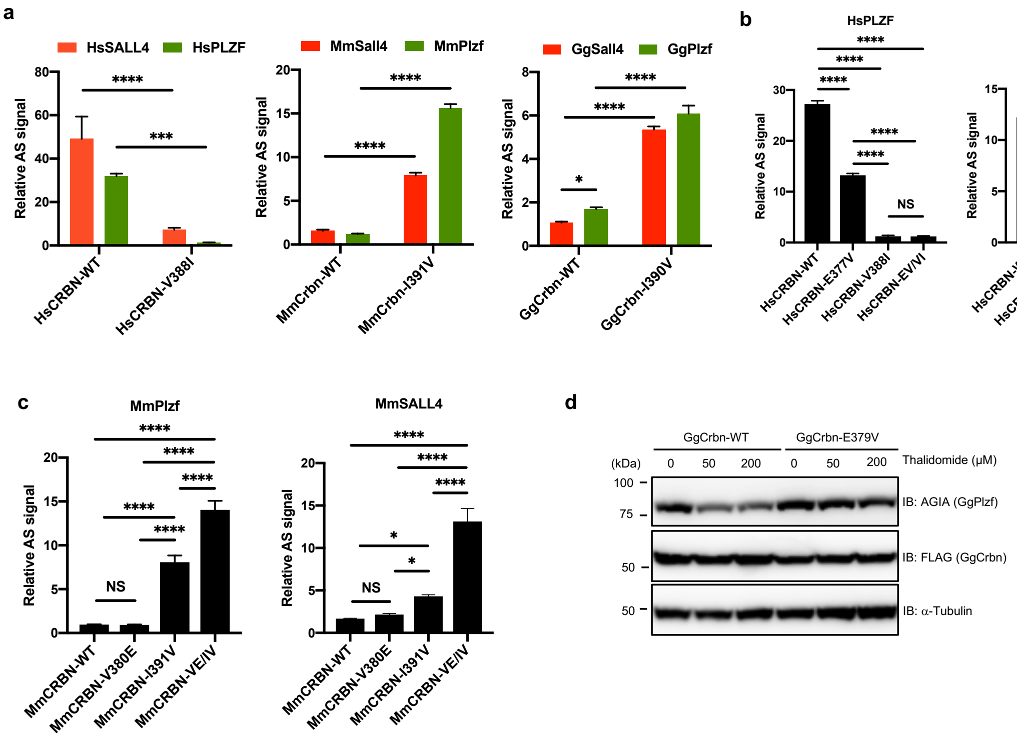


**Supplementary Figure 7. Interaction and protein degradation analyses between PLZF or SALL4 and CRBN with thalidomide among human, mouse and chicken.**

**a**, *In vitro* binding assay using human, mouse, and chicken proteins. Thalidomide-dependent interaction between biotinylated-HsCRBN, -MmCrbn or -GgCrbn, and FLAG-GST-SALL4, or -PLZF (Hs, Mm or Gg) in the presence of DMSO or 200 µM thalidomide was analysed using AlphaScreen technology. **b-c**, *In vitro* binding assay using human or mouse proteins. Thalidomide-dependent interaction in the presence of DMSO or 50 µM thalidomide was analysed using same procedure in Supplementary Fig 7a. **d**, Immunoblot analysis of AGIA-GgPlzf in FLAG-GgCrbn-WT or -E379V expressing $CRBN^{-/-}$ HEK293T cells treated with DMSO, 50 µM or 200 µM thalidomide for 16 h. All relative AS (AlphaScreen) signals are expressed as relative luminescent signal with luminescent signal of DMSO as one. Error bars mean ± standard deviation (n = 3) and *P* values were calculated by one-way or two-way ANOVA with Tukey's post-hoc test (NS = Not Significant, $^{*}P < 0.05$, $^{***}P < 0.001$, and $^{****}P < 0.0001$).

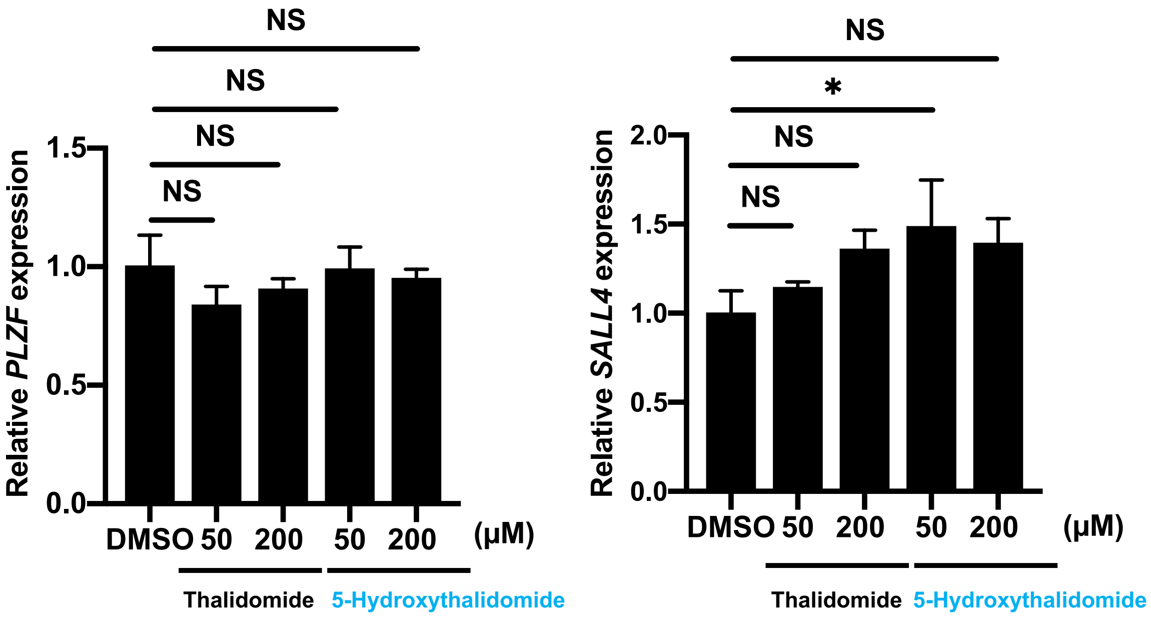


**Supplementary Figure 8. Effect of thalidomide or 5-hydroxythalidomide on mRNA expression of PLZF or SALL4 in HuH7 cells.**

HuH7 cells were treated with the indicated concentrations of DMSO, thalidomide or 5-hydroxythalidomide for 24 h. *PLZF or SALL4* mRNA expression levels were measured by quantitative RT-PCR. Relative mRNA expression used the expression level with DMSO treatment as one. Error bars mean ± standard deviation (n = 3) and *P* values were calculated by one-way ANOVA with Tukey's post-hoc test (NS = Not Significant, and $^{*}P < 0.05$).

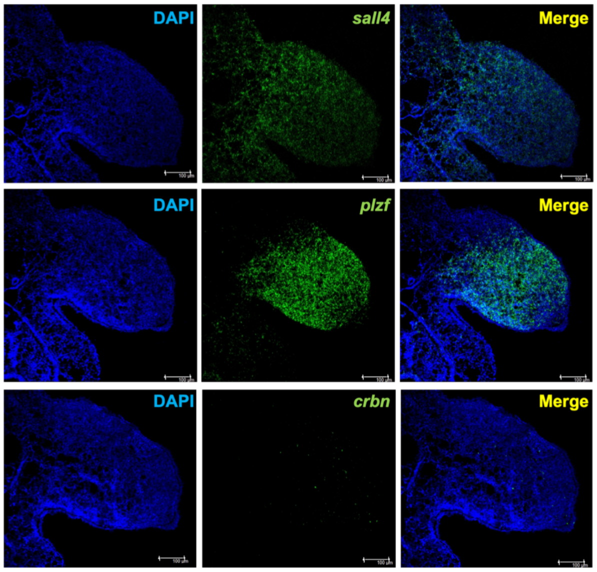


Scale bar = 100 μm

**Supplementary Figure 9. Analysis of *Crbn, Sall4*, and *Plzf* mRNAs in chicken embryos**
*Sall4*, *Plzf,* or *Crbn* mRNA expression in E4 chicken limb bud was analysed by section *in situ* hybridization.

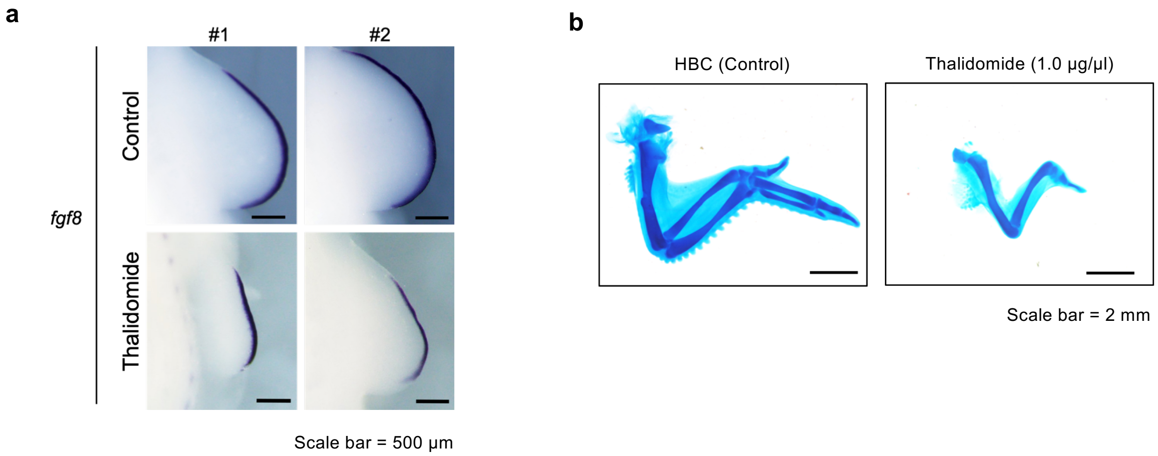


**Supplementary Figure 10. Phenotypes of thalidomide treatment in chicken embryo.**
**a**, *fgf8* mRNA expression in E4 chicken right limb bud embryo was analysed by whole-mount *in situ* hybridization. The two independent samples of control (DMSO, n = 4, upper panel) and thalidomide-treated (1 μg/μl, n = 7, lower panel) chicken embryos are shown. **b,** Limb skeletal stained with Alcian blue. Skeletal patterning of right forelimb in E10 chicken embryos treated with HBC (control, n = 6) or 1 μg/μl thalidomide (n = 6) were analysed by Alcian blue staining.

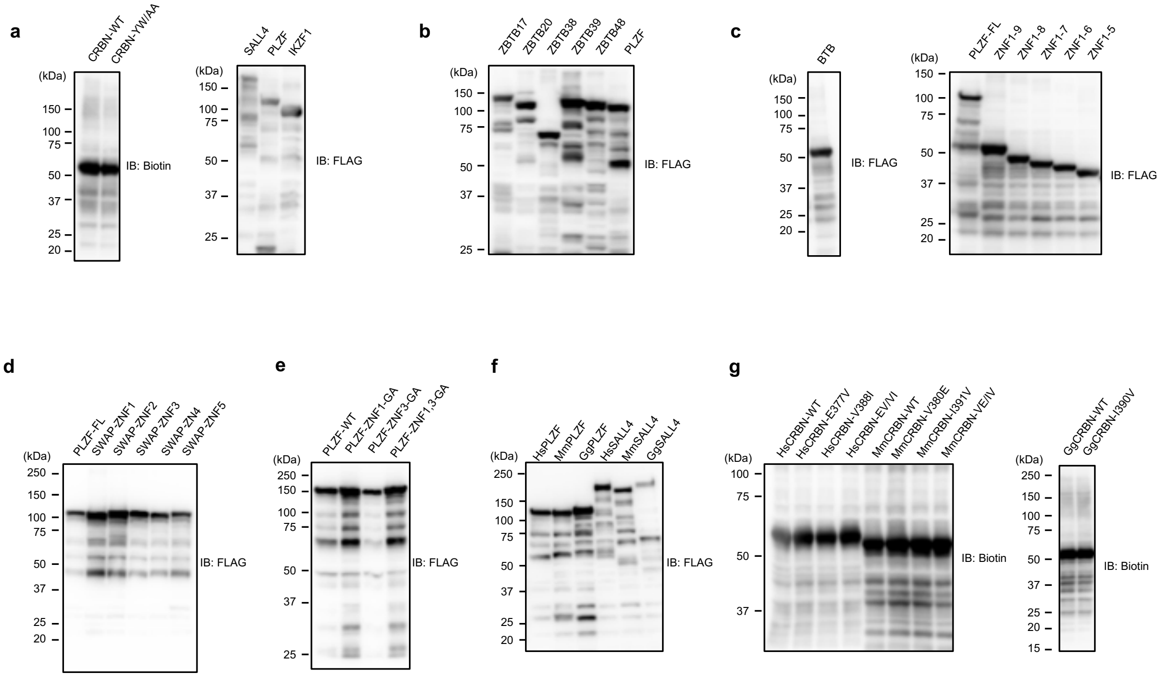


**Supplementary Figure 11. Recombinant proteins by a wheat cell-free system.**

**a-g**, Confirmation of recombinant proteins synthesized using a wheat cell-free system by Immunoblot analysis.

List of proteins in hTF Protein Array

| Entrez Gene Symbol | Kazusa Original clone ID | NCBI RefSeq top hit | AlphaScreen sign | Idenitity to RefSeq sequence |
|---|---|---|---|---|
| AATF | FXC01868,FHC01868,ORS01868,ORH01868 | NM_012138.3 | 318 | Complete match |
| ABRA | FXC02778,FHC02778,ORH02778 | NM_139166.4 | 379 | Complete match |
| ADGRG3 | FHC12673 | NM_170776.4 | 544 | Complete match |
| AEBP1 | FXC01470,FHC01470,ORK01470 | NM_001129.4 | 437 | Complete match |
| AEBP2 | FHC30523 | NM_001114176.1 | 392 | Complete match |
| AFF4 | FHC07632 | NM_014423.3 | 363 | Complete match |
| AGER | FXC03485,FHC03485 | NM_001136.4 | 478 | Complete match |
| AGT | FXC11578,FHC11578 | NM_000029.3 | 486 | Complete match |
| AHR | FXC01249,FHC01249 | NM_001621.4 | 395 | Complete match |
| AIRE | ORH16149P | NM_000383.4 | 441 | Partial match |
| AKT1 | FXC02215,FHC02215,ORK02215,ORH02215P | NM_005163.2 | 542 | Complete match |
| ALK | FXC00337,FHC00337,ORK00337 | NM_004304.4 | 617 | Complete match |
| ALX1 | FHC24829 | NM_006982.2 | 716 | Complete match |
| ALX4 | FXC01180,FHC01180,ORK01180 | NM_021926.3 | 1087 | Complete match |
| ANKRD30A | FHC23819,ORS23819P | NM_052997.2 | 383 | Complete match |
| ANXA3 | FHC21966 | NM_005139.2 | 314 | Complete match |
| AR | FXC11031,FHC11031 | NM_000044.3 | 436 | Complete match |
| ARHGEF2 | ORS09014 | NM_004723.3 | 495 | Complete match |
| ARHGEF5 | FXC01043,FHC01043,ORK01043,ORS01043,ORH01043 | NM_005435.3 | 363 | Complete match |
| ARID3A | FHC11107 | NM_005224.2 | 519 | Complete match |
| ARID3B | ORH14423P | NM_006465.3 | 260 | Partial match |
| ARID4A | FHC12701 | NM_023001.2 | 384 | Complete match |
| ARID5B | FXC11014,FHC11014,ORH11014 | NM_032199.2 | 469 | Complete match |
| ARNT | FXC01533,FHC01533 | NM_001286036.1 | 445 | Complete match |
| ARNT2 | ORH12033 | NM_014862.3 | 454 | Complete match |
| ARNTL | FXC03462,FHC03462 | NM_001297719.1 | 567 | Complete match |
| ARNTL2 | FHC12305,ORH12305 | NM_001248004.1 | 531 | Complete match |
| ARRB1 | FXC02162,FHC02162,ORH02162P | NM_004041.4 | 424 | Complete match |
| ARRB2 | FXC03515,FHC03515 | NM_004313.3 | 611 | Complete match |
| ASCL1 | FXC03176,FHC03176 | NM_004316.3 | 444 | Complete match |
| ASCL2 | FHC24115 | NM_005170.2 | 568 | Complete match |
| ASCL3 | FXC03178,FHC03178 | NM_020646.2 | 667 | Complete match |
| ATF1 | FHC10950,ORH10950 | NM_005171.4 | 1253 | Complete match |
| ATF2 | FXC01751,FHC01751,ORK01751,ORH01751P | NM_001880.3 | 448 | Complete match |
| ATF3 | FXC08278,FHC08278,ORS08278,ORH08278 | NM_001674.3 | 506 | Complete match |
| ATF4 | FXC04200,FHC04200,ORH04200 | NM_001675.4 | 351 | Complete match |
| ATF5 | FXC02605,FHC02605 | NM_012068.5 | 684 | Complete match |
| ATF6B | FXC03600,FHC03600 | NM_004381.4 | 437 | Complete match |
| ATF7 | FXC03273,FHC03273 | NM_006856.2 | 682 | Complete match |
| ATOH1 | FXC03179,FHC03179,ORH03179 | NM_005172.1 | 515 | Complete match |

| | | | | |
|---|---|---|---|---|
| ATOH8 | FXC02368,FHC02368,ORK02368 | NM_032827.6 | 433 | Complete match |
| BACH1 | FXC01715,FHC01715,ORH01715P | NM_001186.3 | 276 | Complete match |
| BACH2 | FXC10469,FHC10469 | NM_021813.3 | 341 | Complete match |
| BARHL1 | FXC03258,FHC03258 | NM_020064.3 | 1129 | Complete match |
| BARHL2 | FHC20405 | NM_020063.1 | 600 | Complete match |
| BARX1 | FHC14877,ORH14877P | NM_021570.4 | 437 | Complete match |
| BARX2 | FXC03756,FHC03756 | NM_003658.4 | 584 | Complete match |
| BATF | FXC02486,FHC02486 | NM_006399.3 | 279 | Complete match |
| BATF2 | FXC02519,FHC02519 | NM_138456.3 | 518 | Complete match |
| BCL10 | FXC03407,FHC03407,ORH03407P | NM_003921.4 | 969 | Complete match |
| BCL11A | FXC00923,FHC00923,ORK00923 | NM_018014.3 | 376 | Complete match |
| BCL11B | FHC25293 | NM_138576.3 | 441 | Complete match |
| BCL6 | FXC01416,FHC01416,ORH01416 | NM_001706.4 | 779 | Complete match |
| BCL6B | FXC02560,FHC02560,ORH02560P | NM_181844.3 | 351 | Complete match |
| BEX1 | ORH29461P | NM_018476.3 | 740 | Complete match |
| BHLHA15 | FXC03298,FHC03298,ORH03298P | NM_177455.3 | 367 | Complete match |
| BHLHE40 | FXC02724,FHC02724 | NM_003670.2 | 515 | Complete match |
| BLZF1 | FXC02588,FHC02588,ORK02588,ORS02588,ORH02588 | NM_003666.2 | 474 | Complete match |
| BMP7 | FXC08377,FHC08377,ORS08377,ORH08377 | NM_001719.2 | 637 | Complete match |
| BMPR1A | FXC09257,FHC09257,ORS09257,ORH09257 | NM_004329.2 | 580 | Complete match |
| BRMS1 | FHC24321,ORH24321P | NM_015399.3 | 331 | Complete match |
| BTBD8 | ORH31344 | NM_183242.3 | 425 | Complete match |
| BTG2 | FXC04297,FHC04297,ORH04297 | NM_006763.2 | 404 | Complete match |
| BTK | FHC05404,ORH05404P | NM_000061.2 | 441 | Complete match |
| BTRC | FXC11937,FHC11937,ORH11937P | NM_033637.3 | 701 | Complete match |
| CAMTA1 | FXC00650,FHC00650,ORK00650 | NM_015215.3 | 384 | Complete match |
| CAMTA2 | FXC01135,FHC01135,ORK01135 | NM_015099.3 | 511 | Complete match |
| CAPN3 | FHC25360,ORH25360P | NM_173088.1 | 395 | Complete match |
| CARD14 | FXC26210,FHC26210 | NM_024110.4 | 679 | Complete match |
| CBFA2T2 | FXC03455,FHC03455,ORH03455P | NM_005093.3 | 412 | Partial match |
| CBFA2T3 | FXC03597,FHC03597 | NM_005187.5 | 428 | Complete match |
| CBFB | FXC02508,FHC02508,ORS02508,ORH02508 | NM_001755.2 | 317 | Complete match |
| CBL | FHC12662 | NM_005188.3 | 486 | Complete match |
| CC2D1B | ORH13286P | NM_032449.2 | 351 | Complete match |
| CDX1 | FXC03255,FHC03255 | NM_001804.2 | 527 | Complete match |
| CDX2 | FXC02540,FHC02540,ORS02540,ORH02540 | NM_001265.4 | 350 | Complete match |
| CDX4 | FHC27424,ORH27424P | NM_005193.1 | 954 | Complete match |
| CEBPA | FXC10825,FHC10825 | NM_004364.4 | 473 | Complete match |
| CEBPB | FXC03181,FHC03181 | NM_005194.3 | 470 | Complete match |
| CEBPD | FHC23262 | NM_005195.3 | 945 | Complete match |
| CEBPE | FXC06663,FHC06663,ORH06663 | NM_001805.3 | 515 | Complete match |
| CEBPG | FXC03182,FHC03182,ORS03182,ORH03182 | NM_001806.3 | 440 | Complete match |

| | | | | |
|---|---|---|---|---|
| CFLAR | FHC09775,ORS09775 | NM_003879.5 | 468 | Complete match |
| CGGBP1 | ORH13213P | NM_003663.3 | 711 | Complete match |
| CHCHD3 | FHC12154,ORH12154P | NM_017812.2 | 335 | Complete match |
| CHP1 | FHC11145 | NM_007236.4 | 308 | Complete match |
| CHUK | FHC10858 | NM_001278.3 | 466 | Complete match |
| CIB1 | FXC02705,FHC02705,ORS02705,ORH02705 | NM_006384.3 | 310 | Complete match |
| CIR1 | FXC03939,FHC03939 | NM_004882.3 | 367 | Complete match |
| CITED1 | FXC03083,FHC03083,ORS03083,ORH03083 | NM_004143.3 | 347 | Complete match |
| CLOCK | FXC00509,FHC00509,ORK00509,ORH00509P | NM_004898.3 | 436 | Complete match |
| CLU | ORS08502,ORH08502 | NM_001831.3 | 386 | Complete match |
| CMKLR1 | FXC02219,FHC02219,ORH02219P | NM_001142343.1 | 511 | Complete match |
| CNBP | FXC02504,FHC02504 | NM_003418.4 | 502 | Complete match |
| CNOT7 | FXC03415,FHC03415,ORH03415P | NM_013354.5 | 444 | Complete match |
| CNOT8 | FXC02751,FHC02751,ORH02751 | NM_004779.5 | 429 | Complete match |
| COMMD1 | FHC21017,ORH21017P | NM_152516.2 | 321 | Complete match |
| COMMD6 | ORH25026P | NM_203495.3 | 305 | Complete match |
| CREB1 | FXC00339,FHC00339,ORK00339,ORS00339,ORH00339 | NM_134442.3 | 380 | Complete match |
| CREB3 | FXC02549,FHC02549,ORH02549P | NM_006368.4 | 592 | Complete match |
| CREB3L1 | FXC04624,FHC04624,ORH04624P | NM_052854.3 | 470 | Complete match |
| CREB3L2 | FXC04625,FHC04625,ORK04625 | NM_194071.3 | 506 | Complete match |
| CREB3L3 | FHC26402 | NM_032607.2 | 363 | Complete match |
| CREB3L4 | FXC03678,FHC03678,ORH03678P | NM_130898.3 | 576 | Complete match |
| CREB5 | ORH14419P | NM_182898.4 | 302 | Partial match |
| CREBBP | FXC11778,FHC11778,ORK11778 | NM_004380.2 | 302 | Complete match |
| CREBL2 | FXC03296,FHC03296,ORK03296 | NM_001310.3 | 204 | Complete match |
| CREBRF | FXC06188,FHC06188,ORH06188 | NM_153607.2 | 412 | Complete match |
| CREM | FHC10995 | NM_181571.2 | 614 | Complete match |
| CRHR2 | FHC22891 | NM_001883.4 | 589 | Complete match |
| CRTC1 | FXC03691,FHC03691 | NM_015321.2 | 457 | Complete match |
| CRTC2 | FHC27707,ORH27707 | NM_181715.2 | 617 | Complete match |
| CRTC3 | ORH16133P | NM_022769.5 | 310 | Partial match |
| CRX | FXC11684,FHC11684 | NM_000554.4 | 968 | Complete match |
| CSRNP1 | FHC12609 | NM_033027.3 | 424 | Complete match |
| CSRNP2 | FHC24725,ORH24725P | NM_030809.2 | 363 | Complete match |
| CSRNP3 | FHC12588 | NM_001172173.2 | 375 | Partial match |
| CTBP1 | FXC03687,FHC03687 | NM_001328.2 | 416 | Complete match |
| CTCF | FXC01807,FHC01807,ORS01807,ORH01807 | NM_006565.3 | 485 | Complete match |
| CTCFL | FHC27078,ORH27078P | NM_080618.3 | 342 | Complete match |
| CTH | ORH13248P | NM_001902.5 | 379 | Complete match |
| CTNNB1 | FXC07714,FHC07714 | NM_001904.3 | 400 | Complete match |
| CUX1 | FXC03290,FHC03290,ORK03290 | NM_001913.4 | 293 | Complete match |
| CUX2 | FXC00499,FHC00499,ORK00499 | NM_015267.3 | 451 | Complete match |
| CYP1B1 | FHC20974,ORH20974P | NM_000104.3 | 486 | Complete match |

| | | | | |
|---|---|---|---|---|
| CYTL1 | ORS10263,ORH10263 | NM_018659.2 | 421 | Complete match |
| DAB2IP | FHC00908,ORK00908,ORH00908 | NM_032552.3 | 396 | Partial match |
| DACH1 | FHC11768 | NM_080759.5 | 367 | Complete match |
| DAP | FHC10855,ORH10855P | NM_004394.2 | 403 | Complete match |
| DDIT3 | FXC06075,FHC06075,ORH06075P | NM_004083.5 | 321 | Complete match |
| DDN | FXC04724,FHC04724,ORK04724 | NM_015086.2 | 457 | Partial match |
| DDRGK1 | FHC26885,ORH26885P | NM_023935.1 | 498 | Complete match |
| DDX58 | FXC01556,FHC01556 | NM_014314.3 | 548 | Complete match |
| DEAF1 | FXC07762,FHC07762,ORK07762 | NM_021008.3 | 802 | Partial match |
| DLX2 | FHC11120 | NM_004405.3 | 773 | Complete match |
| DLX3 | FXC04791,FHC04791,ORH04791 | NM_005220.2 | 834 | Complete match |
| DLX4 | FXC01938,FHC01938,ORH01938P | NM_138281.2 | 658 | Complete match |
| DLX5 | FXC02532,FHC02532,ORH02532P | NM_005221.5 | 727 | Complete match |
| DLX6 | ORH14058P | NM_005222.4 | 630 | Partial match |
| DMBX1 | FXC03767,FHC03767 | NM_172225.1 | 387 | Complete match |
| DMRT1 | FXC02544,FHC02544,ORH02544P | NM_021951.2 | 970 | Complete match |
| DMRT2 | FXC03250,FHC03250 | NM_006557.6 | 466 | Complete match |
| DMRT3 | FHC23446 | NM_021240.3 | 662 | Complete match |
| DMRTA1 | FXC02229,FHC02229,ORH02229P | NM_022160.2 | 593 | Complete match |
| DMRTB1 | FHC20313 | NM_033067.2 | 1126 | Complete match |
| DMRTC1B | FXC03402,FHC03402 | NM_001080851.1 | 313 | Complete match |
| DMRTC2 | FXC02539,FHC02539 | NM_001040283.2 | 482 | Complete match |
| DNAJA3 | FHC04801,ORK04801,ORH04801P | NM_005147.5 | 589 | Complete match |
| DRAP1 | FXC02525,FHC02525 | NM_006442.3 | 404 | Complete match |
| DVL2 | FXC03710,FHC03710,ORH03710P | NM_004422.2 | 399 | Complete match |
| E2F2 | FXC02567,FHC02567 | NM_004091.3 | 388 | Complete match |
| E2F3 | FHC12471,ORS12471 | NM_001949.4 | 543 | Complete match |
| E2F4 | FXC09474,FHC09474,ORS09474 | NM_001950.3 | 323 | Complete match |
| E2F6 | FXC02528,FHC02528,ORH02528P | NM_198256.3 | 612 | Complete match |
| E2F7 | FHC12665 | NM_203394.2 | 478 | Complete match |
| E2F8 | FHC24174,ORH24174P | NM_024680.3 | 683 | Complete match |
| E4F1 | FHC12684 | NM_004424.4 | 490 | Complete match |
| EAF2 | FXC10016,FHC10016,ORS10016,ORH10016 | NM_018456.4 | 379 | Complete match |
| EBF1 | FXC01587,FHC01587,ORH01587P | NM_024007.3 | 380 | Complete match |
| EBF2 | FXC10735,FHC10735,ORH10735 | NM_022659.3 | 470 | Complete match |
| EBF4 | FXC00838,FHC00838,ORK00838 | NM_001110514.1 | 576 | Partial match |
| EDA | FXC02133,FHC02133 | NM_001005609.1 | 375 | Complete match |
| EDA2R | FXC03935,FHC03935,ORH03935P | NM_021783.3 | 400 | Complete match |
| EDF1 | FHC04175 | NM_001281297.1 | 268 | Complete match |
| EGLN1 | ORH15784P | NM_022051.2 | 342 | Complete match |
| EGR1 | FHC31400,ORH31400P | NM_001964.2 | 486 | Complete match |
| EGR2 | FXC02226,FHC02226 | NM_000399.3 | 559 | Complete match |
| EGR3 | FXC11738,FHC11738,ORH11738P | NM_004430.2 | 569 | Complete match |
| EGR4 | FXC03272,FHC03272 | NM_001965.4 | 671 | Partial match |
| EHF | FHC24200,ORH24200 | NM_012153.5 | 481 | Complete match |

| | | | | |
|---|---|---|---|---|
| EIF2AK2 | FXC20968,FHC20968 | NM_001135652.2 | 620 | Complete match |
| ELF1 | FXC01862,FHC01862,ORS01862,ORH01862 | NM_172373.3 | 563 | Complete match |
| ELF2 | FHC09540,ORS09540,ORH09540 | NM_006874.3 | 655 | Complete match |
| ELF3 | FXC03671,FHC03671 | NM_004433.4 | 790 | Complete match |
| ELF4 | FXC08882,FHC08882,ORS08882,ORH08882 | NM_001421.3 | 477 | Complete match |
| ELF5 | FXC09308,FHC09308,ORS09308,ORH09308 | NM_198381.1 | 1531 | Complete match |
| ELK1 | FXC11409,FHC11409 | NM_005229.4 | 1158 | Complete match |
| ELK3 | FXC02727,FHC02727,ORK02727,ORS02727,ORH02727 | NM_005230.3 | 634 | Complete match |
| ELK4 | FHC20788 | NM_001973.3 | 691 | Complete match |
| EN1 | FHC04893,ORH04893 | NM_001426.3 | 547 | Complete match |
| ENO1 | FXC02193,FHC02193,ORK02193,ORS02193,ORH02193 | NM_001428.3 | 273 | Complete match |
| EOMES | FXC03283,FHC03283 | NM_001278182.1 | 801 | Complete match |
| EP300 | FXC01787,FHC01787,ORK01787 | NM_001429.3 | 448 | Complete match |
| EPAS1 | FXC01393,FHC01393 | NM_001430.4 | 576 | Complete match |
| EPHA5 | FXC01385,FHC01385,ORK01385 | NM_001281767.1 | 236 | Complete match |
| ERBB2IP | FXC00772, FHC00772 | NM_018695.3 | 429 | Complete match |
| ERC1 | FXC00176,FHC00176,ORK00176 | NM_178040.4 | 548 | Partial match |
| ERF | FXC04909,FHC04909 | NM_006494.3 | 474 | Complete match |
| ERG | FXC04910,FHC04910,ORH04910 | NM_182918.3 | 699 | Complete match |
| ESR1 | FXC10665,FHC10665,ORH10665 | NM_000125.3 | 456 | Complete match |
| ESRRA | FXC04917,FHC04917 | NM_004451.4 | 605 | Complete match |
| ESRRB | FXC03274,FHC03274 | NM_004452.3 | 383 | Partial match |
| ESRRG | FXC00649,FHC00649,ORK00649 | NM_001438.3 | 814 | Complete match |
| ESX1 | FHC11691 | NM_153448.3 | 839 | Complete match |
| ETS1 | FXC02157,FHC02157,ORK02157 | NM_001143820.1 | 436 | Complete match |
| ETS2 | FXC10074,FHC10074,ORS10074,ORH10074 | NM_005239.5 | 637 | Complete match |
| ETV1 | FXC02117,FHC02117,ORK02117 | NM_001163147.1 | 540 | Complete match |
| ETV2 | FXC03261,FHC03261 | NM_014209.3 | 478 | Complete match |
| ETV3 | FXC03759,FHC03759 | NM_001145312.1 | 625 | Complete match |
| ETV4 | FXC04918,FHC04918,ORS04918,ORH04918 | NM_001986.2 | 662 | Complete match |
| ETV5 | FXC04919,FHC04919,ORS04919,ORH04919 | NM_004454.2 | 658 | Complete match |
| ETV6 | FXC01505,FHC01505,ORK01505,ORH01505P | NM_001987.4 | 911 | Complete match |
| ETV7 | FXC04920,FHC04920,ORH04920P | NM_016135.3 | 747 | Complete match |
| EVX1 | FHC22886,ORH22886 | NM_001989.4 | 462 | Complete match |
| EZH2 | FXC08523,FHC08523,ORH08523 | NM_004456.4 | 471 | Complete match |
| FBXW7 | FXC05007,FHC05007,ORK05007 | NM_018315.4 | 905 | Complete match |
| FER | FHC03985 | NM_005246.2 | 404 | Complete match |
| FERD3L | FXC11360,FHC11360 | NM_152898.2 | 482 | Complete match |
| FEV | FHC09121,ORS09121,ORH09121 | NM_017521.2 | 711 | Complete match |
| FIGLA | FXC02433,FHC02433,ORS02433,ORH02433 | NM_001004311.3 | 486 | Complete match |

| | | | | |
|---|---|---|---|---|
| FLI1 | FXC01531,FHC01531,ORK01531,ORS01531,ORH01531 | NM_002017.4 | 708 | Complete match |
| FOS | FXC01934,FHC01934,ORS01934,ORH01934 | NM_005252.3 | 383 | Complete match |
| FOSB | FXC05218,FHC05218 | NM_006732.2 | 269 | Complete match |
| FOSL1 | FXC02515,FHC02515,ORS02515,ORH02515 | NM_005438.4 | 339 | Complete match |
| FOSL2 | FXC09112,FHC09112,ORS09112,ORH09112 | NM_005253.3 | 379 | Complete match |
| FOXA1 | FXC02786,FHC02786,ORH02786 | NM_004496.3 | 593 | Complete match |
| FOXA2 | FXC02223,FHC02223 | NM_153675.2 | 494 | Complete match |
| FOXA3 | FXC03119,FHC03119,ORH03119P | NM_004497.2 | 604 | Complete match |
| FOXB1 | FXC11731,FHC11731 | NM_012182.2 | 855 | Complete match |
| FOXD3 | FXC03187,FHC03187 | NM_012183.2 | 368 | Complete match |
| FOXD4 | FXC03188,FHC03188,ORH03188P | NM_207305.4 | 588 | Complete match |
| FOXD4L1 | FHC21152,ORS21152,ORH21152 | NM_012184.4 | 564 | Complete match |
| FOXD4L3 | FHC30313,ORS30313P | NM_199135.4 | 445 | Complete match |
| FOXD4L4 | FHC30306,ORS30306 | NM_199244.3 | 429 | Complete match |
| FOXD4L6 | FHC30308,ORH30308P | NM_001085476.1 | 457 | Complete match |
| FOXE1 | FXC03189,FHC03189 | NM_004473.3 | 482 | Complete match |
| FOXH1 | FXC02299,FHC02299,ORS02299 | NM_003923.2 | 381 | Complete match |
| FOXI1 | FXC03128,FHC03128 | NM_012188.4 | 617 | Complete match |
| FOXJ1 | FXC02556,FHC02556,ORH02556P | NM_001454.3 | 354 | Complete match |
| FOXL1 | FXC03190,FHC03190 | NM_005250.2 | 433 | Complete match |
| FOXN2 | FXC03976,FHC03976,ORH03976P | NM_002158.3 | 748 | Complete match |
| FOXN3 | FXC03977,FHC03977,ORH03977P | NM_005197.3 | 558 | Complete match |
| FOXN4 | FXC03275,FHC03275,ORH03275P | NM_213596.2 | 715 | Complete match |
| FOXO3 | FHC22719,ORH22719 | NM_001455.3 | 507 | Complete match |
| FOXO4 | FXC05227,FHC05227 | NM_005938.3 | 1360 | Complete match |
| FOXP1 | FXC03653,FHC03653,ORH03653 | NM_032682.5 | 748 | Complete match |
| FOXP2 | FHC23055,ORH23055P | NM_014491.4 | 770 | Partial match |
| FOXP3 | FXC02319,FHC02319,ORH02319 | NM_014009.3 | 523 | Complete match |
| FOXP4 | FXC03656,FHC03656 | NM_001012426.1 | 638 | Complete match |
| FOXR1 | FHC28077,ORH28077P | NM_181721.2 | 621 | Complete match |
| FOXR2 | FXC03331,FHC03331,ORH03331P | NM_198451.3 | 310 | Complete match |
| FOXS1 | FXC03186,FHC03186,ORS03186,ORH03186 | NM_004118.3 | 592 | Complete match |
| FUBP1 | FHC10077,ORS10077,ORH10077 | NM_003902.4 | 1265 | Partial match |
| FZD2 | FHC26069 | NM_001466.3 | 550 | Complete match |
| FZD4 | FHC24400,ORH24400P | NM_012193.3 | 547 | Complete match |
| FZD6 | FHC03196 | NM_003506.3 | 371 | Complete match |
| GABPA | FXC02115,FHC02115 | NM_002040.3 | 759 | Complete match |
| GABPB1 | FXC06052,FHC06052,ORH06052P | NM_005254.5 | 327 | Complete match |
| GAS7 | FXC00064,FHC00064,ORK00064,ORH00064P | NM_001130831.1 | 535 | Complete match |
| GATA1 | FXC05275,FHC05275 | NM_002049.3 | 847 | Complete match |
| GATA3 | FXC02784,FHC02784,ORH02784 | NM_002051.2 | 756 | Complete match |
| GATA4 | FXC23192,FHC23192 | NM_002052.3 | 663 | Complete match |

| | | | | |
|---|---|---|---|---|
| GATAD1 | FXC10715,FHC10715 | NM_021167.4 | 396 | Complete match |
| GATAD2A | FXC05277,FHC05277 | NM_001300946.1 | 642 | Complete match |
| GATAD2B | FXC07759,FHC07759,ORK07759,ORH07759 | NM_020699.2 | 421 | Complete match |
| GCM1 | FHC22660,ORH22660P | NM_003643.3 | 343 | Complete match |
| GFI1 | FXC02154,FHC02154 | NM_005263.3 | 314 | Complete match |
| GLI2 | FXC07749,FHC07749 | NM_005270.4 | 503 | Complete match |
| GLI3 | FXC07751,FHC07751 | NM_000168.5 | 326 | Complete match |
| GLI4 | FHC08726,ORS08726 | NM_138465.3 | 289 | Complete match |
| GLIS1 | FHC20314 | NM_147193.2 | 355 | Complete match |
| GLIS2 | FXC03277,FHC03277 | NM_032575.2 | 737 | Complete match |
| GLIS3 | FHC10842 | NM_001042413.1 | 542 | Complete match |
| GLMP | ORH20629P | NM_144580.2 | 523 | Complete match |
| GMEB1 | FXC02637,FHC02637,ORH02637P | NM_024482.2 | 613 | Complete match |
| GPBP1 | FXC02884,FHC02884,ORH02884 | NM_001127236.2 | 867 | Complete match |
| GPBP1L1 | FXC03797,FHC03797 | NM_021639.4 | 588 | Complete match |
| GPER1 | FXC01743,FHC01743,ORK01743 | NM_001505.2 | 404 | Complete match |
| GREM1 | FHC25323,ORH25323P | NM_013372.6 | 453 | Complete match |
| GRHL1 | ORH28687P | NM_198182.2 | 681 | Complete match |
| GRHL2 | FXC03945,FHC03945,ORH03945P | NM_024915.3 | 497 | Complete match |
| GRHL3 | FHC14282,ORH14282 | NM_198173.2 | 297 | Partial match |
| GSX1 | FXC03254,FHC03254 | NM_145657.2 | 756 | Complete match |
| GTF2A2 | FXC02483,FHC02483,ORS02483,ORH02483 | NM_004492.2 | 348 | Complete match |
| GTF2H2 | FXC03265,FHC03265 | NM_001515.3 | 499 | Complete match |
| GTF2H3 | FXC02538,FHC02538,ORH02538P | NM_001516.4 | 601 | Complete match |
| GTF2H4 | FHC09855,ORS09855,ORH09855 | NM_001517.4 | 539 | Complete match |
| GTF2IRD1 | FXC03811,FHC03811 | NM_001199207.1 | 531 | Complete match |
| GZF1 | FXC03284,FHC03284 | NM_022482.3 | 473 | Complete match |
| HAND1 | FXC05393,FHC05393 | NM_004821.2 | 478 | Complete match |
| HAND2 | FXC03088,FHC03088 | NM_021973.2 | 461 | Complete match |
| HAVCR2 | FHC22366,ORH22366P | NM_032782.4 | 494 | Complete match |
| HCFC1 | FHC30375 | NM_005334.2 | 465 | Complete match |
| HCK | ORH13531P | NM_001172129.1 | 441 | Complete match |
| HDAC1 | FXC02563,FHC02563,ORS02563,ORH02563 | NM_004964.2 | 732 | Complete match |
| HDAC2 | FHC31712 | NM_001527.3 | 425 | Complete match |
| HDAC3 | FXC02559,FHC02559 | NM_003883.3 | 379 | Complete match |
| HDAC4 | FXC01664,FHC01664,ORK01664 | NM_006037.3 | 498 | Complete match |
| HDAC5 | FXC01982,FHC01982,ORK01982 | NM_005474.4 | 461 | Partial match |
| HES6 | FXC02533,FHC02533,ORH02533P | NM_018645.5 | 358 | Complete match |
| HEY1 | FXC07766,FHC07766,ORK07766,ORH07766P | NM_012258.3 | 445 | Complete match |
| HEY2 | FXC05418,FHC05418,ORH05418P | NM_012259.2 | 498 | Complete match |
| HEYL | FXC02547,FHC02547,ORH02547P | NM_014571.3 | 445 | Complete match |
| HHEX | FXC03933,FHC03933 | NM_002729.4 | 744 | Complete match |
| HIF1A | FXC01550,FHC01550,ORK01550 | NM_001530.3 | 543 | Complete match |

| | | | | |
|---|---|---|---|---|
| HIF3A | FXC03642,FHC03642 | NM_152795.3 | 744 | Complete match |
| HINFP | FHC24492 | NM_015517.4 | 396 | Complete match |
| HIPK2 | FHC11119 | NM_001113239.2 | 433 | Complete match |
| HIRA | FXC01218,FHC01218,ORK01218 | NM_003325.3 | 392 | Complete match |
| HIVEP3 | FXC01693,FHC01693,ORK01693 | NM_024503.4 | 534 | Complete match |
| HLF | FXC10520,FHC10520,ORH10520P | NM_002126.4 | 322 | Complete match |
| HMG20A | FXC02213,FHC02213,ORH02213P | NM_018200.3 | 335 | Complete match |
| HMGA1 | FHC08399,ORS08399,ORH08399 | NM_145899.2 | 814 | Complete match |
| HMGA2 | FHC28157 | NM_003483.4 | 855 | Complete match |
| HMGB1 | FXC03405,FHC03405 | NM_002128.4 | 1261 | Complete match |
| HMGB2 | FXC02522,FHC02522,ORS02522,ORH02522 | NM_002129.3 | 1694 | Complete match |
| HMGN3 | ORH22685 | NM_138730.2 | 441 | Complete match |
| HMOX1 | FXC09676,FHC09676,ORS09676,ORH09676 | NM_002133.2 | 600 | Complete match |
| HNF1A | FXC01943,FHC01943 | NM_000545.5 | 817 | Complete match |
| HNF1B | FXC08908,FHC08908,ORS08908,ORH08908 | NM_000458.3 | 515 | Complete match |
| HNF4A | FXC03271,FHC03271 | NM_000457.4 | 646 | Complete match |
| HNF4G | FHC23316 | NM_004133.4 | 539 | Complete match |
| HNRNPAB | FXC05437,FHC05437 | NM_004499.3 | 644 | Complete match |
| HNRNPK | FHC07856,ORS07856,ORH07856 | NM_002140.3 | 2077 | Complete match |
| HOMEZ | FXC00839,FHC00839,ORK00839 | NM_020834.2 | 461 | Complete match |
| HOXA10 | FXC02805,FHC02805,ORS02805,ORH02805 | NM_018951.4 | 710 | Partial match |
| HOXA2 | FXC03127,FHC03127,ORH03127P | NM_006735.3 | 904 | Complete match |
| HOXA3 | FXC02569,FHC02569,ORS02569,ORH02569 | NM_030661.4 | 851 | Complete match |
| HOXA4 | FXC05440,FHC05440 | NM_002141.4 | 498 | Complete match |
| HOXA5 | FXC10008,FHC10008,ORS10008,ORH10008 | NM_019102.3 | 781 | Complete match |
| HOXA6 | FXC06355,FHC06355,ORH06355P | NM_024014.3 | 510 | Complete match |
| HOXA7 | FXC03251,FHC03251 | NM_006896.3 | 588 | Complete match |
| HOXB2 | FHC26088,ORH26088 | NM_002145.3 | 600 | Complete match |
| HOXB3 | FXC03268,FHC03268,ORS03268P,ORH03268P | NM_002146.4 | 915 | Complete match |
| HOXB4 | FHC26089 | NM_024015.4 | 433 | Complete match |
| HOXB5 | FXC03099,FHC03099,ORH03099P | NM_002147.3 | 490 | Complete match |
| HOXB6 | FXC03090,FHC03090,ORS03090,ORH03090P | NM_018952.4 | 851 | Complete match |
| HOXB7 | FHC26090,ORH26090P | NM_004502.3 | 486 | Complete match |
| HOXB8 | FXC03252,FHC03252 | NM_024016.3 | 810 | Complete match |
| HOXC13 | FHC16522 | NM_017410.2 | 384 | Complete match |
| HOXC4 | FXC03098,FHC03098,ORH03098P | NM_014620.5 | 609 | Complete match |
| HOXC5 | FXC03248,FHC03248 | NM_018953.3 | 462 | Complete match |
| HOXC6 | FXC06704,FHC06704 | NM_004503.3 | 588 | Complete match |
| HOXC8 | FXC03093,FHC03093,ORH03093P | NM_022658.3 | 686 | Complete match |
| HOXD10 | FXC03117,FHC03117,ORH03117 | NM_002148.3 | 687 | Complete match |
| HOXD13 | ORS12037,ORH12037 | NM_000523.3 | 420 | Complete match |

| | | | | |
|---|---|---|---|---|
| HOXD3 | FXC02562,FHC02562,ORS02562,ORH02562 | NM_006898.4 | 843 | Complete match |
| HOXD4 | FXC03096,FHC03096,ORH03096P | NM_014621.2 | 498 | Complete match |
| HOXD8 | FXC03106,FHC03106 | NM_019558.3 | 778 | Complete match |
| HOXD9 | FXC05446,FHC05446 | NM_014213.4 | 654 | Partial match |
| HR | FXC01265,FHC01265,ORK01265 | NM_005144.4 | 347 | Complete match |
| HSF2 | FXC05459,FHC05459,ORH05459P | NM_004506.3 | 622 | Complete match |
| HSF4 | FXC02410,FHC02410,ORS02410,ORH02410 | NM_001538.3 | 670 | Complete match |
| HSFY1 | ORH12828P | NM_001001877.1 | 400 | Partial match |
| HSPA1A | FXC07555,FHC07555,ORK07555,ORH07555P | NM_005345.5 | 363 | Complete match |
| ICAM1 | FHC10061,ORS10061,ORH10061 | NM_000201.2 | 396 | Complete match |
| ID1 | FXC03244,FHC03244 | NM_181353.2 | 425 | Complete match |
| ID2 | FXC05482,FHC05482,ORS05482,ORH05482 | NM_002166.4 | 305 | Complete match |
| ID3 | FXC09826,FHC09826,ORS09826,ORH09826 | NM_002167.4 | 321 | Complete match |
| IFI16 | ORH13794P | NM_005531.2 | 493 | Complete match |
| IGF1R | FXC01578,FHC01578,ORK01578 | NM_001291858.1 | 457 | Complete match |
| IKBKB | FXC03657,FHC03657 | NM_001556.2 | 465 | Complete match |
| IKZF1 | FXC05503,FHC05503 | NM_006060.5 | 2757 | Complete match |
| IKZF2 | ORH15504P | NM_001079526.1 | 600 | Complete match |
| IKZF4 | FXC00279,FHC00279,ORK00279 | NM_001351091.1 | 797 | Partial match |
| IRF1 | FXC08430,FHC08430,ORS08430,ORH08430 | NM_002198.2 | 908 | Complete match |
| IRF2 | FXC05536,FHC05536,ORH05536P | NM_002199.3 | 1366 | Complete match |
| IRF4 | FXC02156,FHC02156,ORH02156P | NM_002460.3 | 309 | Complete match |
| IRF5 | ORS08164,ORH08164 | NM_032643.4 | 716 | Complete match |
| IRF6 | FXC02101,FHC02101 | NM_006147.3 | 318 | Complete match |
| IRF7 | FXC07725,FHC07725 | NM_001572.3 | 494 | Complete match |
| IRF8 | FXC02557,FHC02557 | NM_002163.2 | 1067 | Complete match |
| IRF9 | FHC25116,ORH25116 | NM_006084.4 | 734 | Complete match |
| ISL1 | FXC11363,FHC11363 | NM_002202.2 | 468 | Complete match |
| ISX | FXC02489,FHC02489,ORH02489P | NM_001303508.1 | 405 | Complete match |
| ITCH | FHC11019 | NM_031483.5 | 409 | Complete match |
| ITGB2 | FXC03720,FHC03720 | NM_000211.4 | 510 | Complete match |
| JAK2 | FHC23450 | NM_004972.3 | 609 | Complete match |
| JDP2 | FXC02498,FHC02498 | NM_130469.3 | 551 | Complete match |
| JUN | FXC01532,FHC01532,ORH01532 | NM_002228.3 | 447 | Complete match |
| JUNB | FXC03200,FHC03200,ORS03200,ORH03200 | NM_002229.2 | 444 | Complete match |
| JUP | FXC01242,FHC01242,ORK01242,ORS01242,ORH01242 | NM_002230.2 | 347 | Complete match |
| KAT7 | FXC06090,FHC06090 | NM_001199155.1 | 388 | Complete match |
| KCNIP3 | FXC08652,FHC08652,ORS08652,ORH08652 | NM_013434.4 | 410 | Complete match |
| KDM1A | FXC00571,FHC00571,ORK00571 | NM_015013.3 | 691 | Complete match |
| KDM3A | FXC01605,FHC01605,ORK01605,ORS01605 | NM_018433.5 | 511 | Complete match |

| | | | | |
|---|---|---|---|---|
| KDM5A | FXC01704,FHC01704,ORK01704 | NM_001042603.2 | 691 | Complete match |
| KDM5B | FXC27753,FHC27753 | NM_006618.3 | 555 | Complete match |
| KEAP1 | FXC00420,FHC00420,ORK00420 | NM_012289.3 | 384 | Complete match |
| KIT | FXC01839,FHC01839,ORK01839 | NM_001093772.1 | 408 | Complete match |
| KLF1 | FXC05755,FHC05755 | NM_006563.3 | 400 | Complete match |
| KLF10 | FXC01351,FHC01351,ORK01351 | NM_001032282.3 | 380 | Complete match |
| KLF11 | FXC02371,FHC02371,ORK02371 | NM_001177716.1 | 367 | Complete match |
| KLF12 | FXC01441,FHC01441,ORS01441,ORH01441 | NM_007249.4 | 498 | Complete match |
| KLF13 | FHC25320 | NM_015995.3 | 797 | Complete match |
| KLF15 | FXC02553,FHC02553,ORS02553,ORH02553 | NM_014079.3 | 458 | Complete match |
| KLF2 | FHC11769 | NM_016270.2 | 441 | Complete match |
| KLF3 | FXC01292,FHC01292,ORH01292P | NM_016531.5 | 392 | Complete match |
| KLF4 | FXC01442,FHC01442,ORK01442,ORS01442,ORH01442 | NM_004235.4 | 736 | Complete match |
| KLF6 | FXC07848,FHC07848,ORS07848 | NM_001300.5 | 391 | Complete match |
| KLF7 | FXC05756,FHC05756,ORK05756 | NM_003709.3 | 405 | Complete match |
| KLF9 | FXC03094,FHC03094 | NM_001206.2 | 566 | Complete match |
| KRAS | FXC10770,FHC10770 | NM_004985.4 | 486 | Complete match |
| L3MBTL1 | FHC11979,ORH11979 | NM_015478.6 | 383 | Complete match |
| L3MBTL4 | FXC01942,FHC01942 | NM_001365765.1 | 568 | Partial match |
| LBX1 | FHC27877 | NM_006562.4 | 753 | Complete match |
| LCOR | FXC00280,FHC00280,ORK00280 | NM_032440.3 | 445 | Complete match |
| LEF1 | FXC16601,FHC16601 | NM_001130713.2 | 691 | Complete match |
| LGALS9 | ORH25936P | NM_002308.3 | 404 | Complete match |
| LHX1 | FXC02613,FHC02613,ORK02613,ORH02613 | NM_005568.4 | 605 | Complete match |
| LHX6 | FHC05826,ORH05826 | NM_001242334.1 | 363 | Complete match |
| LMO2 | FXC02167,FHC02167,ORK02167,ORS02167,ORH02167 | NM_001142315.1 | 265 | Complete match |
| LMO4 | FXC01247,FHC01247,ORK01247,ORS01247 | NM_006769.3 | 334 | Complete match |
| LMX1B | FXC10792,FHC10792,ORH10792P | NM_002316.3 | 609 | Complete match |
| LRP5 | FHC24344 | NM_002335.3 | 778 | Complete match |
| LRP6 | FHC36555M | NM_002336.2 | 920 | Complete match |
| LTF | FXC03739,FHC03739 | NM_002343.4 | 689 | Complete match |
| LYL1 | FXC02526,FHC02526 | NM_005583.4 | 440 | Complete match |
| LZTR1 | FXC01911,FHC01911,ORK01911 | NM_006767.3 | 552 | Complete match |
| LZTS1 | FHC23208,ORH23208 | NM_021020.3 | 432 | Complete match |
| MAFB | FXC03224,FHC03224,ORH03224P | NM_005461.4 | 740 | Complete match |
| MAFF | FHC27251,ORH27251P | NM_012323.3 | 482 | Complete match |
| MAFG | FXC03078,FHC03078,ORS03078 | NM_002359.3 | 662 | Complete match |
| MAFK | FXC01764,FHC01764,ORH01764 | NM_002360.3 | 384 | Complete match |
| MAX | FXC02095,FHC02095,ORH02095P | NM_002382.4 | 416 | Complete match |
| MBTPS2 | FXC01877,FHC01877 | NM_015884.3 | 359 | Complete match |
| MECOM | FXC03843,FHC03843,ORH03843P | NM_001366473.1 | 556 | Partial match |
| MECP2 | FXC02747,FHC02747,ORS02747,ORH02747 | NM_004992.3 | 437 | Complete match |

| | | | | |
|---|---|---|---|---|
| MED1 | FHC16545 | NM_004774.4 | 417 | Partial match |
| MEF2A | FXC01295,FHC01295,ORK01295 | NM_001130926.1 | 350 | Complete match |
| MEF2B | FHC10992 | NM_001145785.1 | 527 | Complete match |
| MEF2C | FXC01355,FHC01355 | NM_002397.4 | 355 | Complete match |
| MEF2D | FXC00981,FHC00981,ORK00981 | NM_005920.4 | 263 | Partial match |
| MEIS1 | FXC03842,FHC03842 | NM_002398.2 | 390 | Complete match |
| MEIS2 | FXC01422,FHC01422,ORK01422 | NM_170677.4 | 387 | Complete match |
| MEIS3 | FXC02636,FHC02636,ORK02636 | NM_001009813.2 | 383 | Complete match |
| MEN1 | FXC10467,FHC10467 | NM_000244.3 | 473 | Complete match |
| MEOX1 | FXC03676,FHC03676,ORH03676 | NM_004527.3 | 888 | Complete match |
| MEOX2 | FXC05969,FHC05969 | NM_005924.4 | 810 | Complete match |
| MESP1 | FHC25503 | NM_018670.3 | 531 | Complete match |
| MID2 | FHC12697 | NM_052817.2 | 588 | Complete match |
| MITF | FXC03436,FHC03436 | NM_198159.2 | 662 | Complete match |
| MIXL1 | FHC20841,ORH20841P | NM_031944.2 | 1067 | Complete match |
| MKL1 | FXC00835,FHC00835,ORK00835 | NM_020831.4 | 522 | Complete match |
| MKL2 | FHC13131,ORH13131 | NM_001365416.1 | 453 | Partial match |
| MKX | FXC03670,FHC03670,ORH03670P | NM_173576.2 | 384 | Complete match |
| MLLT10 | FHC12050,ORH12050 | NM_004641.3 | 376 | Complete match |
| MLX | FXC03683,FHC03683 | NM_198205.1 | 419 | Complete match |
| MLXIP | ORH15215P | NM_014938.6 | 354 | Partial match |
| MLXIPL | FXC03526,FHC03526 | NM_032951.2 | 585 | Complete match |
| MSC | FXC02517,FHC02517,ORH02517P | NM_005098.3 | 572 | Complete match |
| MSGN1 | ORS29953 | NM_001105569.1 | 432 | Complete match |
| MSRB2 | FHC11532,ORH11532P | NM_012228.3 | 851 | Complete match |
| MSX1 | FXC06045,FHC06045 | NM_002448.3 | 493 | Partial match |
| MSX2 | FHC08787,ORS08787,ORH08787 | NM_002449.4 | 542 | Complete match |
| MTA1 | FHC10817 | NM_004689.4 | 428 | Partial match |
| MTA2 | FXC03608,FHC03608,ORH03608P | NM_004739.3 | 535 | Complete match |
| MTA3 | FXC00789,FHC00789,ORK00789 | NM_001282755.1 | 421 | Complete match |
| MTDH | ORH23352P | NM_178812.3 | 429 | Complete match |
| MTF1 | FXC03655,FHC03655 | NM_005955.2 | 334 | Complete match |
| MTPN | FXC06060,FHC06060,ORH06060 | NM_145808.3 | 314 | Complete match |
| MXD1 | FXC02529,FHC02529,ORH02529P | NM_002357.3 | 469 | Complete match |
| MYB | FXC03658,FHC03658,ORH03658 | NM_005375.2 | 556 | Complete match |
| MYBL1 | FHC31237 | NM_001080416.3 | 490 | Complete match |
| MYBL2 | FXC09919,FHC09919,ORS09919,ORH09919 | NM_002466.3 | 437 | Complete match |
| MYC | FXC10454,FHC10454 | NM_002467.4 | 309 | Complete match |
| MYCL | FXC03677,FHC03677 | NM_001033082.2 | 437 | Complete match |
| MYD88 | FXC06070,FHC06070 | NM_002468.4 | 506 | Complete match |
| MYF5 | FXC11352,FHC11352 | NM_005593.2 | 384 | Complete match |
| MYF6 | FXC08920,FHC08920,ORS08920,ORH08920 | NM_002469.2 | 416 | Complete match |
| MYNN | FXC09480,FHC09480,ORS09480,ORH09480 | NM_018657.4 | 510 | Complete match |
| MYOCD | ORH12815P | NM_001146312.2 | 392 | Complete match |

| | | | | |
|---|---|---|---|---|
| MYOD1 | FXC02545,FHC02545,ORH02545 | NM_002478.4 | 477 | Complete match |
| MYOG | FXC02531,FHC02531,ORH02531P | NM_002479.5 | 428 | Complete match |
| MYPOP | FHC12648,ORH12648 | NM_001012643.3 | 916 | Complete match |
| MYRF | FXC01138,FHC01138 | NM_001127392.2 | 646 | Complete match |
| MYT1L | FXC01147,FHC01147,ORK01147 | NM_001329848.1 | 669 | Partial match |
| MZF1 | FHC12723,ORH12723P | NM_003422.2 | 355 | Complete match |
| NACC2 | FHC08802,ORS08802,ORH08802 | NM_144653.4 | 464 | Complete match |
| NANOG | FHC12077 | NM_024865.3 | 388 | Complete match |
| NCOA3 | FXC01529,FHC01529 | NM_001174087.1 | 837 | Complete match |
| NCOR1 | FXC00170,FHC00170,ORK00170 | NM_001190440.1 | 456 | Complete match |
| NDN | FXC03337,FHC03337,ORH03337P | NM_002487.2 | 744 | Complete match |
| NDP | FXC10830,FHC10830,ORH10830 | NM_000266.3 | 482 | Complete match |
| NEUROD1 | FXC01498,FHC01498,ORK01498,ORS01498,ORH01498 | NM_002500.4 | 517 | Complete match |
| NEUROD2 | FXC03205,FHC03205,ORH03205P | NM_006160.3 | 609 | Complete match |
| NEUROD6 | FXC03207,FHC03207 | NM_022728.3 | 600 | Complete match |
| NEUROG1 | FXC03208,FHC03208,ORS03208,ORH03208 | NM_006161.2 | 481 | Complete match |
| NEUROG2 | FXC03209,FHC03209 | NM_024019.3 | 490 | Complete match |
| NFAM1 | FXC03389,FHC03389,ORH03389 | NM_145912.5 | 470 | Complete match |
| NFATC1 | FHC12349,ORH12349 | NM_172390.2 | 355 | Complete match |
| NFATC3 | FHC09749,ORH09749 | NM_173165.2 | 685 | Complete match |
| NFE2 | FXC06142,FHC06142,ORS06142,ORH06142 | NM_006163.2 | 355 | Complete match |
| NFE2L1 | FXC01045,FHC01045,ORK01045 | NM_003204.2 | 691 | Complete match |
| NFE2L2 | FXC06144,FHC06144,ORH06144P | NM_006164.4 | 236 | Complete match |
| NFIA | FXC00836,FHC00836,ORK00836 | NM_001134673.3 | 494 | Complete match |
| NFIC | FHC08611,ORS08611,ORH08611 | NM_005597.3 | 466 | Complete match |
| NFIL3 | FXC03211,FHC03211,ORS03211,ORH03211P | NM_005384.2 | 498 | Complete match |
| NFIX | FXC01876,FHC01876 | NM_002501.3 | 428 | Partial match |
| NFKB1 | FXC07774,FHC07774,ORK07774 | NM_001165412.1 | 453 | Complete match |
| NFKB2 | FXC01287,FHC01287 | NM_002502.5 | 514 | Complete match |
| NFKBIA | FXC10563,FHC10563,ORH10563P | NM_020529.2 | 502 | Complete match |
| NFKBIB | FXC06146,FHC06146,ORS06146,ORH06146 | NM_002503.4 | 333 | Complete match |
| NFKBID | ORH29921P | NM_139239.1 | 417 | Complete match |
| NFKBIL1 | FXC02810,FHC02810,ORH02810 | NM_005007.3 | 641 | Complete match |
| NFX1 | FXC01879,FHC01879,ORK01879 | NM_147134.2 | 502 | Complete match |
| NFYB | FXC02518,FHC02518,ORS02518,ORH02518 | NM_006166.3 | 470 | Complete match |
| NFYC | FXC01734,FHC01734 | NM_001142587.1 | 358 | Complete match |
| NHLH1 | FXC03204,FHC03204,ORH03204P | NM_005598.3 | 412 | Complete match |
| NHLH2 | FXC07617,FHC07617,ORH07617 | NM_005599.3 | 502 | Complete match |
| NKX2-1 | FHC10939 | NM_003317.3 | 727 | Complete match |
| NKX2-2 | FXC03100,FHC03100,ORH03100 | NM_002509.3 | 798 | Complete match |
| NKX2-5 | FXC06155,FHC06155,ORS06155,ORH06155 | NM_004387.3 | 625 | Complete match |
| NKX2-6 | FXC03107,FHC03107 | NM_001136271.2 | 834 | Complete match |

| | | | | |
|---|---|---|---|---|
| NKX2-8 | FHC25144 | NM_014360.2 | 691 | Complete match |
| NKX3-1 | FXC06156,FHC06156,ORH06156 | NM_006167.3 | 424 | Complete match |
| NKX3-2 | FXC06157,FHC06157 | NM_001189.3 | 498 | Complete match |
| NKX6-1 | FHC12664 | NM_006168.2 | 790 | Complete match |
| NKX6-2 | FHC24074 | NM_177400.2 | 757 | Complete match |
| NLRC4 | FHC28696 | NM_021209.4 | 379 | Complete match |
| NLRP12 | FHC09241,ORS09241,ORH09241 | NM_144687.3 | 416 | Complete match |
| NLRP3 | FXC01254,FHC01254,ORK01254 | NM_001127461.2 | 678 | Complete match |
| NOBOX | FXC10888,FHC10888,ORH10888 | NM_001080413.3 | 506 | Partial match |
| NOD1 | ORH22889P | NM_006092.2 | 526 | Complete match |
| NOD2 | FXC11015,FHC11015,ORH11015 | NM_022162.2 | 613 | Complete match |
| NODAL | FHC12598,ORH12598P | NM_018055.4 | 584 | Complete match |
| NPAS1 | FHC26706,ORH26706P | NM_002517.2 | 452 | Complete match |
| NPAS2 | FHC12549 | NM_002518.3 | 517 | Complete match |
| NPAS3 | FXC06177,FHC06177 | NM_001165893.1 | 911 | Complete match |
| NPM1 | FXC02685,FHC02685,ORS02685,ORH02685 | NM_002520.6 | 1190 | Complete match |
| NR0B1 | FXC02225,FHC02225,ORH02225P | NM_000475.4 | 490 | Complete match |
| NR0B2 | FHC20160,ORH20160P | NM_021969.2 | 556 | Complete match |
| NR1D1 | FXC06190,FHC06190 | NM_021724.4 | 306 | Complete match |
| NR1D2 | FXC01850,FHC01850,ORK01850 | NM_005126.4 | 425 | Complete match |
| NR1H3 | FXC06191,FHC06191 | NM_005693.3 | 457 | Complete match |
| NR1H4 | FXC02564,FHC02564 | NM_001206992.1 | 647 | Complete match |
| NR1I2 | FXC31716,FHC31716 | NM_003889.3 | 559 | Complete match |
| NR1I3 | FHC20672 | NM_001077482.2 | 396 | Complete match |
| NR2C1 | FHC24833 | NM_003297.3 | 524 | Complete match |
| NR2C2 | FXC01537,FHC01537 | NM_003298.4 | 707 | Complete match |
| NR2E1 | FXC10500,FHC10500,ORH10500P | NM_003269.4 | 707 | Complete match |
| NR2F2 | FHC10946 | NM_021005.3 | 490 | Complete match |
| NR3C1 | FXC10483,FHC10483,ORK10483,ORH10483P | NM_000176.2 | 515 | Complete match |
| NR3C2 | FXC22071,FHC22071 | NM_000901.4 | 347 | Complete match |
| NR4A1 | FHC08830,ORS08830,ORH08830 | NM_002135.4 | 491 | Complete match |
| NR4A2 | FHC21198 | NM_006186.3 | 654 | Complete match |
| NR4A3 | FXC01270,FHC01270 | NM_006981.3 | 633 | Complete match |
| NR5A1 | FXC02552,FHC02552,ORH02552 | NM_004959.4 | 793 | Complete match |
| NR5A2 | FHC11953 | NM_003822.4 | 567 | Complete match |
| NR6A1 | FHC12632 | NM_001489.4 | 638 | Complete match |
| NRF1 | FXC06193,FHC06193 | NM_005011.4 | 412 | Complete match |
| NRL | FXC06194,FHC06194 | NM_006177.3 | 801 | Complete match |
| NTRK1 | FHC20644 | NM_002529.3 | 523 | Complete match |
| NTRK1 | FHC20644 | NM_002529.3 | 482 | Complete match |
| NUCKS1 | FXC09741,FHC09741,ORS09741 | NM_022731.4 | 355 | Complete match |
| ONECUT1 | FXC03270,FHC03270,ORH03270P | NM_004498.2 | 539 | Complete match |
| OPRD1 | FXC02316,FHC02316,ORH02316 | NM_000911.3 | 638 | Complete match |
| OSR2 | ORH31689P | NM_053001.3 | 543 | Complete match |

| | | | | |
|---|---|---|---|---|
| OTX1 | FXC03934,FHC03934 | NM_014562.3 | 502 | Complete match |
| OTX2 | FXC09429,FHC09429,ORS09429,ORH09429 | NM_021728.3 | 863 | Complete match |
| OVOL1 | FXC02512,FHC02512,ORH02512P | NM_004561.3 | 620 | Complete match |
| OVOL2 | FXC02521,FHC02521,ORH02521P | NM_021220.3 | 859 | Complete match |
| PA2G4 | FXC02725,FHC02725,ORS02725,ORH02725 | NM_006191.2 | 436 | Complete match |
| PARK2 | FXC03885,FHC03885 | NM_004562.2 | 351 | Complete match |
| PARK7 | FXC06271,FHC06271 | NM_007262.4 | 312 | Complete match |
| PATZ1 | FXC01717,FHC01717 | NM_032050.1 | 926 | Complete match |
| PAX3 | FXC21337 | NM_181457.3 | 844 | Complete match |
| PAX4 | FXC11401,FHC11401 | NM_006193.2 | 609 | Complete match |
| PAX5 | FXC11730,FHC11730,ORH11730 | NM_016734.2 | 695 | Complete match |
| PAX6 | FXC08594,FHC08594,ORS08594,ORH08594 | NM_000280.4 | 1160 | Complete match |
| PAX7 | FHC20120,ORH20120P | NM_013945.2 | 850 | Complete match |
| PAX8 | FXC09750,FHC09750,ORS09750,ORH09750 | NM_003466.3 | 494 | Complete match |
| PAX9 | FXC06282,FHC06282,ORS06282,ORH06282 | NM_006194.3 | 841 | Complete match |
| PAXBP1 | FHC27155,ORH27155 | NM_016631.3 | 318 | Complete match |
| PBX1 | FXC01756,FHC01756 | NM_002585.3 | 448 | Complete match |
| PBX2 | FXC06283,FHC06283,ORH06283P | NM_002586.4 | 436 | Complete match |
| PBX3 | FXC06284,FHC06284 | NM_006195.5 | 407 | Complete match |
| PBX4 | FHC26553,ORH26553P | NM_025245.2 | 494 | Complete match |
| PCGF2 | FXC02780,FHC02780,ORH02780 | NM_007144.2 | 326 | Complete match |
| PCGF6 | FXC06019,FHC06019 | NM_001011663.1 | 556 | Complete match |
| PDX1 | FXC03103,FHC03103 | NM_000209.3 | 1421 | Complete match |
| PEG3 | FXC00496,FHC00496,ORK00496 | NM_001146187.1 | 625 | Complete match |
| PELI1 | FHC21022,ORH21022P | NM_020651.3 | 375 | Complete match |
| PEX14 | FXC03505,FHC03505 | NM_004565.2 | 514 | Complete match |
| PGBD1 | FHC12721,ORH12721P | NM_032507.3 | 453 | Complete match |
| PHB | FXC02675,FHC02675 | NM_002634.3 | 355 | Complete match |
| PHB2 | FXC08753,FHC08753,ORS08753,ORH08753 | NM_001144831.1 | 621 | Complete match |
| PHF1 | FHC11093 | NM_024165.2 | 469 | Complete match |
| PHF5A | FXC06346,FHC06346,ORH06346 | NM_032758.3 | 301 | Complete match |
| PHOX2A | FHC24373,ORH24373 | NM_005169.3 | 709 | Complete match |
| PHTF1 | FHC20481 | NM_006608.2 | 670 | Complete match |
| PIAS2 | FXC03457,FHC03457 | NM_004671.3 | 1101 | Complete match |
| PIAS4 | FXC06356,FHC06356 | NM_015897.3 | 740 | Complete match |
| PITX1 | FXC02541,FHC02541,ORH02541P | NM_002653.4 | 437 | Complete match |
| PITX2 | FXC08718,FHC08718,ORS08718 | NM_000325.5 | 485 | Complete match |
| PITX3 | FXC02537,FHC02537,ORS02537,ORH02537 | NM_005029.3 | 1007 | Complete match |
| PKNOX1 | FXC01386,FHC01386,ORH01386P | NM_004571.4 | 421 | Complete match |
| PLA2G1B | FXC06377,FHC06377,ORH06377P | NM_000928.2 | 551 | Complete match |
| PLAG1 | FXC01334,FHC01334 | NM_002655.2 | 387 | Complete match |
| PLAGL1 | FXC03214,FHC03214 | NM_002656.3 | 564 | Complete match |

| | | | | |
|---|---|---|---|---|
| PLAGL2 | FXC00452,FHC00452,ORK00452,ORS00452,ORH00452 | NM_002657.3 | 458 | Complete match |
| PLSCR1 | FHC09448,ORS09448,ORH09448 | NM_021105.2 | 400 | Complete match |
| POU1F1 | FXC11948,FHC11948,ORH11948 | NM_000306.3 | 760 | Complete match |
| POU2F1 | FXC03170,FHC03170,ORK03170 | NM_001365849.1 | 564 | Partial match |
| POU2F2 | FHC28631 | NM_002698.4 | 773 | Complete match |
| POU2F3 | FHC24501,ORH24501 | NM_014352.3 | 597 | Complete match |
| POU3F1 | FHC27644,ORH27644 | NM_002699.3 | 818 | Complete match |
| POU3F4 | FXC03215,FHC03215 | NM_000307.4 | 1060 | Complete match |
| POU4F1 | FHC16557 | NM_006237.3 | 481 | Complete match |
| POU4F2 | FHC16524 | NM_004575.2 | 506 | Complete match |
| POU4F3 | FXC03116,FHC03116,ORH03116P | NM_002700.2 | 359 | Complete match |
| POU5F1 | FXC07739,FHC07739,ORH07739 | NM_002701.5 | 534 | Complete match |
| POU5F1B | FHC16556 | NM_001159542.1 | 365 | Complete match |
| POU5F2 | FXC03892,FHC03892 | NM_153216.1 | 519 | Complete match |
| POU6F2 | FXC11729,FHC11729,ORH11729 | NM_007252.3 | 764 | Partial match |
| PPARA | FXC03156,FHC03156,ORH03156 | NM_005036.4 | 498 | Complete match |
| PPARD | FXC15718,ORH15718 | NM_177435.2 | 424 | Complete match |
| PPARG | FXC08305,FHC08305,ORS08305,ORH08305 | NM_005037.5 | 333 | Complete match |
| PPARGC1A | FXC21879M,FHC21879,ORH21879 | NM_013261.3 | 433 | Complete match |
| PRDM1 | ORH15364P | NM_182907.2 | 555 | Complete match |
| PRDM5 | FXC06475,FHC06475,ORH06475P | NM_001300823.1 | 405 | Complete match |
| PRDX3 | FXC03411,FHC03411,ORH03411P | NM_006793.4 | 559 | Complete match |
| PRKCB | FXC10533,FHC10533,ORH10533P | NM_002738.6 | 518 | Complete match |
| PRKCH | FXC01867,FHC01867,ORS01867,ORH01867 | NM_006255.4 | 469 | Complete match |
| PRKCI | FHC21760 | NM_002740.5 | 408 | Complete match |
| PRKCQ | FXC03603,FHC03603 | NM_006257.4 | 527 | Complete match |
| PRKCZ | FXC06489,FHC06489,ORH06489P | NM_002744.4 | 518 | Complete match |
| PRKD1 | FXC25129,FHC25129 | NM_002742.2 | 596 | Complete match |
| PRKD2 | FXC26703,FHC26703 | NM_016457.4 | 591 | Complete match |
| PRMT2 | FXC02606,FHC02606,ORH02606P | NM_001535.4 | 425 | Complete match |
| PRNP | FXC02097,FHC02097,ORH02097P | NM_000311.3 | 470 | Complete match |
| PROX1 | FXC01732,FHC01732 | NM_002763.4 | 449 | Complete match |
| PSMA6 | FXC09822,FHC09822,ORS09822 | NM_002791.2 | 468 | Complete match |
| PSMD10 | FXC10631,FHC10631,ORH10631P | NM_002814.3 | 374 | Complete match |
| PTCH1 | FXC10747,FHC10747,ORH10747 | NM_000264.3 | 384 | Complete match |
| PTF1A | FXC07724,FHC07724 | NM_178161.2 | 527 | Complete match |
| PTGER3 | FHC20366 | NM_198718.1 | 416 | Complete match |
| PTGIS | FHC12543,ORH12543 | NM_000961.3 | 564 | Complete match |
| PTH | FXC03075,FHC03075,ORH03075P | NM_000315.2 | 612 | Complete match |
| PTHLH | FHC10954 | NM_198965.1 | 429 | Complete match |
| PTTG1 | FXC02516,FHC02516,ORH02516P | NM_004219.3 | 355 | Complete match |
| PURA | FXC03216,FHC03216 | NM_005859.4 | 363 | Complete match |
| PURB | FXC03333,FHC03333,ORH03333P | NM_033224.4 | 527 | Complete match |
| PYCARD | FXC06547,FHC06547 | NM_013258.4 | 361 | Complete match |

| | | | | |
|---|---|---|---|---|
| PYDC1 | FXC25684,FHC25684 | NM_152901.3 | 350 | Complete match |
| RAB7B | FHC11097,ORH11097P | NM_177403.5 | 371 | Complete match |
| RAD21 | FXC01957,FHC01957,ORK01957,ORH01957P | NM_006265.2 | 535 | Complete match |
| RAI1 | FXC01638,FHC01638,ORK01638 | NM_030665.4 | 298 | Partial match |
| RARA | FHC09936,ORS09936 | NM_000964.3 | 699 | Complete match |
| RARB | FHC21460,ORH21460 | NM_000965.4 | 970 | Complete match |
| RARG | FXC03419,FHC03419,ORH03419 | NM_000966.5 | 841 | Complete match |
| RB1 | FXC01227,FHC01227,ORS01227,ORH01227 | NM_000321.2 | 596 | Complete match |
| RBCK1 | FHC26861 | NM_031229.2 | 379 | Complete match |
| RBPJ | FXC03722,FHC03722 | NM_005349.3 | 736 | Complete match |
| RBPJL | FHC27032,ORH27032 | NM_014276.3 | 539 | Complete match |
| RCAN1 | ORH27162P | NM_004414.7 | 441 | Partial match |
| RCOR1 | FXC02300,FHC02300 | NM_015156.4 | 625 | Partial match |
| REL | FXC06623,FHC06623,ORH06623P | NM_001291746.1 | 474 | Complete match |
| RELA | FXC06624,FHC06624 | NM_021975.3 | 498 | Complete match |
| RELB | FHC30830,ORH30830P | NM_006509.3 | 396 | Complete match |
| REST | FXC01798,FHC01798 | NM_005612.4 | 494 | Complete match |
| REXO4 | FHC07628,ORH07628P | NM_020385.3 | 424 | Complete match |
| RFX1 | FXC11645,FHC11645,ORK11645 | NM_002918.4 | 461 | Complete match |
| RFX2 | FXC03607,FHC03607 | NM_134433.2 | 817 | Complete match |
| RFX3 | FXC03650,FHC03650,ORH03650P | NM_134428.2 | 527 | Complete match |
| RFX4 | ORH10255 | NM_032491.5 | 408 | Complete match |
| RFX5 | FXC02801,FHC02801,ORK02801,ORS02801,ORH02801 | NM_000449.3 | 523 | Complete match |
| RFX6 | ORH22741P | NM_173560.3 | 482 | Complete match |
| RFXANK | FXC06641,FHC06641,ORK06641 | NM_003721.3 | 330 | Complete match |
| RFXAP | FHC07763 | NM_000538.3 | 609 | Complete match |
| RGCC | FHC29713,ORH29713P | NM_014059.2 | 322 | Complete match |
| RHEBL1 | FXC03848,FHC03848 | NM_144593.2 | 497 | Complete match |
| RHOXF1 | FXC02509,FHC02509,ORK02509 | NM_139282.2 | 851 | Complete match |
| RLIM | FXC10005,FHC10005,ORS10005,ORH10005 | NM_016120.3 | 498 | Complete match |
| RNF2 | FXC02551,FHC02551,ORS02551,ORH02551 | NM_007212.3 | 351 | Complete match |
| RNF25 | FXC03430,FHC03430,ORH03430P | NM_022453.2 | 367 | Complete match |
| RNF31 | FXC01571,FHC01571,ORK01571 | NM_017999.4 | 515 | Complete match |
| RNF4 | FXC02720,FHC02720,ORK02720 | NM_002938.4 | 437 | Complete match |
| RNF41 | FXC03829,FHC03829,ORH03829P | NM_005785.3 | 470 | Complete match |
| RORA | FHC25415 | NM_134262.2 | 432 | Complete match |
| RORB | FHC23532 | NM_006914.3 | 929 | Complete match |
| RORC | FHC09376,ORS09376,ORH09376 | NM_005060.3 | 1326 | Complete match |
| RPS27A | FXC11355,FHC11355,ORH11355P | NM_002954.5 | 464 | Complete match |
| RPS3 | FXC02085,FHC02085,ORS02085,ORH02085 | NM_001005.4 | 408 | Complete match |
| RPS6KA4 | FHC11752 | NM_001300802.1 | 503 | Complete match |
| RUNX1 | FXC01784,FHC01784,ORH01784P | NM_001754.4 | 515 | Complete match |

| | | | | |
|---|---|---|---|---|
| RUNX2 | FHC22638 | NM_001024630.4 | 338 | Partial match |
| RUNX3 | FHC20145,ORH20145 | NM_001031680.2 | 354 | Complete match |
| RWDD3 | FXC06713,FHC06713,ORH06713 | NM_001128142.1 | 355 | Complete match |
| RXRA | FXC03289,FHC03289,ORK03289,ORH03289 | NM_002957.5 | 437 | Complete match |
| RXRB | FXC01357,FHC01357 | NM_001270401.1 | 294 | Complete match |
| RXRG | FXC03714,FHC03714,ORH03714 | NM_006917.4 | 302 | Complete match |
| S100A12 | FXC03069,FHC03069,ORH03069P | NM_005621.1 | 285 | Complete match |
| S100A8 | FHC08254,ORS08254,ORH08254 | NM_002964.4 | 317 | Complete match |
| S100A9 | FHC20576,ORH20576 | NM_002965.3 | 354 | Complete match |
| SALL1 | FHC25699 | NM_001127892.1 | 412 | Complete match |
| SALL2 | FXC00516,FHC00516,ORK00516 | NM_005407.2 | 580 | Complete match |
| SALL4 | ORH27068P | NM_020436.3 | 3549 | Complete match |
| SATB1 | FXC11928,FHC11928 | NM_002971.4 | 358 | Complete match |
| SCAND1 | FXC03217,FHC03217,ORH03217 | NM_016558.3 | 424 | Complete match |
| SCMH1 | FXC09056,FHC09056,ORS09056,ORH09056 | NM_001031694.2 | 330 | Complete match |
| SCML1 | FHC27317,ORH27317 | NM_001037535.2 | 379 | Complete match |
| SCRT1 | FXC01341,FHC01341 | NM_031309.5 | 576 | Complete match |
| SCX | FHC30292 | NM_001080514.2 | 405 | Complete match |
| SETD6 | FHC25718 | NM_024860.2 | 314 | Complete match |
| SFRP4 | ORH22908P | NM_003014.3 | 375 | Complete match |
| SHH | FXC02308,FHC02308,ORH02308 | NM_000193.2 | 510 | Complete match |
| SHOX | FHC27276,ORS27276,ORH27276 | NM_006883.2 | 499 | Complete match |
| SIGIRR | FHC08157,ORS08157,ORH08157 | NM_021805.2 | 541 | Complete match |
| SIK1 | FHC10354,ORS10354,ORH10354 | NM_173354.3 | 515 | Complete match |
| SIM1 | FHC06829,ORH06829 | NM_005068.2 | 613 | Complete match |
| SIM2 | FHC11667,ORH11667P | NM_009586.3 | 494 | Complete match |
| SIN3A | FXC11647,FHC11647,ORK11647 | NM_015477.2 | 482 | Complete match |
| SIVA1 | ORH25310P | NM_021709.2 | 331 | Complete match |
| SIX1 | FHC25194,ORH25194P | NM_005982.3 | 456 | Complete match |
| SIX2 | FXC06833,FHC06833,ORH06833P | NM_016932.4 | 289 | Complete match |
| SIX3 | FXC03259,FHC03259 | NM_005413.3 | 645 | Complete match |
| SIX4 | FHC12680 | NM_017420.4 | 342 | Complete match |
| SIX6 | FHC25193,ORH25193 | NM_007374.2 | 617 | Complete match |
| SKIL | FHC28946,ORH28946 | NM_005414.4 | 412 | Complete match |
| SLC26A3 | FXC11726,FHC11726,ORH11726 | NM_000111.2 | 515 | Complete match |
| SLC30A9 | FXC02803,FHC02803,ORH02803P | NM_006345.3 | 601 | Complete match |
| SMAD1 | ORS08032,ORH08032 | NM_005900.2 | 490 | Complete match |
| SMAD2 | FXC03382,FHC03382,ORK03382,ORS03382,ORH03382 | NM_005901.5 | 359 | Complete match |
| SMAD3 | FXC02648,FHC02648,ORH02648P | NM_005902.3 | 527 | Complete match |
| SMAD4 | FXC01560,FHC01560,ORK01560,ORS01560,ORH01560 | NM_005359.5 | 698 | Complete match |
| SMAD5 | FXC06899,FHC06899,ORH06899P | NM_005903.6 | 413 | Complete match |
| SMAD6 | FHC12705 | NM_005585.4 | 395 | Complete match |
| SMAD7 | FXC01575,FHC01575,ORK01575 | NM_005904.3 | 482 | Complete match |

| | | | | |
|---|---|---|---|---|
| SMAD9 | FXC02137,FHC02137 | NM_001127217.2 | 417 | Complete match |
| SMARCB1 | FXC03158,FHC03158,ORH03158 | NM_003073.3 | 433 | Complete match |
| SMO | FXC09959,FHC09959,ORH09959 | NM_005631.4 | 1000 | Complete match |
| SNAI1 | FXC02510,FHC02510,ORS02510,ORH02510 | NM_005985.3 | 691 | Complete match |
| SNAI2 | FXC02514,FHC02514,ORH02514 | NM_003068.4 | 1098 | Complete match |
| SNAI3 | FXC06912,FHC06912,ORH06912P | NM_178310.3 | 551 | Complete match |
| SNAPC2 | FXC03396,FHC03396,ORH03396 | NM_003083.3 | 529 | Complete match |
| SNAPC4 | FHC11081 | NM_003086.2 | 461 | Complete match |
| SNAPC5 | FXC02482,FHC02482 | NM_006049.2 | 250 | Complete match |
| SOHLH1 | ORH29408P | NM_001101677.1 | 531 | Partial match |
| SOHLH2 | FXC03491,FHC03491 | NM_017826.2 | 1040 | Complete match |
| SOX10 | FXC06946,FHC06946,ORH06946 | NM_006941.3 | 716 | Complete match |
| SOX11 | FXC03219,FHC03219 | NM_003108.3 | 794 | Complete match |
| SOX12 | FHC26859,ORH26859P | NM_006943.3 | 534 | Complete match |
| SOX13 | FXC10779,FHC10779 | NM_005686.2 | 559 | Complete match |
| SOX14 | FXC03220,FHC03220,ORH03220P | NM_004189.3 | 558 | Complete match |
| SOX15 | FHC07936,ORS07936,ORH07936 | NM_006942.1 | 1146 | Complete match |
| SOX17 | FXC03267,FHC03267 | NM_022454.3 | 683 | Complete match |
| SOX2 | FXC03221,FHC03221,ORH03221 | NM_003106.3 | 482 | Complete match |
| SOX21 | FXC03222,FHC03222 | NM_007084.3 | 605 | Complete match |
| SOX30 | FXC03612,FHC03612 | NM_178424.1 | 531 | Complete match |
| SOX4 | FHC11999 | NM_003107.2 | 346 | Complete match |
| SOX5 | FHC11740 | NM_006940.4 | 445 | Complete match |
| SOX6 | FXC02819,FHC02819,ORH02819 | NM_017508.2 | 392 | Complete match |
| SOX7 | FXC03130,FHC03130 | NM_031439.3 | 826 | Complete match |
| SOX8 | FXC02570,FHC02570,ORH02570P | NM_014587.4 | 445 | Complete match |
| SOX9 | FXC01349,FHC01349 | NM_000346.3 | 739 | Complete match |
| SP1 | FXC07743,FHC07743 | NM_138473.2 | 343 | Complete match |
| SP100 | FHC13365,ORH13365P | NM_001206702.1 | 263 | Complete match |
| SP3 | FXC07756,FHC07756,ORH07756P | NM_003111.4 | 609 | Complete match |
| SP4 | FHC31424,ORH31424 | NM_003112.3 | 490 | Complete match |
| SPDEF | FXC01941,FHC01941,ORH01941 | NM_012391.2 | 449 | Complete match |
| SPEN | FHC40193M | NM_015001.2 | 953 | Complete match |
| SPHK1 | FHC26201 | NM_182965.2 | 343 | Complete match |
| SPI1 | FXC03256,FHC03256 | NM_003120.2 | 456 | Complete match |
| SPIC | FHC24853,ORH24853P | NM_152323.1 | 433 | Complete match |
| SPOP | FXC02905,FHC02905,ORK02905,ORH02905 | NM_003563.3 | 506 | Complete match |
| SPZ1 | ORS09498,ORH09498 | NM_032567.3 | 498 | Complete match |
| SREBF1 | FHC12086 | NM_001005291.2 | 379 | Complete match |
| SREBF2 | FXC02129,FHC02129 | NM_004599.3 | 390 | Complete match |
| SRY | FXC03218,FHC03218,ORH03218 | NM_003140.2 | 699 | Complete match |
| ST18 | FXC00555,FHC00555,ORK00555 | NM_014682.2 | 397 | Complete match |
| STAG1 | FXC01192,FHC01192,ORK01192 | NM_005862.2 | 465 | Complete match |
| STAG2 | FXC11817,FHC11817,ORK11817,ORH11817P | NM_006603.4 | 538 | Complete match |

| | | | | |
|---|---|---|---|---|
| STAT1 | ORH13908P | NM_139266.2 | 392 | Complete match |
| STAT2 | FXC01316,FHC01316,ORK01316,ORH01316P | NM_005419.3 | 650 | Complete match |
| STAT3 | FXC01466,FHC01466 | NM_003150.3 | 420 | Complete match |
| STAT4 | FHC09368,ORS09368,ORH09368 | NM_003151.3 | 403 | Complete match |
| STAT5B | FXC01264,FHC01264,ORH01264 | NM_012448.3 | 433 | Complete match |
| STAT6 | FXC01445,FHC01445,ORH01445P | NM_003153.4 | 519 | Complete match |
| SUB1 | FXC01369,FHC01369 | NM_006713.3 | 723 | Complete match |
| SUFU | FHC08683,ORS08683,ORH08683 | NM_016169.3 | 355 | Complete match |
| SUPT4H1 | FXC03831,FHC03831,ORH03831P | NM_003168.2 | 363 | Complete match |
| TADA2A | FXC01576,FHC01576,ORK01576 | NM_001488.4 | 727 | Complete match |
| TADA2B | FHC31029 | NM_152293.2 | 437 | Complete match |
| TADA3 | FXC02633,FHC02633,ORK02633 | NM_133480.2 | 318 | Complete match |
| TAF10 | FHC24133 | NM_006284.3 | 298 | Complete match |
| TAF12 | FXC09981,FHC09981,ORS09981,ORH09981 | NM_005644.3 | 466 | Complete match |
| TAF13 | FXC05999,FHC05999,ORH05999P | NM_005645.3 | 310 | Complete match |
| TAF1B | FXC03763,FHC03763 | NM_005680.2 | 419 | Complete match |
| TAF2 | FHC07055,ORH07055 | NM_003184.3 | 388 | Complete match |
| TAF3 | FHC11143 | NM_031923.3 | 473 | Complete match |
| TAF4B | FXC03662,FHC03662 | NM_005640.2 | 543 | Complete match |
| TAF5 | FXC03285,FHC03285 | NM_006951.3 | 551 | Complete match |
| TAF5L | ORH14420P | NM_001025247.1 | 424 | Complete match |
| TAF6 | FXC08940,FHC08940,ORS08940,ORH08940 | NM_005641.3 | 652 | Complete match |
| TAF6L | FHC12542 | NM_006473.3 | 346 | Complete match |
| TAF7 | FXC01502,FHC01502,ORK01502,ORS01502,ORH01502 | NM_005642.2 | 273 | Complete match |
| TAL1 | FXC02210,FHC02210,ORK02210 | NM_003189.5 | 514 | Complete match |
| TAL2 | FXC03240,FHC03240,ORH03240P | NM_005421.2 | 453 | Complete match |
| TARDBP | FXC01302,FHC01302,ORK01302 | NM_007375.3 | 510 | Complete match |
| TAX1BP1 | FXC02883,FHC02883,ORH02883 | NM_006024.6 | 291 | Complete match |
| TBP | FHC11099 | NM_003194.5 | 371 | Partial match |
| TBPL2 | FHC12596 | NM_199047.2 | 457 | Complete match |
| TBR1 | FXC11886,FHC11886,ORH11886P | NM_006593.2 | 461 | Complete match |
| TBX10 | FHC28057 | NM_005995.4 | 572 | Complete match |
| TBX15 | ORH27685P | NM_152380.2 | 351 | Complete match |
| TBX18 | FHC12628,ORH12628P | NM_001080508.2 | 457 | Complete match |
| TBX19 | FHC20700,ORH20700 | NM_005149.2 | 392 | Complete match |
| TBX20 | FXC03772,FHC03772 | NM_001077653.2 | 593 | Complete match |
| TBX22 | FXC03910,FHC03910,ORH03910P | NM_016954.2 | 612 | Complete match |
| TBX3 | FHC10185,ORS10185,ORH10185 | NM_016569.3 | 732 | Complete match |
| TBX5 | FXC03447,FHC03447,ORH03447 | NM_000192.3 | 712 | Complete match |
| TBX6 | FXC07072,FHC07072 | NM_004608.3 | 863 | Complete match |
| TCEAL1 | FXC03223,FHC03223,ORS03223,ORH03223 | NM_004780.2 | 645 | Complete match |
| TCF12 | FXC01255,FHC01255,ORK01255,ORH01255 | NM_207036.1 | 695 | Complete match |

| | | | | |
|---|---|---|---|---|
| TCF15 | FHC28802 | NM_004609.3 | 290 | Complete match |
| TCF19 | FXC03667,FHC03667 | NM_001077511.1 | 519 | Complete match |
| TCF20 | FXC11595,FHC11595,ORK11595 | NM_005650.2 | 347 | Complete match |
| TCF21 | FXC03082,FHC03082,ORH03082 | NM_003206.3 | 490 | Complete match |
| TCF25 | FXC11958,FHC11958,ORH11958 | NM_014972.2 | 575 | Complete match |
| TCF3 | FXC01482,FHC01482 | NM_003200.3 | 366 | Complete match |
| TCF4 | FXC03619,FHC03619 | NM_001243226.2 | 564 | Complete match |
| TCF7 | FHC22279,ORH22279 | NM_003202.3 | 361 | Complete match |
| TCF7L1 | FHC21071 | NM_031283.2 | 469 | Complete match |
| TCF7L2 | FHC11660,ORH11660P | NM_001146284.1 | 596 | Complete match |
| TEAD1 | FXC11407,FHC11407 | NM_021961.6 | 830 | Partial match |
| TEAD2 | FXC07084,FHC07084,ORS07084,ORH07084 | NM_003598.1 | 388 | Complete match |
| TEAD3 | FHC40638M | NM_003214.3 | 756 | Complete match |
| TEAD4 | FXC11399,FHC11399 | NM_201443.2 | 285 | Complete match |
| TEF | FXC01581,FHC01581,ORK01581 | NM_001145398.2 | 343 | Complete match |
| TERF2IP | FXC02596,FHC02596,ORH02596P | NM_018975.3 | 289 | Complete match |
| TFAM | FXC02491,FHC02491,ORH02491P | NM_003201.2 | 539 | Complete match |
| TFAP2A | FXC08913,FHC08913,ORS08913,ORH08913 | NM_001032280.2 | 490 | Complete match |
| TFAP2B | FXC09636,FHC09636,ORS09636,ORH09636 | NM_003221.3 | 589 | Complete match |
| TFAP2C | FHC11683 | NM_003222.3 | 572 | Complete match |
| TFAP2D | FHC22652 | NM_172238.3 | 518 | Complete match |
| TFAP2E | FHC12602 | NM_178548.3 | 548 | Complete match |
| TFAP4 | FXC07094,FHC07094,ORH07094P | NM_003223.2 | 666 | Complete match |
| TFCP2 | FXC02799,FHC02799,ORK02799,ORH02799P | NM_005653.4 | 445 | Complete match |
| TFCP2L1 | FXC07633,FHC07633 | NM_014553.2 | 539 | Complete match |
| TFDP1 | FXC08570,FHC08570,ORS08570,ORH08570 | NM_007111.4 | 482 | Complete match |
| TFDP2 | FXC11405,FHC11405 | NM_006286.4 | 550 | Complete match |
| TFE3 | FXC10474,FHC10474,ORK10474 | NM_006521.5 | 945 | Complete match |
| TFEB | ORH13364P | NM_007162.2 | 441 | Complete match |
| TFEC | FHC23058 | NM_012252.3 | 457 | Complete match |
| TGFB1 | FHC28628,ORH28628P | NM_000660.5 | 425 | Complete match |
| TGIF1 | FXC03413,FHC03413 | NM_170695.3 | 543 | Complete match |
| TGIF2 | FXC03092,FHC03092,ORS03092,ORH03092 | NM_021809.6 | 879 | Complete match |
| THRA | FXC07745,FHC07745,ORS07745,ORH07745 | NM_003250.5 | 543 | Complete match |
| THRB | FXC01325,FHC01325,ORH01325 | NM_000461.4 | 424 | Complete match |
| TICAM1 | ORH15381P | NM_182919.3 | 287 | Partial match |
| TIRAP | FHC09421,ORS09421,ORH09421 | NM_001039661.1 | 469 | Partial match |
| TLX1 | FXC07127,FHC07127 | NM_005521.3 | 556 | Complete match |
| TNFAIP3 | FXC11936,FHC11936,ORH11936P | NM_006290.3 | 433 | Complete match |
| TNFRSF11A | FXC10751,FHC10751,ORH10751 | NM_003839.3 | 456 | Complete match |
| TNFRSF4 | FHC20008 | NM_003327.3 | 382 | Complete match |
| TNFSF11 | FXC24993,FHC24993,ORH24993P | NM_003701.3 | 576 | Complete match |

| | | | | |
|---|---|---|---|---|
| TNFSF18 | FHC14055,ORH14055P | NM_005092.3 | 363 | Partial match |
| TNFSF4 | FHC10802 | NM_003326.4 | 433 | Complete match |
| TP53 | FXC40948M | NM_001126118.1 | 416 | Complete match |
| TP53BP1 | FXC00998,FHC00998,ORK00998 | NM_001141979.1 | 346 | Complete match |
| TP63 | FXC21805M | NM_001114982.1 | 793 | Complete match |
| TP73 | FHC20035M | NM_001126241.2 | 371 | Complete match |
| TRAF1 | FHC10871,ORH10871P | NM_005658.4 | 367 | Complete match |
| TRAF2 | FXC09417,FHC09417,ORS09417,ORH09417 | NM_021138.3 | 416 | Complete match |
| TRAF3 | FXC03551,FHC03551 | NM_003300.3 | 486 | Complete match |
| TRAF5 | FHC20808 | NM_004619.3 | 400 | Complete match |
| TRAF6 | FXC01545,FHC01545,ORK01545 | NM_004620.3 | 391 | Complete match |
| TRERF1 | ORH12066 | NM_033502.3 | 555 | Complete match |
| TRIB1 | FXC01553,FHC01553,ORK01553,ORH01553P | NM_025195.3 | 248 | Complete match |
| TRIM13 | FXC11734,FHC11734,ORH11734P | NM_005798.4 | 646 | Complete match |
| TRIM14 | FXC11594,FHC11594 | NM_014788.3 | 522 | Complete match |
| TRIM15 | FXC02555,FHC02555,ORH02555P | NM_033229.2 | 518 | Complete match |
| TRIM21 | FXC03697,FHC03697,ORH03697 | NM_003141.3 | 416 | Complete match |
| TRIM22 | FXC02572,FHC02572 | NM_006074.4 | 412 | Complete match |
| TRIM25 | FXC01299,FHC01299 | NM_005082.4 | 519 | Complete match |
| TRIM26 | FHC22535 | NM_003449.5 | 465 | Partial match |
| TRIM27 | FXC08698,FHC08698,ORH08698 | NM_006510.4 | 469 | Complete match |
| TRIM29 | FXC07200,FHC07200,ORH07200 | NM_012101.3 | 388 | Complete match |
| TRIM31 | FXC03855,FHC03855 | NM_007028.3 | 375 | Complete match |
| TRIM32 | FXC03381,FHC03381,ORS03381,ORH03381 | NM_012210.3 | 495 | Complete match |
| TRIM37 | FHC07202,ORK07202,ORH07202P | NM_015294.3 | 409 | Complete match |
| TRIM38 | FXC03875,FHC03875 | NM_006355.4 | 395 | Complete match |
| TRIM40 | FXC03870,FHC03870,ORH03870 | NM_001286633.1 | 343 | Complete match |
| TRIM5 | FHC27937,ORH27937P | NM_033092.2 | 510 | Complete match |
| TRIM52 | FXC03821,FHC03821,ORH03821P | NM_032765.2 | 526 | Complete match |
| TRIM62 | FHC20199 | NM_018207.2 | 580 | Complete match |
| TRIM8 | FHC09071,ORS09071,ORH09071 | NM_030912.2 | 523 | Complete match |
| TRPS1 | FHC23365 | NM_014112.4 | 437 | Complete match |
| TSC22D2 | FXC00601,FHC00601,ORK00601 | NM_014779.3 | 393 | Complete match |
| TSC22D3 | FXC03403,FHC03403 | NM_198057.2 | 330 | Complete match |
| TSC22D4 | FXC07225,FHC07225,ORH07225P | NM_030935.4 | 342 | Complete match |
| TULP4 | FXC00825,FHC00825,ORK00825 | NM_001007466.2 | 523 | Complete match |
| TWIST2 | FXC09473,FHC09473,ORS09473,ORH09473 | NM_057179.2 | 588 | Complete match |
| UBB | FHC31552 | NM_018955.3 | 285 | Complete match |
| UBN1 | FXC01915,FHC01915,ORK01915 | NM_001079514.2 | 633 | Complete match |
| UFL1 | FXC00633,FHC00633,ORK00633 | NM_015323.4 | 600 | Complete match |
| UHRF1 | FHC12006,ORH12006 | NM_013282.4 | 367 | Complete match |
| USF2 | FXC07752,FHC07752 | NM_003367.2 | 432 | Complete match |
| VAV1 | FHC26428 | NM_005428.3 | 392 | Complete match |

| | | | | |
|---|---|---|---|---|
| VAX2 | FXC03879,FHC03879,ORH03879P | NM_012476.2 | 1218 | Complete match |
| VDR | FXC02558,FHC02558,ORH02558 | NM_000376.2 | 371 | Complete match |
| VSX1 | FHC26942,ORH26942 | NM_014588.5 | 454 | Complete match |
| WFS1 | FHC09316,ORH09316 | NM_006005.3 | 527 | Complete match |
| WT1 | ORH14336P | NM_001198551.1 | 348 | Complete match |
| WWP2 | FXC08703,FHC08703,ORS08703,ORH08703 | NM_007014.4 | 535 | Complete match |
| XBP1 | FXC03159,FHC03159,ORS03159,ORH03159 | NM_005080.3 | 604 | Complete match |
| XCL1 | FHC31349,ORH31349P | NM_002995.2 | 543 | Complete match |
| YBX1 | FXC01499,FHC01499,ORK01499,ORS01499,ORH01499 | NM_004559.3 | 282 | Complete match |
| YBX3 | FHC04649,ORS04649,ORH04649 | NM_003651.4 | 424 | Complete match |
| YEATS4 | FXC02697,FHC02697,ORK02697,ORH02697 | NM_006530.3 | 551 | Complete match |
| YY1 | FXC07394,FHC07394,ORS07394,ORH07394 | NM_003403.4 | 416 | Complete match |
| ZBTB11 | FHC11456 | NM_014415.3 | 461 | Complete match |
| ZBTB14 | FXC02729,FHC02729,ORH02729P | NM_003409.4 | 548 | Complete match |
| ZBTB16 | FXC03647,FHC03647,ORH03647 | NM_006006.4 | 2220 | Complete match |
| ZBTB17 | FXC03524,FHC03524 | NM_003443.2 | 494 | Complete match |
| ZBTB18 | FXC01424,FHC01424,ORH01424P | NM_006352.4 | 387 | Complete match |
| ZBTB20 | FXC02752,FHC02752,ORK02752,ORH02752 | NM_015642.5 | 506 | Complete match |
| ZBTB25 | FXC10716,FHC10716,ORH10716P | NM_006977.3 | 674 | Complete match |
| ZBTB38 | ORH14168P | NM_001080412.2 | 416 | Partial match |
| ZBTB4 | FXC00858,FHC00858,ORK00858,ORH00858P | NM_020899.3 | 482 | Complete match |
| ZBTB7A | FHC26401 | NM_015898.2 | 585 | Complete match |
| ZBTB7B | FXC02231,FHC02231,ORH02231P | NM_001256455.1 | 395 | Complete match |
| ZC3H12A | FHC20221,ORH20221P | NM_025079.2 | 269 | Complete match |
| ZC3H8 | FHC30900 | NM_032494.2 | 412 | Complete match |
| ZCCHC11 | ORH15286 | NM_001009881.2 | 425 | Complete match |
| ZEB1 | FXC01390,FHC01390 | NM_030751.5 | 326 | Partial match |
| ZFAT | FXC02036,FHC02036,ORK02036 | NM_020863.3 | 551 | Complete match |
| ZFHX3 | ORH13658P | NM_001164766.1 | 511 | Partial match |
| ZFP14 | FXC00863,FHC00863,ORK00863 | NM_020917.2 | 408 | Complete match |
| ZFP28 | FXC00831,FHC00831,ORK00831,ORH00831P | NM_020828.1 | 769 | Complete match |
| ZFP3 | FXC03197,FHC03197,ORH03197P | NM_153018.2 | 384 | Complete match |
| ZFP30 | FXC00698,FHC00698,ORK00698,ORH00698 | NM_014898.2 | 494 | Complete match |
| ZFP36L1 | FXC01454,FHC01454,ORS01454,ORH01454 | NM_004926.3 | 1015 | Complete match |
| ZFP36L2 | FHC11083 | NM_006887.4 | 717 | Complete match |
| ZFP37 | FHC23608,ORH23608P | NM_003408.2 | 395 | Complete match |
| ZFP42 | FXC03231,FHC03231 | NM_174900.4 | 465 | Complete match |
| ZFP69B | FXC10428,FHC10428,ORH10428P | NM_023070.3 | 522 | Partial match |
| ZFP82 | FXC00300,FHC00300,ORK00300 | NM_133466.2 | 330 | Complete match |
| ZFPM1 | FHC28405M | NM_153813.2 | 556 | Complete match |

| | | | | |
|---|---|---|---|---|
| ZFPM2 | FHC11749 | NM_012082.3 | 379 | Complete match |
| ZGLP1 | FHC30791 | NM_001103167.1 | 580 | Complete match |
| ZGPAT | FXC00286,FHC00286,ORK00286 | NM_181485.2 | 403 | Complete match |
| ZHX1 | FXC07776,FHC07776,ORK07776 | NM_007222.4 | 363 | Complete match |
| ZHX2 | FXC00659,FHC00659,ORK00659 | NM_014943.3 | 393 | Complete match |
| ZHX3 | FHC07426,ORK07426 | NM_015035.3 | 301 | Complete match |
| ZIC1 | FXC07427,FHC07427,ORH07427P | NM_003412.3 | 648 | Complete match |
| ZIC3 | FXC02318,FHC02318,ORH02318 | NM_003413.3 | 470 | Complete match |
| ZIK1 | ORH16369 | NM_001010879.4 | 338 | Partial match |
| ZIM2 | ORH28675P | NM_015363.4 | 412 | Complete match |
| ZIM3 | FHC28677,ORH28677 | NM_052882.1 | 385 | Complete match |
| ZKSCAN1 | FXC01039,FHC01039,ORK01039,ORH01039P | NM_003439.2 | 466 | Complete match |
| ZKSCAN2 | FHC12667,ORH12667P | NM_001012981.4 | 818 | Complete match |
| ZKSCAN3 | FHC11724 | NM_024493.3 | 310 | Complete match |
| ZKSCAN4 | FHC10011,ORS10011,ORH10011 | NM_019110.4 | 400 | Complete match |
| ZKSCAN5 | FHC11236,ORH11236P | NM_014569.3 | 367 | Complete match |
| ZKSCAN7 | FHC12625 | NM_018651.3 | 326 | Complete match |
| ZKSCAN8 | FHC12657,ORH12657 | NM_006298.3 | 367 | Complete match |
| ZMYM5 | ORH24953P | NM_001039649.2 | 580 | Complete match |
| ZNF112 | ORH26677 | NM_013380.3 | 564 | Complete match |
| ZNF12 | FHC31160 | NM_016265.3 | 583 | Complete match |
| ZNF121 | ORS28571,ORH28571 | NM_001008727.2 | 285 | Complete match |
| ZNF131 | FXC07439,FHC07439 | NM_001297548.1 | 634 | Complete match |
| ZNF132 | FHC11045,ORH11045P | NM_003433.3 | 469 | Complete match |
| ZNF133 | FXC01509,FHC01509,ORK01509 | NM_003434.5 | 494 | Complete match |
| ZNF134 | FXC10790,FHC10790,ORH10790P | NM_003435.3 | 380 | Complete match |
| ZNF135 | FXC02807,FHC02807 | NM_001289401.1 | 518 | Complete match |
| ZNF137P | ORH13070P | | 469 | Complete match |
| ZNF138 | FHC11129 | NM_006524.3 | 476 | Complete match |
| ZNF140 | FXC01586,FHC01586,ORH01586P | NM_003440.3 | 592 | Complete match |
| ZNF146 | FXC02723,FHC02723,ORK02723,ORH02723P | NM_007145.2 | 378 | Complete match |
| ZNF148 | FXC03621,FHC03621 | NM_021964.2 | 539 | Complete match |
| ZNF154 | FXC00968,FHC00968,ORK00968 | NM_001085384.2 | 302 | Complete match |
| ZNF155 | FHC26675,ORH26675 | NM_003445.3 | 498 | Complete match |
| ZNF157 | FXC03940,FHC03940 | NM_003446.3 | 530 | Complete match |
| ZNF165 | FHC22525,ORH22525 | NM_003447.3 | 346 | Complete match |
| ZNF174 | FXC03414,FHC03414 | NM_003450.2 | 425 | Complete match |
| ZNF175 | FXC03604,FHC03604,ORH03604P | NM_007147.2 | 449 | Complete match |
| ZNF18 | FXC07444,FHC07444,ORH07444P | NM_144680.3 | 502 | Complete match |
| ZNF182 | FXC11640,FHC11640,ORK11640 | NM_006962.1 | 641 | Complete match |
| ZNF189 | FHC11665,ORH11665P | NM_001278231.1 | 422 | Complete match |
| ZNF19 | FXC03978,FHC03978,ORH03978P | NM_006961.3 | 568 | Complete match |
| ZNF197 | FXC11837,FHC11837,ORK11837 | NM_006991.3 | 428 | Complete match |
| ZNF2 | FXC07446,FHC07446 | NM_021088.3 | 392 | Complete match |

| | | | | |
|---|---|---|---|---|
| ZNF202 | FXC00326,FHC00326,ORK00326,ORH00326P | NM_003455.3 | 334 | Complete match |
| ZNF205 | FXC07447,FHC07447,ORH07447P | NM_003456.2 | 379 | Complete match |
| ZNF211 | FHC26839 | NM_006385.3 | 458 | Complete match |
| ZNF213 | FXC07448,FHC07448,ORH07448P | NM_004220.2 | 523 | Complete match |
| ZNF215 | FHC24137 | NM_013250.2 | 485 | Complete match |
| ZNF217 | FHC31747 | NM_006526.2 | 919 | Complete match |
| ZNF224 | FXC01211,FHC01211,ORK01211,ORH01211P | NM_013398.2 | 575 | Complete match |
| ZNF227 | ORH26676P | NM_182490.2 | 445 | Complete match |
| ZNF23 | ORH25767P | NM_145911.2 | 347 | Complete match |
| ZNF232 | ORH11970 | NM_014519.2 | 339 | Complete match |
| ZNF239 | FXC03229,FHC03229 | NM_005674.2 | 477 | Complete match |
| ZNF24 | FXC11380,FHC11380,ORH11380P | NM_006965.2 | 412 | Complete match |
| ZNF250 | FXC01237,FHC01237 | NM_001109689.3 | 384 | Complete match |
| ZNF254 | ORH28603P | NM_203282.3 | 474 | Complete match |
| ZNF256 | FXC07450,FHC07450 | NM_005773.2 | 462 | Complete match |
| ZNF26 | ORH28195P | NM_019591.3 | 445 | Complete match |
| ZNF260 | ORH16238P | NM_001012756.2 | 449 | Complete match |
| ZNF263 | FXC11798,FHC11798,ORK11798,ORH11798P | NM_005741.4 | 530 | Complete match |
| ZNF267 | ORH28385P | NM_003414.5 | 457 | Complete match |
| ZNF268 | FHC30561,ORH30561P | NM_003415.2 | 449 | Complete match |
| ZNF274 | FXC03631,FHC03631,ORH03631P | NM_133502.2 | 469 | Complete match |
| ZNF280A | FXC07758,FHC07758,ORH07758P | NM_080740.4 | 462 | Complete match |
| ZNF280B | FXC03154,FHC03154,ORS03154,ORH03154 | NM_080764.3 | 354 | Complete match |
| ZNF280C | ORH27521 | NM_017666.4 | 396 | Complete match |
| ZNF280D | FXC00250,FHC00250,ORK00250 | NM_001002843.2 | 621 | Complete match |
| ZNF281 | FXC01275,FHC01275,ORH01275P | NM_012482.4 | 605 | Complete match |
| ZNF286A | FXC00939,FHC00939,ORK00939 | NM_020652.2 | 400 | Complete match |
| ZNF292 | FXC07771,ORK07771 | NM_015021.3 | 514 | Partial match |
| ZNF3 | FXC02646,FHC02646,ORK02646 | NM_032924.4 | 445 | Complete match |
| ZNF30 | ORH16381 | NM_194325.2 | 347 | Complete match |
| ZNF300 | FXC07455,FHC07455,ORH07455P | NM_052860.2 | 445 | Complete match |
| ZNF302 | FXC07654,FHC07654,ORH07654P | NM_018443.3 | 591 | Complete match |
| ZNF304 | FXC03634,FHC03634 | NM_020657.3 | 576 | Complete match |
| ZNF311 | ORH29129 | NM_001010877.2 | 331 | Complete match |
| ZNF317 | FHC07457,ORK07457,ORH07457P | NM_020933.4 | 437 | Complete match |
| ZNF320 | ORH28665 | NM_207333.2 | 343 | Complete match |
| ZNF322 | FXC01932,FHC01932,ORH01932 | NM_024639.4 | 359 | Complete match |
| ZNF334 | ORH28817 | NM_018102.4 | 440 | Complete match |
| ZNF33A | FXC00392,FHC00392,ORK00392 | NM_006974.2 | 532 | Complete match |
| ZNF345 | FXC11735,FHC11735,ORH11735P | NM_003419.4 | 506 | Complete match |
| ZNF35 | FXC03941,FHC03941,ORH03941P | NM_003420.3 | 527 | Complete match |
| ZNF350 | FXC07462,FHC07462,ORH07462P | NM_021632.3 | 458 | Complete match |
| ZNF354A | FXC07463,FHC07463,ORH07463P | NM_005649.2 | 597 | Complete match |

| | | | | |
|---|---|---|---|---|
| ZNF354B | FXC07464,FHC07464 | NM_058230.2 | 404 | Complete match |
| ZNF354C | ORH16279P | NM_014594.2 | 515 | Partial match |
| ZNF367 | FHC23565 | NM_153695.3 | 449 | Complete match |
| ZNF37A | FXC03448,FHC03448,ORH03448P | NM_003421.2 | 634 | Complete match |
| ZNF382 | ORS28615 | NM_032825.4 | 355 | Complete match |
| ZNF383 | FXC10734,FHC10734,ORH10734 | NM_152604.1 | 517 | Complete match |
| ZNF396 | FHC26290,ORH26290P | NM_145756.2 | 407 | Complete match |
| ZNF397 | FHC28540,ORH28540 | NM_032347.2 | 320 | Complete match |
| ZNF41 | FHC11105,ORH11105P | NM_007130.2 | 437 | Complete match |
| ZNF415 | FHC10812 | NM_001352138.1 | 423 | Partial match |
| ZNF418 | FXC00954,FHC00954,ORK00954,ORH00954P | NM_133460.1 | 441 | Complete match |
| ZNF420 | ORH26603 | NM_144689.3 | 363 | Complete match |
| ZNF43 | FXC07470,FHC07470,ORH07470 | NM_003423.3 | 437 | Complete match |
| ZNF431 | FXC00960,FHC00960,ORK00960,ORH00960P | NM_133473.2 | 377 | Complete match |
| ZNF432 | FXC00643,FHC00643,ORK00643 | NM_014650.2 | 453 | Complete match |
| ZNF436 | FXC01633,FHC01633,ORK01633,ORH01633 | NM_030634.2 | 416 | Complete match |
| ZNF438 | ORH23807 | NM_182755.2 | 347 | Complete match |
| ZNF444 | FXC03394,FHC03394 | NM_001253792.1 | 441 | Complete match |
| ZNF445 | FXC01855,FHC01855,ORK01855 | NM_181489.5 | 473 | Complete match |
| ZNF446 | FXC07475,FHC07475 | NM_017908.3 | 363 | Complete match |
| ZNF45 | FXC03614,FHC03614 | NM_003425.3 | 482 | Complete match |
| ZNF451 | FXC00565,FHC00565,ORK00565 | NM_001031623.2 | 371 | Complete match |
| ZNF454 | FXC07478,FHC07478,ORH07478P | NM_182594.2 | 351 | Complete match |
| ZNF461 | FXC01368,FHC01368,ORH01368P | NM_153257.5 | 359 | Partial match |
| ZNF480 | ORS08973,ORH08973 | NM_144684.4 | 469 | Complete match |
| ZNF483 | FXC00958,FHC00958,ORK00958 | NM_133464.3 | 432 | Complete match |
| ZNF484 | ORH29383P | NM_031486.2 | 559 | Complete match |
| ZNF490 | FXC00199,FHC00199,ORK00199,ORH00199P | NM_020714.2 | 424 | Complete match |
| ZNF493 | FXC07480,FHC07480,ORK07480 | NM_175910.6 | 548 | Complete match |
| ZNF496 | ORS09908,ORH09908 | NM_032752.1 | 339 | Complete match |
| ZNF500 | FXC00562,FHC00562,ORK00562 | NM_021646.2 | 347 | Complete match |
| ZNF501 | FXC03018,FHC03018 | NM_145044.3 | 273 | Partial match |
| ZNF514 | FXC02976,FHC02976,ORK02976 | NM_032788.1 | 470 | Complete match |
| ZNF516 | FXC00467,FHC00467,ORK00467 | NM_014643.3 | 453 | Complete match |
| ZNF528 | FHC12005,ORH12005 | NM_032423.2 | 318 | Complete match |
| ZNF540 | ORH26607 | NM_152606.4 | 384 | Complete match |
| ZNF542P | FXC03194,FHC03194,ORK03194,ORH03194P | NR_033418.1 | 469 | Partial match |
| ZNF546 | ORH26630P | NM_178544.4 | 592 | Complete match |
| ZNF548 | ORH30863P | NM_152909.3 | 662 | Complete match |
| ZNF549 | FXC06055,FHC06055,ORH06055 | NM_001199295.1 | 579 | Complete match |
| ZNF552 | FXC11245,FHC11245 | NM_024762.3 | 485 | Complete match |
| ZNF571 | FXC02768,FHC02768,ORH02768 | NM_016536.4 | 457 | Complete match |
| ZNF573 | ORH11982 | NM_001172690.1 | 461 | Partial match |

| | | | | |
|---|---|---|---|---|
| ZNF577 | ORH26782 | NM_032679.2 | 413 | Partial match |
| ZNF583 | FXC02620,FHC02620,ORH02620 | NM_152478.2 | 314 | Complete match |
| ZNF584 | ORH26848 | NM_173548.1 | 359 | Complete match |
| ZNF585B | ORH26604 | NM_152279.3 | 441 | Complete match |
| ZNF586 | FXC03846,FHC03846 | NM_017652.3 | 535 | Complete match |
| ZNF606 | FXC00932,FHC00932,ORK00932,ORH00932P | NM_025027.3 | 355 | Complete match |
| ZNF607 | ORH14071P | NM_032689.5 | 358 | Partial match |
| ZNF621 | ORH12003 | NM_198484.4 | 531 | Complete match |
| ZNF623 | FXC01987,FHC01987,ORK01987,ORH01987 | NM_014789.3 | 393 | Complete match |
| ZNF629 | FXC00506,FHC00506,ORK00506 | NM_001080417.1 | 568 | Complete match |
| ZNF639 | FHC21765,ORH21765 | NM_016331.2 | 363 | Complete match |
| ZNF641 | FXC11384,FHC11384 | NM_152320.2 | 363 | Complete match |
| ZNF649 | ORH04863 | NM_023074.3 | 408 | Complete match |
| ZNF655 | ORH22999 | NM_138494.2 | 372 | Complete match |
| ZNF660 | FXC00417,FHC00417,ORH00417P | NM_173658.2 | 408 | Complete match |
| ZNF668 | FXC01451,FHC01451,ORK01451 | NM_024706.4 | 326 | Complete match |
| ZNF677 | ORH26790 | NM_182609.2 | 367 | Complete match |
| ZNF691 | FXC03193,FHC03193,ORH03193 | NM_015911.3 | 353 | Complete match |
| ZNF699 | ORH29897 | NM_198535.1 | 347 | Complete match |
| ZNF7 | FHC23437,ORH23437 | NM_003416.3 | 896 | Complete match |
| ZNF71 | FXC11736,FHC11736,ORH11736P | NM_021216.4 | 367 | Complete match |
| ZNF726 | ORH15923P | NM_001244038.1 | 327 | Complete match |
| ZNF75D | FXC03758,FHC03758,ORH03758 | NM_007131.4 | 523 | Complete match |
| ZNF761 | FXC07722,FHC07722,ORK07722 | NM_001008401.3 | 511 | Complete match |
| ZNF773 | ORH11964 | NM_198542.2 | 367 | Complete match |
| ZNF780A | ORS28623 | NM_001010880.2 | 388 | Complete match |
| ZNF792 | ORH13233P | NM_175872.5 | 343 | Complete match |
| ZNF8 | FXC01460,FHC01460,ORK01460,ORH01460P | NM_021089.2 | 383 | Complete match |
| ZNF808 | ORH13088P | | 392 | Complete match |
| ZNF81 | FXC03282,FHC03282 | NM_007137.3 | 570 | Complete match |
| ZNF816 | ORH28666 | NM_001031665.2 | 428 | Complete match |
| ZNF821 | FXC03694,FHC03694 | NM_001201552.1 | 441 | Complete match |
| ZNF829 | FXC11264,FHC11264,ORH11264P | NM_001037232.3 | 424 | Complete match |
| ZNF83 | FXC03233,FHC03233 | NM_018300.3 | 465 | Complete match |
| ZNF837 | ORH16292P | NM_138466.1 | 335 | Complete match |
| ZNF85 | FXC10669,FHC10669 | NM_003429.4 | 428 | Complete match |
| ZNF880 | ORH12329P | NM_001145434.2 | 295 | Partial match |
| ZNF90 | FXC03247,FHC03247 | NM_007138.2 | 601 | Complete match |
| ZNF92 | FHC29232 | NM_007139.3 | 416 | Complete match |
| ZRANB2 | FHC07627,ORH07627P | NM_005455.4 | 289 | Complete match |
| ZSCAN1 | FHC13090,ORH13090P | NM_182572.3 | 658 | Partial match |
| ZSCAN12 | FXC00534,FHC00534,ORK00534 | NM_001163391.1 | 330 | Partial match |
| ZSCAN16 | FXC07525,FHC07525,ORH07525 | NM_025231.1 | 505 | Complete match |
| ZSCAN18 | FXC02603,FHC02603,ORK02603 | NM_023926.4 | 314 | Complete match |

| | | | | |
|---|---|---|---|---|
| ZSCAN2 | FHC25490,ORH25490P | NM_017894.5 | 285 | Complete match |
| ZSCAN20 | FHC15514,ORH15514P | NM_145238.3 | 320 | Partial match |
| ZSCAN21 | FXC02732,FHC02732,ORH02732 | NM_145914.2 | 392 | Complete match |
| ZSCAN22 | FXC07527,FHC07527,ORH07527 | NM_181846.2 | 310 | Complete match |
| ZSCAN26 | FXC02200,FHC02200,ORK02200 | NM_001023560.3 | 456 | Complete match |
| ZSCAN29 | FXC03798,FHC03798 | NM_152455.3 | 494 | Complete match |
| ZSCAN30 | FHC29873 | NM_001112734.3 | 383 | Complete match |
| ZSCAN32 | ORH12804P | NM_001284527.2 | 539 | Partial match |
| ZSCAN4 | FHC26841,ORH26841 | NM_152677.2 | 428 | Complete match |
| ZSCAN5A | FXC07529,FHC07529,ORS07529,ORH07529 | NM_024303.1 | 363 | Complete match |
| ZSCAN5B | FHC12611 | NM_001080456.2 | 583 | Complete match |
| ZSCAN9 | FXC06024,FHC06024,ORH06024 | NM_006299.4 | 486 | Complete match |
| ZXDA | FHC27411,ORH27411 | NM_007156.4 | 514 | Complete match |
| ZXDC | ORH30087 | NM_025112.5 | 260 | Partial match |
